# Kinetic organization of the genome revealed by ultra-resolution, multiscale live imaging

**DOI:** 10.1101/2025.03.27.645817

**Authors:** Joo Lee, Liang-Fu Chen, Simon Gaudin, Kavvya Gupta, Andrew Spakowitz, Alistair Nicol Boettiger

## Abstract

In the last decade, sequencing methods like Hi-C have made it clear the genome is intricately folded, and that this organization contributes significantly to the control of gene expression and thence cell fate and behavior. Single-cell DNA tracing microscopy and polymer physics-based simulations of genome folding have proposed these population-scale patterns arise from motor-driven, heterogeneous movement, rather than stable 3D genomic architecture, implying that motion, rather than structure, is key to understanding genome function. However, tools to directly observe this motion *in vivo* have been limited in coverage and resolution. Here we describe TRansposon Assisted Chromatin Kinetic Imaging Technology (TRACK-IT), which combines a suite of imaging and labeling improvements to achieve ultra-resolution in space and time, with self-mapping transposons to distribute labels across the chromosome, uncovering dynamic behaviors across four orders of magnitude of genomic separation. We find that sequences separated by sub-megabase distances, typically 200-500 nm of nanometers apart, can transition to close proximity in tens of seconds - faster than previously hypothesized. This rapid motion is dependent upon cohesin and is exhibited only within certain genomic domains. Domain borders act as kinetic impediments to this search process, substantially slowing the rate and frequency of the transition to proximity. The genomic separation-dependent scaling of the search time for cis-interactions within a domain violates predictions of diffusion, suggesting motor driven folding. This distinctive scaling is lost following cohesin depletion, replaced with a behavior consistent with diffusion. Finally, we found cohesin containing cells exhibited rare, processive movements, not seen in cohesin depleted cells. These processive trajectories exhibit extrusion rates of ∼2.7 kb/s across three distinct genomic intervals, faster than recent *in vitro* measurements and prior estimates from *in vivo* data. Taken together, these results reveal a genome in motion across multiple genomic and temporal scales, where motor-dependent extrusion divides the sequence, not into spatially separate domains, but into kinetically separated domains that experience accelerated local search.

## Introduction

The mammalian genome is spatially organized in the nucleus, with complex, sequence-specific and cell-type-specific, contact patterns visible at all scales from kilobases to chromosomes. These contact patterns have been extensively mapped by Hi-C sequencing as pairwise contact-frequency heatmaps. Importantly, many recurring features of these heatmaps, such as dots, triangles, stripes, and checkerboards, correlate with functional maps of enhancer activity, epigenetic state, DNA replication and DNA repair - suggesting these spatial patterns play a role in major genome functions (*1*, *2*). Furthermore, perturbations that affect 3D genome organization impact these critical genome functions (*1*, *2*). Recent single-cell DNA-tracing microscopy has shown that these characteristic features from contact-frequency heatmaps are not *structures* that exist in folded genomes of individual cells, but *statistical patterns* that arise from averaging of many diverse folds from the many cells of the population (*1*). This confirmed the prediction from polymer modeling work (*3–6*), which suggested the patterns in contact heatmaps arise from statistical averaging (blurring) of a dynamic process in a single cell, and thus that genome *motion* regulates genomic functions like transcription and repair. However, little is known about the speed and timescales of genome motion as a function of genome sequence, genomic separation, and chromatin state.

The statistical patterns of genome organization are believed to be shaped by three major mechanisms; the diffusive behavior of the large chromosome polymers in a densely packed nucleus, the microphase-separation of the hetero/euchromatic components of the chromosome, and the extrusion of chromatin loops by the cohesin complex (*7*, *8*). The diffusion of long polymers that are packed in an unknotted “crumpled” (or “fractal”) globule is believed to explain the approximately inverse scaling of average contact-frequency as a function of genomic separation, first uncovered by Hi-C (*9*, *10*). Preferential adhesive interactions among similar chromatin types are believed to underlie the checkerboard patterns of contacts, called compartments (*9*, *11–14*). The extrusion of chromatin loops by cohesin, averaging >100 kb, is believed to increase local contact frequency within domains delineated by CTCF (called topologically associating domains, TADs). CTCF bound at the borders of TADs prevents cohesin loop-extrusion from pulling together elements of neighboring TADs, and thus leads to both triangles at the diagonal and CTCF-dots (*4*, *15*, *16*). This loop extrusion model gave an intuitive prediction for the effects of either cohesin or CTCF removal, later validated by experiment (*17–19*), and predicts some single cell-properties such as three-way contact frequencies also validated by experiments (*20–22*). Some groups have observed that these population-level TAD, their heterogeneous single-cell origins, and their dependence on cohesin and CTCF can also be reproduced by models of phase-separating “strings and binders” (*5*, *23*) or diffusing slip-links (*24*), rather than loop extrusion.

However, while the contribution of these different mechanisms to the steady-state, population-averaged features has been analyzed with simulations, Hi-C sequencing, and microscopy, our understanding of the dynamics and the timescale of genome motion as a function of sequence, distance, and chromatin state is just beginning. Recent experiments have shown direct evidence for loop extrusion *in vitro*, finding that cohesin can pull together DNA sequences separated by shear-flow, at a rate of ∼1 kb/s (*25*, *26*). This provides groundbreaking insight into understanding drivers of movement, but how these dynamics are modulated by *in vivo* context remains unexplored. Meanwhile pioneering live-cell imaging with pairs of labeled chromatin sites demonstrated substantial dynamic behaviors of these loci (*27*, *28*). Subsequent work on additional loci combined labeling and protein depletion experiments to show that both CTCF and cohesin restrict the mobility of the labeled sites (*29*, *30*), providing the first direct evidence for the dynamic explanation of the CTCF-dots, a much discussed statistical feature first observed by Hi-C.

Further progress from live imaging has been limited by the available tools, particularly in available temporal resolution, spatial resolution, and the ability to rapidly label many different genome regions. Indeed, despite extensive technology development for live-cell imaging (*31*–*54*), to our knowledge, less than a dozen cis-pairs have been labeled and quantitatively tracked in mammalian cells to date (*27*, *29*, *30*, *55*, *56*), covering only a small fraction of length-scales and features uncovered from fixed-cell approaches. Where fixed-cell data covers over five orders of magnitude of differences in contact frequency across a similar range of genomic lengths and a diverse set of chromatin states, available live-cell data rarely exceed two orders of magnitude in time and largely focus on a single cis-interaction per study (**fig. S1**). Most of these single loci studies have focused particularly on CTCF-anchored TAD boundaries to the exclusion of other genomic states. The limited depth in the time domain stems from limits in temporal resolution of recent approaches (primarily 20-30s) and data handling challenges, whereas the focus on single loci, rather than comparing across genomic contexts, stems largely from the difficulty in construction and validation of live-cell fluorescent labels. These recent works also illustrate the challenges arising from current limitations of spatial resolution, as the reported measurement uncertainties of ∼200 nm are sizable compared to the 100-600 nm range of 3D distances explored by the loci (*29*, *30*, *57*), making it challenging to separate movement from measurement error. This challenge has necessitated added assumptions and complex statistical inference in recent work in order to estimate kinetic properties such as contact duration (*29*, *30*) (see (*58*), for discussion).

Here we introduce TRansposon Assisted Chromatin Kinetic Imaging Technology (TRACK-IT), which allows us to measure chromatin motion in ultra-resolution in space and time, with distributed labels across a single chromosome. TRACK-IT utilizes transposon remobilization and *in situ* transcript sequencing, creating and precisely mapping hundreds of cell lines across diverse chromatin states and genomic separations with live-compatible fluorescent labels. We introduce optimized fluorescent labels that are brighter, more photostable, and miniaturized to better approximate a point-source and improve spatial resolution, compared to those currently used for DNA tracking. We analyze genome motion across genomic scales from a few kilobases to whole chromosomes, with sub-second resolution for 30 min of imaging and spatial precision on par with our prior fixed-cell super resolution work (<50 nm). We observe rapid motion at sub-megabase scales, transitioning from average distance to close proximity (<50 nm) in less than 20s. We find this fast search within TADs is cohesin-dependent and is lost across TAD borders. Cohesin not only accelerates intra-TAD search, but it mitigates much of the effect of genomic separation on search times, such that contacts between elements separated by a few hundred kilobases takes only marginally longer to form than contacts between elements a few tens of kilobases. Once cohesin was removed, we saw the effect of genomic separation on search become pronounced and consistent with analytic theory for diffusive space-filling polymers, rapidly scaling with genomic separation resulting in estimated multi-day search across megabases. Finally, we detect long-range processive motion specifically in cohesin-containing cells, which constrains an estimate of the *in vivo* extrusion speed of cohesin. Together, these results provide a quantitative characterization of genome motion, from the scales of a few kilobases to whole chromosomes, and reveal a genome that is not spatially partitioned in single cells, but rather kinetically divided into domains of cohesin-accelerated local search.

## Results

### High-throughput, ultra-resolution 4D genome imaging

To survey genome motion across a range of genomic length scales and sequence contexts, we combined Cas9-mediated genome editing, sleeping beauty transposition, and ultra-resolution, live-cell, fluorescent-labeling technology. We refer to this approach as TRansposon Assisted Chromatin Kinetic Imaging Technology (TRACK-IT) (**Fig. 1A**). The TRACK-IT cargo, with a pair of optimized TetO and CuO arrays juxtaposed, is integrated into a genomic region of interest with Cas9. Only the CuO array is contained within a Sleeping Beauty transposon, which upon mobilization, re-integrates across the genome, with a preference for proximal *cis* sites (*60*) (**fig. S2B-C**). We chose the Sleeping Beauty transposon for the neutral re- integration profile it exhibited (**fig. S2A-B**), contrasting the widely used PiggyBac transposon’s predisposition to reinsert into actively transcribed, open chromatin (*61*). We included a T7 promoter in an ‘outward’ direction within the transposon, allowing these new insertions to be self-mapping; by *in situ* transcription of the insertion sites followed by shallow-read sequencing to identify ITR-genome junctions (**Fig. 1B and S3**). We integrated the TRACK-IT construct in mouse embryonic stem cells, at a region of chromosome 6 that we previously studied using ORCA (**Fig. 1D**), which has a variety of loop-dots, TADs, genes and distinct chromatin states, reflective of the complexity typical of the mammalian genome (*20*). A single TRACK-IT experiment (3e5 cells transfected with Sleeping Beauty transposase) resulted in over 750 mobilization events across the genome (**fig. S2A-B**). Integrations covered a broad range of chromatin states (**Fig. 1C and S2D-E**) and provided dense coverage near the original insertion site (∼130 integrations within a 2 Mb window), and sparser coverage in distal regions and in *trans* (**fig. S2C**). Considering the small number of published cell lines, these data indicated TRACK-IT can rapidly extend the number of lines available to survey the effect of genomic context and separation (**Fig. 1C**). To screen the desired integration events, we developed an automated, high-throughput microscopy approach to screen individual clones in 96-well plates (**Fig. 1B**). This allowed microscopy validation of the different transposition events, and enabled image-based selection of proximal and distal cis-integration events, prior to sequence-mapping (**Fig. 1B**). Of the 750 insertion pairs, we selected 11 clones at logarithmically distributed genomic separations for further analyses (**Fig. 1D**). These clones spanned a variety of 3D genome features and distinct chromatin states (**Fig. 1D and S2D**); from tightly linked loci, 5 kb apart, (center-to-center separation between each labels) in a similar chromatin environment, to elements crossing sub-TAD and TAD borders (∼100s to ∼1000 kb), to those spanning multiple LADs, fully 73 Mb apart; collectively enabling investigation of genome behaviors across scales (**Fig. 1D-E**).

**Fig. 1.**
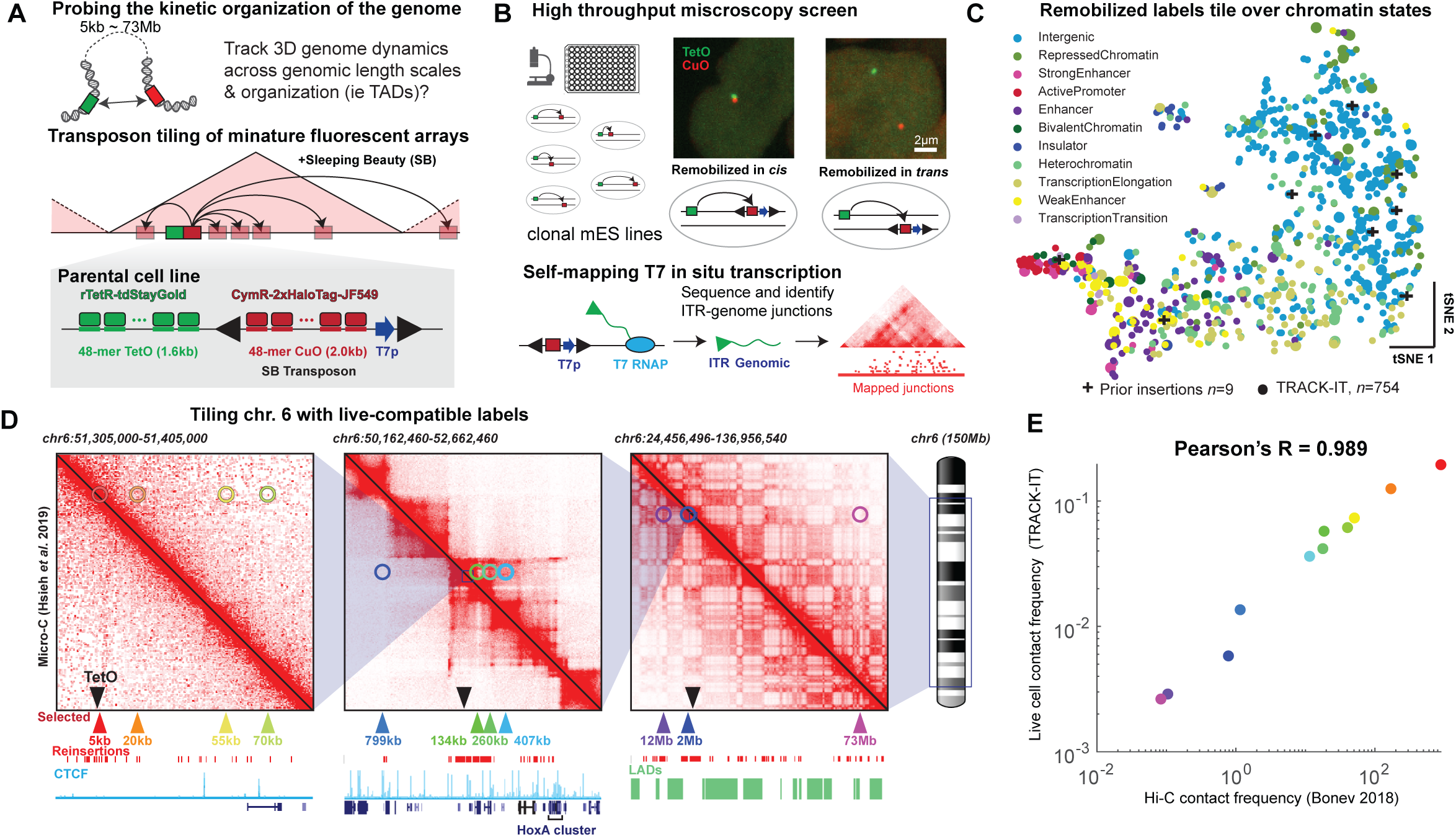
TRansposon Assisted Chromatin Kinetic Imaging Technology. **(A)** Schematic of fluorescent reporters and transposon based tiling. **(B)** Self-mapping with T7p transcripts. Example of cis (left panel) and trans (right panel) clones identified by automated microscopy. The precise position of each selected clone is mapped by *in situ transcription* using the T7 promoter placed adjacent to the transposing CuO tag followed by RNA-seq of these transcripts. **(C)** tSNE embedding of the diverse epigenetic states at the CuO integration sites after transposition, based on ChIP-seq data (see **fig. S2**). Each dot is a uniquely mapped cell line, the color indicates the ChromHMM state (*59*). The data from 754 lines made by TRACK-IT (larger marker sizes are within 1Mb from insertion sites) are overlaid on 9 mESC cell lines labeled for live-cell tracking in prior publications. **(D)** Genomic positions of 11 cell lines selected for further study. The top panels show micro-C data for spatial context and the bottom tracks show CTCF, gene-models, and LADs. The remaining integrations are shown as red ticks below the selected integrations (colored triangles), which are labeled based on their genomic separation from the TetO tag (inverted black triangle). **(E)** Correlation of the contact frequency measured from Live-cell imaging with TRACK-IT, and from Hi-C data from mESC (Bonev. 2017), using a 50 nm contact threshold for the TRACK-IT data.

We validated these TRACK-IT labels and measurements by comparing the contact frequency measured by live imaging of these 11 lines to prior data from Hi-C (*62*) (**Fig. 1E**). We used a 50 nm proximity threshold to define contact. Results from alternative thresholds from 30 nm to 500 nm are shown in **fig. S4D**. The Hi-C and TRACK-IT data correlated quantitatively, (Pearson’s *R*=0.98), close to agreement between the replicate TRACK-IT experiments (Pearson’s *R*=0.99) (**fig. S4A-B**). This provides a direct cross-validation of Hi-C contact frequencies with live-cell contact frequencies from distance measurements. For further validation, we compared the 3D distances (in nm) measured with TRACK-IT to previous measurements using ORCA in fixed cells, lacking TRACK-IT labels (**fig. S4C**), which also showed quantitative agreement (Pearson’s *R*=0.98), indicating that the live-cell labels did not substantially perturb genome structure. These live-cell experiments thus support, and are supported by, alternate measurements for investigating 3D genome organization with orthogonal limitations of fixation, hybridization, and/or ligation.

The quality of time series data is constrained by both the spatial and temporal resolution of the data, and can be characterized by the ratio of the amount of movement seen between consecutive frames, *F*, relative to the dynamic range of movement over the entire observation window, *D* (**Fig. 2A**). If the tracking speed is slow relative to the motion of the object, it will appear to jump discontinuously. If the measurement error in the 3D position is large, once again it will appear to jump discontinuously, even if tracking speed is fast. A continuous pattern arises only when both temporal and spatial resolution are high, resulting in a small ratio of the frame-to-frame motion (*F*) relative to the interquartile range of the entire movie (*D*), indicating a high quality time series (**Fig. 2A**). In order to optimize the quality of the TRACK-IT time series data (e.g. **Fig. 2C**), we made modifications to earlier state-of-the-art approaches to further improve the spatial and temporal resolution, including optimized fluorescent labels, miniaturized arrays, and improvements in optical imaging (**Fig. 2B**, and **Methods**). We selected a newly discovered, shockingly photostable, green fluorescent protein from *Cytaeis uchidae*, called StayGold (*63*), and screened a series of rationally designed linker variants between two tandem copies of StayGold to optimize expression in mouse embryonic stem cells (mESCs) (see **Methods** - 1. Optimization of fluorescence labels for DNA labeling). We paired this with Halotag-conjugating JF549 (*64–66*), a cell permeable organic dye brighter and more photostable than the genetically encoded fluorophores (*65*, *67*, *68*) used in prior work. We integrated both the rTetR-tdStayGold and CymR-2xHaloTag in hybrid (CAST/129) mESCs, and screened for clones with optimal expression levels to maximize signal-to-background and minimize cell-cell variability in expression levels (see **Methods**). This pair showed minimal loss of detected foci even after 3,600 z-stacks (**Fig. 2J**), imaged 0.5 s/stack (**Fig. 2H**).

**Fig. 2.**
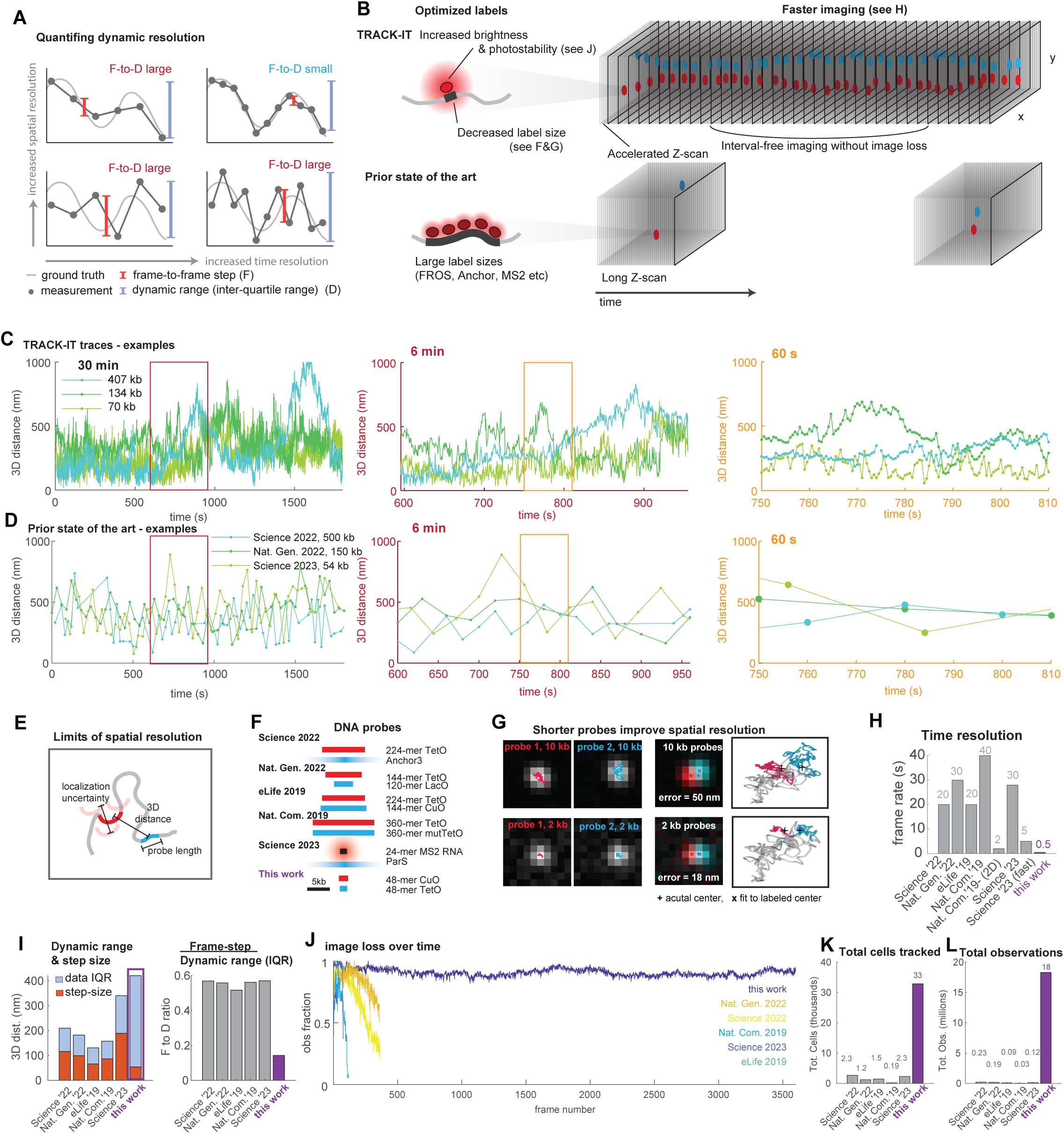
**Ultra-resolution live genome imaging**. **(A)** Schematic explanation of the frame-step vs. dynamic range. Small F-to-D ratio arises from better tracking of the underlying dynamics and requires improvements in both the spatial and time resolution. **(B)** Strategies adopted in this work to improve spatial and temporal resolution. (See also **Methods** - 1. Optimization of fluorescence labels for DNA labeling) **(C)** Three example traces from TRACK-IT data, zoom-in regions expand the time resolution. Traces are color-coded by genomic separation, as in **Fig. 1D**. **(D)** Example traces from three recent publications with similar genomic separations as in (C), chosen from the top decile of coverage. Note the x and y axes are the same as in (C). **(E)** Schematic illustration of the effect of label size on center-to-center 3D distance measurements. **(F)** Label size, benchmarked to prior state-of-the-art. **(G)** Simulations of the effect of label size on the measurement error for the label center. Polymer lengths were calibrated by ORCA, the point-spread function and camera binning were chosen to reflect ideal instrumentation. **(H)** Benchmarking reported time resolution. **I.** F-to-D ratio, dynamic range (blue) vs frame-to-frame step size (orange) and as a ratio (right). Note dynamic range is defined from the interquartile range of the data. **(J)** Fraction of traces with detected events per frame, a proxy for photostability. **(K)** Left, total number of cells analyzed in recent studies compared to this data set. **(L)** total number of observations per publication. Corresponding values for each dataset in each publication for the statistics in (J) and (K) are shown in **fig. S1B**.

The increased brightness and stability of these fluorophores allowed them to work well with fewer binding sites, contributing to improved spatial resolution. Prior work has used ∼200 copies or more of operator sites (**Fig. 2B, F**, and **S1A**), this moderate length of chromatin (10 kb) is a flexible polymer often ∼100-250 nm long (from fixed-cell measurements with ORCA (*58*, *69*, *70*)), that further limits spatial resolution (**Fig. 2E, G**). Alternative fluorescent labels based on the ParS/ParB system also suffer from the uncertainty in the label size of ParB foci (which assemble by stochastic spreading up to ∼20 kb (*71*, *72*), **Fig. 2F**). We used a substantially smaller repeat length (48-mer, 1.6 kb and 2.0 kb for TetO and CuO array, respectively), and also introduced random, heterogeneous spacers in between the binding sites (**fig. S1A** and **Methods**) - facilitating cloning and reducing the potential for repeat-mediated heterochromatin, as described in a recent report (*73*). We optimized a custom microscope setup for imaging these probes, using an accelerated *z*-scanning method, a microlens dual-spinning disk to minimize light loss and laser power, and dual, back-illuminated 95% quantum efficiency scientific CMOS cameras (see **Methods**). Residual 3D chromatic offsets were measured by imaging nucleoplasmic refractive index-matched 3D gel-embedded beads prior to every experiment, providing an alignment precision of 13 nm in *x-y* and 23 nm in *z* (**fig. S5**). With these multifaceted improvements, the TRACK-IT data showed a median 3-D frame-to-frame displacement of ∼50 nm across all 11 chromosomal length scales measured. This provides an upper bound to the spatial resolution, since measurement error should produce uncorrelated displacement errors between frames (**Fig. 2G-J**. and **fig. S1B**). We refer to this as “ultra-resolution”, as it approaches the scale achieved with fixed-cell super-resolution methods (*69*, *74*, *75*), and as it represents overcoming not only the diffraction limit, but also the motion-blur and label-size limits that are barriers to resolution in live-3D-genome imaging. These improvements reduced the F-to-D ratio (**Fig. 2I**), enabled higher temporal resolution (**Fig. 2H**) and more frames per trace (**Fig. 2J** and **S6**); providing unprecedented detail of genome motion (**Fig. 2C vs D**).

Using this approach we imaged all 11 of our initial TRACK-IT clones in replicate for 30 minutes each, collecting 300-1700 traces (cells) per clone over two replicate experiments for each (**fig. S1B**). We imaged asynchronously cycling cells, and distinguished G1 from S/G2 by the presence of doublet foci in replicated S/G2 cells (**fig. S7**). The corresponding ∼75 Tb of images resulted in over 18 million data points (combined with the perturbation experiments and the second target locus described below, this increased the total number of publicly available live-cell data points from a previous total 655 thousand (**Fig. 2K-L and S1B**)), which were processed on custom infrastructure using a pipeline optimized for computational efficiency to accommodate our imaging throughput (see ***Methods***). This data provides a high resolution view of genome motion across chromosome scales.

### MSCD measurements obey patterns predicted by polymer physics

Since all of the pairs are tethered together within the chromosome, the physical constraint provided by the polymer nature of DNA leads to several predictions about the dynamics of the inter-locus distance. The high resolution data from multiple loci across chromosome scales by TRACK-IT provides an opportunity to study the relationship between 3D chromatin dynamics and the genomic separation, and test the predictions of polymer physics. For a polymer, it is expected that the mean squared change in distance (MSCD, a.k.a. two-point MSD) starts off with power-law scaling with exponent *b*, *M_2_*(*t*) ∼ *t^b^*, at small time intervals and asymptotically flattens out at a value equal to twice the population variance in distance (2<*R*^2^>) at long intervals due to the tethering effect. Shorter polymers are expected to asymptote sooner. For a freely-jointed polymer in a dilute solution (“ideal chain”), the scaling exponent is ½ (*30*, *57*, *76*), for densely-packed, unknotted (“crumpled”) polymers it is expected to be 2/5 (*77*, *78*) (for details, see **Supp. Polymer Physics 1**). The data from 11 initial TRACK-IT clones were largely consistent with the MSCD expected for crumpled polymers (**Fig. 3A**). MSCD from loci with short genomic separation rapidly saturates at the expected steady-state (**Fig. 3A**, thick bars), whereas for loci separated by 12 Mb or more, the MSCD never reached its expected steady-state during the experiment, nor flattened, but followed the 2/5th scaling across the nearly 4 orders of magnitude in time. Labels with intermediate genomic separations roughly followed the 2/5th scaling at short time scales and showed different levels of saturation corresponding to their tether length (**Fig. 3A** and **S8**), also as predicted by the theory for crumpled polymers (*77*, *78*).

**Fig. 3.**
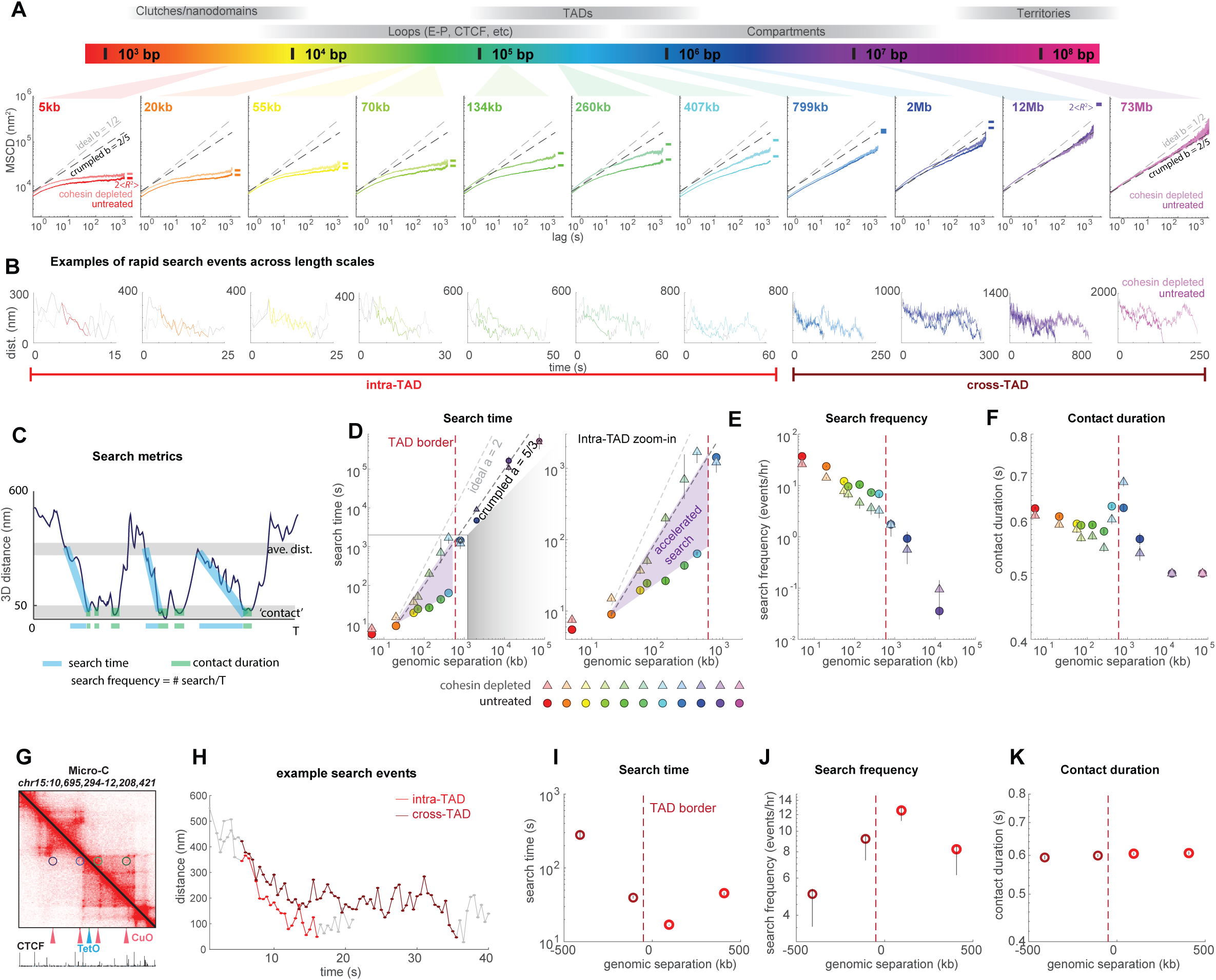
cohesin accelerates intra-TAD interactions. **(A)** MSCD as a function of genomic separation, denoted by color. Results after cohesin depletion are shown as lines with faded colors. Scaling exponents b = ½ (black dashed line) and = (gray dashed line) are overlaid. 95% confidence intervals are overlaid on each datapoint. **(B)** Examples of search events, which pass from the typical 3D distance of the observed pair to proximity, observed at all genomic scales. Gray lines show the motion for 5 seconds before and after the search event. Search events are colored as in (A). **(C)** Cartoon depiction illustrating the definitions of search time, contact duration, search frequency. **(D)** Median search time. Errorbars denote 95% confidence interval on the median. See **fig. S13** for distributions. Data from untreated cells is in circles, dTAG treated (cohesin depleted) cells in triangles, color coded by genomic separation as in (A). Power-law fits (lines on the log-log plot) are shown and the slope indicated in the legend. **(E)** Search frequency. Color code as in (D). Errorbars denote 95% confidence interval. **F.** Contact duration. **(G)** Micro-C of a TAD on chr15, with the position of the fluorescent TetO and CuO labels in the 4 cell lines marked. CTCF ChIP-seq is shown below for reference. **(H)** Example traces showing search events for an intra-TAD search and a cross-TAD search. **(I)** Median search times, as in (D), for the cell lines shown in (G). **(J)** Search frequency, for the cell lines shown in (G). **(K)** Contact duration, for the cell lines shown in (G). Contact threshold, 50 nm.

### Cis-interaction search-times suggest motor driven folding

Across all genomic scales, we captured events (*n=*21,805 events across the 11 lines) in which the spot-pairs transitioned from their typical (median) distance for the pair, to “contact” (<50 nm, as in our Hi-C comparison (**fig. S4A**) in a contiguous path spanning five to a few hundred frames (a few seconds to minutes), without returning to the average distance before reaching contact (see examples in **Fig. 3B**). Surprisingly, such transitions were often below the Nyquist sampling frequency limit for prior studies with 20 or 30 s resolution (**Fig. 3A**), and thus could not be reliably captured with earlier methodology (*27–30*, *57*, *79*), indicating that the 3D structures of the genome at the scale of loop-dots, TADs, and compartments, changes faster than previously anticipated or observed.

We further quantified the observed motion by computing the search time, search frequency, and the contact duration for each fluorescent pair (**Fig. 3C**). We defined the ***search time*** as the average time required to go from average distance for the given loci pair to contact (**Fig. 3C**). We defined ***search frequency*** as the number of times the pair transitioned from average distance to contact relative to the total observation time. Finally, we recorded the amount of time spent in contact as the ***contact duration*** (**Fig. 3C**). Any empirical measure of distributions of waiting times is affected by the finite nature of the observation window, which truncates processes that started before this window or do not complete by its end, leading to oversampling of shorter events. We used the Kaplan and Meier nonparametric estimator (*30*, *57*, *80*) to correct for this truncation effect (a. k. a. ‘censoring’ (*80*)), after validating the approach by comparing polymer simulations with different truncation times (**fig. S9A-C**).

Surprisingly, within the TAD, the median search time was less than 1 minute (**Fig. 3D**) and more surprisingly, the search time exhibited only weak dependence on the genomic separation. Indeed elements separated by 20 kb on average took only ∼20 s, those separated by 260 kb took ∼40 s, and for 407 kb, only ∼60 s. By contrast, we saw a dramatic increase in search times for more distal points that crossed outside the TAD, increasing abruptly to over 1000 s at 799 kb. Outside the TAD, we observed a stronger dependence on genomic separation as well as much longer search times. To more robustly measure these long search times and their scaling at megabase separations, we relaxed the contact threshold, and rescaled the values based on the original threshold (see **Methods** and **fig. S9C**). We benchmarked this approach with simulated polymer data truncated to different observation windows (**fig. S9C**). With this approach, we estimated the median search time for elements at 12 Mb to be ∼2e5 and 73 Mb to be ∼1e6 s (**Fig. 3D**).

The weak dependence of search time on genomic separation seen within the TAD is inconsistent with purely diffusive search according to current polymer theory, suggesting an *active* search mechanism. For the simple ‘ideal chain’ polymer, classic polymer theory predicts a power-law relation between the monomer-separation and the search time, with a scaling exponent, *a*, of 2 (*76*). Recent theory shows the diffusive search for crumpled polymers does not slow down as fast as a function of monomer-separation, with a lower exponent of 5/3 (*78*) (See **Supplement, Polymer Physics 3** for details). However, the measured search time scaling within the TAD (*a=*0.60, 95%-CI=(0.44, 0.76)) is not predicted by any of these models (**Fig. 3D**). By contrast the cross-TAD values did follow the predicted scaling for diffusing crumpled globules, *a*∼5/3 (**Fig. 3D**). These observations suggested that motor activity (e.g. by cohesin) may contribute to accelerated search and the flatter dependence of search time on genomic separation.

### Cohesin accelerates cis-interactions within a TAD

To test this hypothesized role for cohesin in these dynamics, we used dTAG-13 to acutely deplete the FKBP^F36V^-tagged endogenous RAD21, a core component of cohesin, from all 11 of the cell lines and imaged the cohesin-depleted cells in parallel with the untreated controls. We then examined the MSCDs for these cohesin-depleted samples (**Fig. 3A)**. Interestingly, particularly for labels separated by less than 600 kb we found the MSCDs are faster without cohesin (**Fig. 3A**, lighter lines), indicating that the cohesin slows chromatin movement rather than accelerating it. This observation is consistent with previous MSCD-analyses of loci at 150 kb and 505 kb separation with and without cohesin, and is reproduced in the MSCD analysis of simulations in our hands (see **Supplement, Polymer Physics 2**) and some recent work (*29*, *30*).

Despite this acceleration effect of cohesin depletion observed in the MSCDs, intra-TAD search times were substantially slower/longer in cells depleted of cohesin (**Fig. 3B** and **D**). Moreover, the discontinuity seen across the TAD border / at Mb-scale separation vanished after cohesin depletion, while the cross-TAD search times showed little change between the untreated and cohesin-depleted cells (**Fig. 3D**). Instead, search times in cohesin depleted cells exhibited a power-law increase in search-time as a function of genomic separation (*a*=1.5, 95%- CI=(1.3, 1.7)), consistent with the 5/3 scaling predicted from the diffusion theory for crumpled polymers (*78*). This indicated that cohesin is able to accelerate search times of sub-Mb separated elements within a TAD, without substantially perturbing the search behavior at larger separations or between TADs. Moreover, it illustrates that genome motion in mammals without cohesin is substantially diffusive, and that cohesin is responsible for the surprisingly rapid and uniform search times within TADs, which deviated from diffusive predictions. Furthermore, the variation in intra-TAD search time also increased upon cohesin depletion, while the distribution of search times for cross-TAD pairs exhibited little change (**fig. S13**). Similar search times were observed for G1 and G2 cells, with similar scaling (**fig. S10**). Smaller/larger contact thresholds had the expected effect of increasing/decreasing search times, but did not substantially change the qualitative scaling behavior observed (**fig. S11**).

### Cohesin increases search frequency within a TAD

In addition to changing how long it takes for the typical search event to occur, it is possible that cohesin also affects the frequency of search events per minute or hour (which could be regulated independently, for example by the frequency of cohesin loading, rather than the processivity and extrusion speed of individual cohesins which may affect the search time). We thus counted the number of search events and normalized this by the number of observations in each trace (**Fig. 3E**). As expected, the search frequency is rarer for elements separated by larger genomic separations. For example, within the TAD, at 55 kb separation we found 10 searches/hr, while at 407 kb separation we observed ∼7/hr, which dropped to little over 1/hr at larger separation across TAD borders (**Fig. 3E**). We observed a modest discontinuity when the second label is moved across the TAD border. When cohesin was depleted, the search events were observed less frequently at sub-Mb scales, for example, ∼7/hr at 55 kb or ∼3/hr at 407 kb - though they were not substantially changed at larger length scales (**Fig. 3E**), indicating that cohesin also increases the number of search events per unit time.

### Contact duration is independent of TADs and genomic separation

An affinity / microphase separation model of TADs (*81–83*) would predict longer contact durations for intra-TAD than cross-TAD contacts. Other differences in chromatin context and genomic separation may also affect dwell times. To explore this, we next computed the amount of time each pair spent in contacts as a function of the different genomic separation and chromatin contexts of our 11 pairs. Contact durations were corrected using the Kaplan-Meier estimator, adjusting for over-estimation of brief contacts due to the temporal resolution of our data. We observed an average of ∼0.6 s, with minimal dependence on intra vs. cross-TAD, genomic separation, or cell cycle for these 11 pairs (**Fig. 3F and S10**). A few minor variations are nonetheless statistically significant. We see a minor decrease in contact duration upon cohesin depletion within the TAD ≤407 kb (*p*=0.026, KS-test). This difference alone is unlikely to contribute to the emergence of the TAD in the population data, as the contact duration for the first cross-TAD pair (799 kb apart) is actually longer than most of the intra-TAD pairs, contrasting the prediction for affinity-microphase separation (*p*=0.018, KS-test). If loop-extruding cohesin leads to a slight increase in the contact duration of intra-TAD loci as they get extruded, this could explain the observed slight difference. A systematic comparison of 6 epigenetic marks in the vicinity of these sites (+/-2.5 kb) did not reveal any significant correlations between chromatin state and contact duration, (**fig. S11**). Higher temporal resolution may be needed to more sensitively dissect the effects of chromatin context on these brief interactions, which are close to the Nyquist resolution of our data.

While individual contact durations were short, most trajectories returned repeatedly to within the contact threshold following the initial search. We quantified the total contact time per search from the ratio of the absolute contact frequency (**Fig. 1E**) and the search frequency (**Fig. 3F**), which gave an average of ∼22 s of contact, or 36 contacts/per search. Interestingly, these values were rather uniform across all our measured label pairs with or without cohesin, (independent of separation distance), (20 s to 30 s) indicating that for these loci, the role of cohesin on the total contact frequency was largely through faster search, not longer dwell times.

### TAD borders regulate speed and frequency of cis-interactions

To better understand the effect of TAD borders, we conducted further experiments with a new target region in a ∼500 kb TAD on Chr15 (*84*). We placed the TetO label ∼50 kb inside the TAD boundary and inserted the CuO array at selected genomic separation-matched loci 100 kb upstream (cross-TAD) or 100 kb downstream (intra-TAD) (**Fig. 3G**). Two further cell lines were generated with the second label placed 400 kb upstream or downstream, which was approximately as far from the initial insertion while still being within the same TAD and ∼50 kb from the boundary (**Fig. 3G**). We then imaged the movement of these labels as described for the previous 11 lines and captured 18,295 more search events from the average distance to contact (**Fig. 3H**). Notably, the search times were substantially faster for intra-TAD pairs (**Fig. 3I**), while search events were also substantially more frequent (**Fig. 3J**). We again find the contact duration to be independent of genomic separation and the underlying TAD (**Fig. 3K**). Together with the data by TRACK-IT, these results again show that TAD borders regulate the speed and frequency of cis-interactions and not only their population average distances. Similar cross-TAD differences were observed for a range of threshold values from 30 to 150 nm, and similar strength of effect was observed in both G1 and G2 cells, across the 15 cell lines (**Figs. S10 and S11**).

### Cohesin extrusion produces processive movement on stochastically stretched chromatin, of 2.7 kb/s

While the statistical behaviors of 3D motion show a strong dependence on the presence of cohesin, we next asked if individual traces show signatures of active extrusion. Some of the search events, such as one seen in 407 kb separation, showed a tantalizingly processive motion that may be consistent with “reeling in” of intervening chromatin between the probes (**Fig. 4A**).

**Fig. 4.**
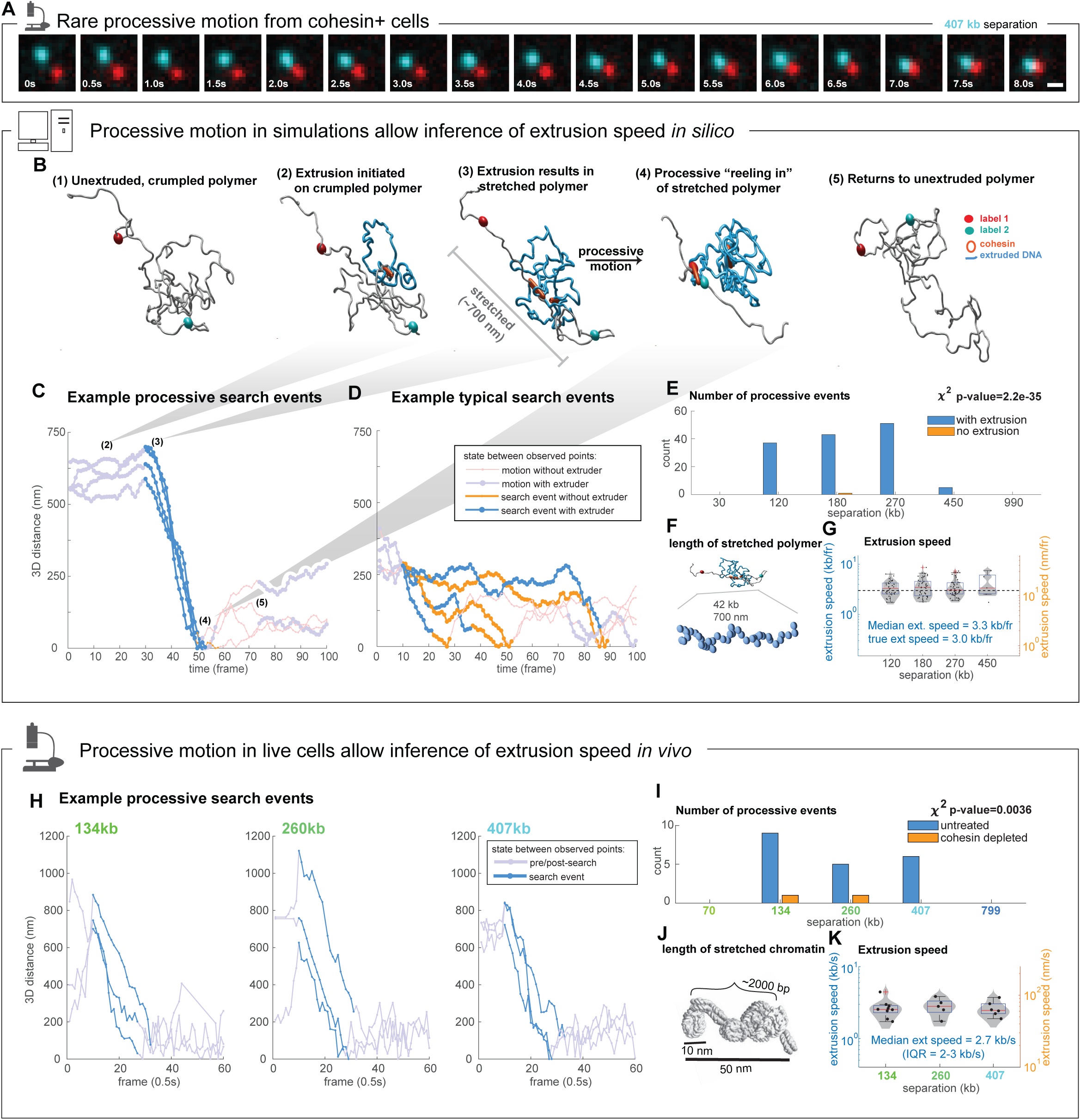
Inferring cohesin speed from traces with extended unidirectional motion. **(A)** Representative time-lapse images of processive motion observed in cohesin containing cells, for labels 407 kb apart. Scale bar, 500 nm. **(B)** Snap-shots from a simulation of loop extrusion in a crumpled polymer. Labels are colored as in (A), the extruded portion of the simulated polymer (gray) is shown in cyan. Gray bar at snapshot 3 shows the extent of the stretched polymer after cohesin has extruded the intervening slack. **(C)** Examples of processive search events as defined in the text from simulation of loop extrusion. X-axis shows simulation time, y-axis the 3D distance change over the observation window. The intervals in which one or more loop extruder (LE i.e. cohesin) occupied the interval between the two labeled loci are shown as thick blue lines. **(D)** As in (C), but typical, non-processive search events. Note the intervals in which a loop extruder is not present (thick orange lines). **(E)** Number of processive events by genomic separation (1.5kb per simulated monomer) between the labels. **(F)** Molecular dimensions of stretched chromatin in simulations of processive motion. **(G)** Violin and scatter plot of the inferred extrusion speed for 15 processive search events *in silico* (see text). **(H)** Examples of the 3D distance change over time for processive traces in cell lines with labels 134 kb, 260, and 407 kb apart. The processive portion (see text) is denoted in light blue, the preceding and subsequent motion in light purple. **(I)** Number of processive events by genomic separation between the labels. **(J)** Image of a stretched nucleosome fiber captured from electron tomography by ChomEMT and nucleosome modeling, used in estimating the length of stretched DNA. Image adapted from Ou et al 2017 (*85*). **(K)** Extrusion speeds inferred from traces of processive motion *in vivo*, shown as a violin plot. Individual data points shown as black dots, the mean and median are red and black lines.

Although rare, such a processive pattern is indeed predicted to occur from simulations of loop extrusion, when a cohesin that extrudes chromatin faster than chromatin diffuses is loaded near the center between the two labels that are sufficiently separated in 3D (**Fig. 4B**). Initially, such cohesin were primarily involved in extruding the intervening, crumpled, chromatin (**Fig. 4B** (**1**)**-**(**2**)). Once the cohesin finished extruding this slack, the intervening chromatin briefly adopted a stretched, largely linear configuration (**Fig. 4B** (**3**)), during which continued extrusion produced distinctively processive motion (**Fig. 4B** (**4**)). From this motion, the cohesin motor speed may be estimated from the slope of the processive trajectory and length scales of the stretched chromatin.

To test how accurately we could identify windows of active extrusion and measure cohesin extrusion speed from 3D-distance trajectories, we first analyzed results from established approaches for simulation of cohesin loop extrusion (*4*, *86*), where the position of cohesin and its applied forces are known precisely (**Fig. 4B**). We found processive motion in simulated traces always indicated the presence of an actively extruding cohesin molecule between the labels when the travel range at least twice the average 3D distance of the labels, with at least 16 consecutive frames, in which less than 20% reversed direction and in which the total reverse distance was less than 20% of the forward distance (**Fig. 4C**, blue lines). In contrast, search events across shorter 3D distances, shorter times, or greater fraction of reverse steps did not always contain an extruding cohesin between the labels, even in cohesin-positive simulations (**Fig. 4D** blue and orange lines). We observed a similar effect across a range of simulated genomic separations (**Fig. 4E**).

We also used these simulated traces to test and validate an approach to estimate the speed of cohesin extrusion *along the chromatin*, informed only by the 3D positions of two labels (since that is the only observable in our experimental data). We computed that the 700 nm distance in these simulations corresponds to approximately 42 kb of simulated stretched chromatin (**Fig. 4F**, also see **Methods**). We applied this conversion to all the simulated traces exhibiting processive motion, for loci pairs between 120 and 450 kb apart, which resulted in average extrusion speed of 3.3 kb/frame, in good agreement with true extrusion speed used to in the simulation that generated these traces of 3.0 kb/frame (**Fig. 4G**). We can also use these data to provide bounds on these estimates (see **Methods**).

We then searched for examples of such processive motion in the experimental data from TRACK-IT, with and without cohesin, using the same parameters that correctly isolated cohesin dependent movement in the simulation data. Data from the 407 kb label pair contained 7 events of processive motion in cells with cohesin (untreated) and none from cells following cohesin depletion (**Fig. 4H**). From pairs at 260 kb and 134 kb we observed 5 processive events each from cells with cohesin (**Fig. 4H**). Cells treated with dTag to deplete cohesin showed little processive motion - only 2 events were detected across our cell lines, possibly due to incomplete degradation. This specificity of processive traces to the untreated population (*p*=3.6e-3 chi-squared test) further suggests these are cohesin-driven events (**Fig. 4I**). More closely spaced pairs (<134 kb separation) did not exhibit processive intervals by these criteria, similar to our simulations at close intervals, possibly due to the rarity of obtaining a degree of linear distance that would take 8 s (16 frames) for cohesin to close. Loci at >407 kb also did not exhibit processive events, potentially due to the TAD border separating these and/or the larger genomic separation relative to the processivity of cohesin *in vivo* (**Fig. 4I**). This selectivity of processive events for the ∼134∼407 kb scale is consistent with the predicted cohesin loop extrusion-driven movements (**Fig. 4I vs. E**).

Remarkably, all three cell lines exhibiting processive events in the presence of cohesin exhibited a similar range of closing speeds of ∼70 nm/s (covering 600 nm to 1200 nm in 9 to 18 s, respectively) (**Fig. 4H,I,J**), suggestive of a single extruding motor. This paralleled the behavior we observed in simulation, where extrusion of the last 40 kb of DNA that produced processive motion was also driven by a single extruder and exhibited a constant speed, even when secondary extruders were nested within the slack (**Fig. 4C**). Calibrating with published measurements of chromatin fibers from *in situ* electron microscopy (**Fig. 4K**) and our observations from simulations (**Fig. 4B, E**), these distances correspond to 24-48 kb of stretched chromatin. This provides an estimate for each processive trace of the extrusion speed in both kb/s and nm/s (**Fig. 4L**). All three cell lines exhibit a similar average speed of 2.7 kb/s, with an interquartile range of 2.3-3.0 kb/s (**Fig. 4L**), modestly faster than average ∼1 kb/s observed *in vitro* under opposing flow (*25*, *26*).

## Discussion

Here we examined genome motion across length scales from 5 to over 73,000 kilobases with high spatial and temporal resolution, using TRACK-IT. These data provide a quantitative picture of the mean-squared change in displacement, the search time, search frequency, and contact duration, across genomic length scales. We found cells exhibit superdiffusive search times for sequences within the same TAD in a cohesin-dependent manner. By contrast, cross-TAD search times and searches in cells lacking cohesin showed a surprising agreement with the scaling predicted from a theory of crumpled, unknotted, polymers moving diffusively, with modest deviation likely due to other effects secondary to diffusion (**Fig. 3D**). We found search times with cohesin within TADs were surprisingly fast, 1 minute or less for all points within the two TADs examined, and less than 20 seconds for the 100 kb scale common to many known cis-regulatory interactions (**Fig. 5**). These search times increased substantially upon cohesin depletion, with the strongest effects seen at the greatest intra-TAD separations. We also detected periods of surprisingly processive, cohesin-dependent motion, enabling an estimate of the loop extrusion speed of cohesin on single DNA molecules *in vivo*. Together, these observations have implications for our understanding of the nature of TADs, for understanding the relative contributions of diffusion and motor-driven assembly underlying genome structure across multiple scales, and for the speed and precision of long-range transcription regulation. While we restrict our conclusions to the genomic regions of chr6 and chr15 of mESCs studied, we speculate here how these properties might extend more generally across cohesin-dependent TADs and discuss how our measurements compare to previous estimates and predictions for genome motion.

**Fig. 5.**
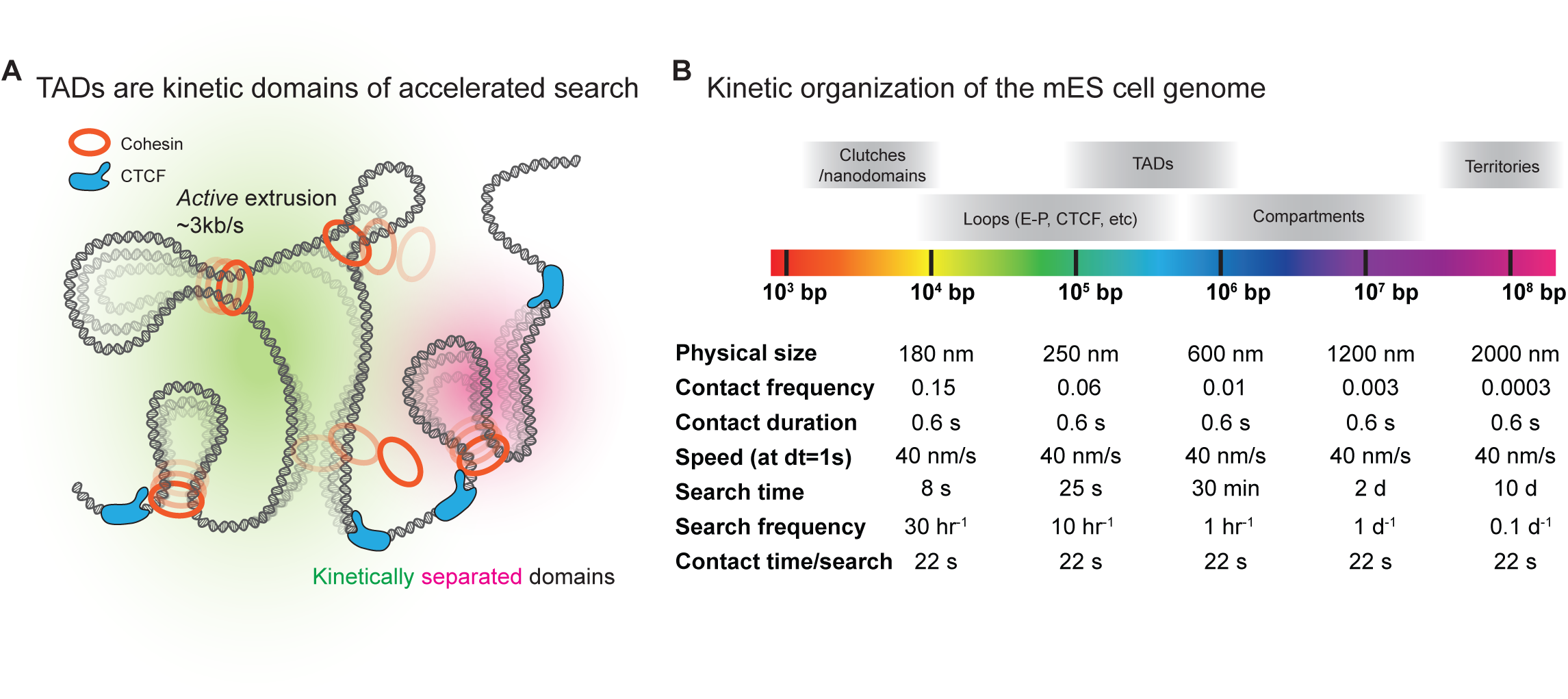
**Kinetic organization of the genome**. **(A)** genome is organized into kinetic domains, where within, experience cohesin-accelerated search times. TAD borders (CTCF binding sites) function as kinetic, not structural, barriers. **(B)** Summary table of some key molecular and kinetic properties of chromatin measured here as a function of genomic separation.

### The speed of cohesin loop extrusion

The high temporal and spatial resolution of our approach enabled direct observation of the processive motion predicted from cohesin loop extrusion in live cells. While we cannot definitely know that processive motion is directly driven by a cohesin at the base of the extruded chromatin loop as simultaneously imaging cohesin would be prohibitively difficult, two lines of evidence support our claim. First, observed processive motions are cohesin-dependent; almost exclusively seen in untreated cells, but not in cohesin-depleted cells. Second, processive traces were found only in specific inter-label genomic separations (134∼407 kb), as predicted by simulations of loop extrusion. This is due to the opposing requirement for large initial 3D distance to allow for multiple consecutive steps (which is unlikely for small genomic separations) and small genomic separations that allow for a single extruder to processively extrude the intervening chromatin (which is unlikely in larger genomic separations). These evidences argue for a non-diffusive, directed motion indicative of loop extrusive motor activity, allowing for a direct estimation of *in vivo* speeds of active loop extrusion.

Previous work has approximated the *in vivo* extrusion speed at 0.1 kb/s (*30*, *87*), at face value notably different from ∼2.7 kb/s indicated in our data (**Fig. 4K**). This earlier estimate was derived from live-cell fluorescence recovery after photobleaching (FRAP) of fluorescently tagged cohesin in G1 cells, which estimated the residence time to average ∼20 minutes (*88*), combined with fitting polymer models to Hi-C data, which estimated the average cohesin loop size at ∼120 kb. While this ratio of average loop length to average residence time gives an intuitive estimate of extrusion speed (0.1 kb/s), it uses the simplifying assumption that all the residence time is spent in active extrusion, and that the average loop size is set by the speed and lifetime (rather than the density of extrusion blockers, for instance). A recent preprint also reports that simulations using a 0.1kb/s extrusion speed, also assuming constant and uniform extrusion, gave the best agreement to the “closing speed” inferred from the live-cell data of three different TADs (*89*). Should chromatin associated cohesin sometimes exist in a stalled state (due to NIPBL loss, PDS5 binding, and/or CTCF stalling, etc.) (*90*), these approaches will only provide a lower bound to the speed of active extrusion. Notably, recent modeling of the cohesin biochemical network from *in vivo* protein abundance and binding kinetics data concluded that actively extruding, NIPBL bound cohesin constitutes a minor (∼15%) fraction of total G1 bound cohesin (*90*), possibly bridging rapid speed of active loop extrusion seen in in our processive traces with previous estimates.

Previous *in vitro* live imaging of cohesin recorded extrusion speeds averaging 1 kb/s (*25*, *26*), closer to, but still slower than our *in vivo* estimates. The absence or sparsity of nucleosomes in *in vitro* studies may lead to a slower observed extrusion time, as cohesin may need to take more steps to reel in this less compact polymer (e.g. a full complement of nucleosomes would shorten the chromatin polymer by 6 fold *in vivo*). Supporting this, *in vitro*, it has been shown that nucleosomes (and larger particles) do not provide a barrier to extrusive cohesin (*26*, *91*), and the ‘swing and clamp’ mechanism proposed for cohesin walking (*92*), would cover more DNA bases in a single swing if that DNA is compacted around intervening nucleosomes. Further, these *in vitro* speeds were measured under fluid flow to stretch out un-extruded DNA and increasing the flow rate reduced the extrusion speed (*26*). Should the extrusion events that we captured *in vivo* have experienced less opposing force, this could contribute to the faster extrusion rates estimated. We also note that our approach based on processive motion may capture only the faster extrusion events – those that extrude substantially slower might allow time for the labels to both diffuse closer and drift further apart during the extrusion window, which we would classify as non-processive motion.

### TADs are kinetic domains of accelerated search

We found loci pairs within the same TAD exhibited significantly faster transitions between their typical 3D distance and contact than observed for loci in separate TADs, and made significantly more transitions per hour. Removal of cohesin significantly slowed intra-TAD interactions while having little measurable effect on the speed or frequency of longer-range, cross-TAD interactions. This suggests that the TADs studied here should be understood as domains of distinctive *kinetic*s, at least in addition to, if not instead of, being understood as genomic *structures*, as sometimes depicted (**Fig. 5A**) (*93*, *94*). The kinetic nature of TAD offers an intuitive solution to the challenge posed by the extensive heterogeneity of genome folding seen in individual cells as shown by both fixed-cell (*20*, *75*, *95*, *96*) and live-cell measurements (*29*, *30*), in which the “representative” structure mirroring the population average is negligibly infrequent and thus cannot reasonably account for regulatory function. Rapid intra-TAD kinetics would allow for elements to quickly shuffle through possible microstates (the individual single-cell structures found in fixed-cell measurements) within their borders, and facilitate statistical averaging needed for genomic processes seemingly regulated by underlying TADs in single cells. Notably, these kinetically separated domains do not necessitate often assumed spatial segregation from juxtaposing domains (which are often modest or absent in single cells (*20*, *75*, *95*, *97*)) nor TAD border looping (which are infrequent (*20*, *96*) and dynamic (*29*, *30*)) .

### Implications of these multi-scale dynamics for cis-regulatory control of gene expression

Second-scale chromatin motion at the TAD scale may allow for more robust and sensitive enhancer regulation. Transcription activity at the single-cell level, correlates with changes in contact frequency, measured at the population scale (*69*, *98*). However, differences in contact frequency are often modest (a factor of two on average), even across borders whose deletion results in on/off changes in gene-expression (*98*). If contacts with distal elements within the TAD typically took tens of minutes, then any system responsive to contact frequency would need to observe the process for many hours in order to robustly sense the observed modest differences in contact frequency. Such a long sensing window is difficult to reconcile with the speed of transcriptional responses in developing systems. However, we found that sequences separated by 5-400 kb (the length scale of most enhancer-promoter communication) conduct searches in tens of seconds which occur at frequencies on the scale of minutes. Such rapid surveillance would allow for modest contact differences to be sensed reliably on the time scale of transcriptional response. Analogous arguments have been made for the design of enhancers sensitive to small fold changes in TF concentrations (*99*) or for chemotaxing bacteria to sense small fold changes in chemoattractant (*100*). Certainly, much work remains to be done understanding the molecular mechanism behind this contact frequency-dependent activation, though some models have been suggested (*84*, *98*). We also note that while cohesin had a profound effect on the search times at genomic separations of >100 kb, the search times for elements separated by 5-70 kb showed rather milder differences. This suggests a further explanation for recent findings that enhancer-promoter communication of several genes was largely insensitive to cohesin loss at separations of up to 20 or 100 kb (depending on the experiment), but quite sensitive at separations of >100 kb (*101–105*).

These rapid changes in the 3D distance of intra-TAD elements have implications for the interpretation of recent live cell experiments. In particular, recent studies examined the correlations between enhancer and promoter labels and transcription activity (*28*, *56*, *106*). Our data show that elements separated by equivalent genomic separations often exhibit “kiss-and-run” (*107*) behavior within the 30s window used in these studies. Such transient interactions also offer an explanation of why the correlations detected between enhancer and promoter 3D positions and nascent transcription in fixed cells are weak (albeit statistically significant in many cases) (*69*, *108*). In the 30s-120s required to produce a nascent transcript of sufficient length for robust detection by FISH, there is ample time for the distance to have increased beyond the contact threshold, and ample time for a genomic region not driving expression to come into contact with the gene. While our measurements focused on the motion of observation points along the chromatin polymer, rather than track individual enhancer-promoter pairs, these search times, search frequencies, and contact durations provide a baseline (**Fig. 5B**) from which to make predictions for regulatory elements, which are also subject to the motion of the chromatin polymer.

In contrast to the rapid motion at the intra-TAD scale, we found elements separated by TAD borders to exhibit long search times, steeply scaling with genomic separation, consistent with the theoretical predictions of a crumpled polymer. For elements separated by 10s of megabases, the average search time (2∼10 days) is estimated to be longer than the cell cycle length (∼0.5 day) of the mouse ES cell under study, suggesting that, on average, these chromosomal-scale distal elements are unlikely to reliably come into contact before the next cell division. This observation implies that examples of extreme long-range cis-regulation (*109–112*) may require significantly longer cell cycles (or be post-mitotic), and/or additional mechanisms to bridge these large separations (compartments/phase-separation (*11*, *112*), inverted nuclei (*12*, *111*, *112*), etc). Indeed, many of the previous examples of long-range contacts are found in long-lived postmitotic neuronal cells in mammals (*109*, *111*, *112*) and *Drosophila* (*110*).

## Limitations

Our study is subject to several limitations. Our analyses were conducted only in embryonic stem cells, and it is possible that the kinetic properties of the genome in the pluripotent state are vastly different than in later development or in different terminally differentiated lineages. Our experiments focused on a series of labels incorporated on just two chromosomes, with each series spanning one prominent TAD boundary. It is conceivable that the kinetic properties of TAD boundaries vary substantially across the genome, and may be modulated by different cofactors also enriched at these borders. It is possible that binding of the TetR and CymR with their StayGold and Halo-tag cargos substantially alters genome kinetics, even though the impact on the contact frequency in a population snap-shot was not measurably changed. We have endeavored to minimize this potential by shrinking the size of the bound regions compared to earlier work, and by selecting a fluorescent labeling system that does not melt the DNA double helix or stall the progression of DNA (or RNA) polymerases, but we cannot rule out a kinetic impact of labeling.

## Acknowledgments

We thank Luca Giorgetti and members of his team and Anders Hansen and members of his team for helpful discussion and feedback on our results. We thank Simon Grosse Holz for helpful discussions about data analysis and polymer physics.

## Author contributions

Conceptualization: JL, LC, ANB

Methodology: JL, SG, ANB, AS

Investigation: JL, LC, KG

Visualization: JL, ANB

Funding acquisition: ANB, AS, SG

Project administration: JL, ANB, AS

Supervision: ANB, AS

Writing – original draft: JL, LC, SG, ANB

Writing – review & editing: AS

## Data and materials availability

Software developed for this project is available at https://github.com/BoettigerLab/TRACK-IT.git. Software for microscope control is available at https://github.com/BoettigerLab/nanoscope-control.git. This repository is a fork of the storm-control software developed in the Zhuang Lab (*116*). Software for polymer simulations is available at https://github.com/BoettigerLab/polychrom. This repository is a fork of the polychrom software developed by the open2c consortia (*86*).

Transposon mapping data (isT-seq) were deposited into GEO under accession number GSE289566 and are available at the following URL: https://www.ncbi.nlm.nih.gov/geo/query/acc.cgi?acc=GSE289566. Live cell tracking data has been submitted for hosting at the 4DN data portal and is under review. Accession numbers will be updated when available. [Submission in Process]. Data from prior ChIP-seq analysis for K27ac, K27me3, K36me1, K36me3 and CTCF, re-used in this work, is available on GEO GSE146451, K9me3 data GSE180003, and ATAC-seq data GSE99746 (GSM2651154).

## Supplementary Material

Materials and Methods

Supplement: Polymer Physics Figs. S1 to S13

Tables S1 to S3 References (1–164)

