## Supplemental Material for "Kinetic organization of the genome revealed by ultra-resolution, multiscale live imaging"

**The file includes:**

Materials and Methods

Supplement: Polymer Physics

Figs. S1 to S13

Tables S1 to S3

References

**Materials and Methods**

1. **Optimization of fluorescent labels for DNA tracking**
2. **Generation and TRACK-IT plasmids**
3. **Generation and culture of cell lines**
4. **Imaging and calibration**
5. **Image analysis**
6. **Trajectory analysis**
7. **Loop extrusion simulations**
8. **Other analysis**

**1. Optimization of fluorescent labels for DNA tracking**

In order to improve resolution and signal to noise ratio (SNR) of our fluorescent DNA labels, we began by evaluating strengths and weaknesses or recent approaches. Chromatin live imaging methods largely fall into two categories, methods dependent on genome editing and methods using transient transfection. The first group of methods, includes the widely used fluorescent repressor-operator systems (FROS), which require insertion of exogenous labels that bind fluorescent probes into the genome. FROS have the advantage of using bacterial repressors that have much stronger DNA binding affinity to its consensus sequence (compared to mammalian TFs) and no endogenous binding sites in the mammalian genome (*28*, *114*–*116*), allowing for an array of operator sequences to serve as a highly specific and robust fluorescent label. In addition, DNA binding for many bacterial repressors is allosterically regulated by small molecules, such as tetracycline/doxycycline and cumate for TetR and CymR respectively, allowing inducible binding to the labels even when constitutively expressed. However, a large number of repeated operator sequences (>100) are often needed to recruit a sufficient number of fluorescent repressors. Highly repetitive and large size of operator array can lead to difficulties in its use in genome editing as well as potential heterochromatization of the insert (*28*, *73*, *117*).

The Anchor system utilizes prokaryotic ParB proteins which bind to centromere-like ParS sequences (*42*). In contrast to the FROS system with fixed number of repeat numbers, ParB proteins spread from the ParS sequence up to 10kb away through bridge mediated 3D spreading (*118*, *119*). While this property is useful for allowing ParS to serve as a robust label while being shorter than most FROS arrays, uncertainty in exact extent of ParB spreading at single cell level can limit spatial resolution (**Fig. 2E-G**).

MS2/PP7 RNA aptamers inserted and expressed from a loci of interest are sometimes used as a proxy for the loci position (*57*, *79*, *106*). These labels by nature require the loci of interest to be actively transcribing. As the loci can accumulate multiple MS2/PP7 transcripts over a transcriptional burst, physical size/proximity to the loci of interest likely vary over time and pose a limit to spatial resolution.

The second group of methods utilizes fluorescently labeled endonuclease-dead CRISPR/Cas9 or Cas12a to directly target endogenous sequences (*31*, *44*, *50*, *120*). Importantly, these methods bypass the need for label insertion. Yet, enough unique gRNAs or signal amplification schemes are still required for robust labeling and often require optimization at individual target loci. Moreover, constitutive expression of dCas9 is associated with genome instability (*121*), hindering its use as fluorescent label orthogonal to genome function. Furthermore, label density (number of dCas9/12a binding sites/genomic size) is likely lower than FROS labels as gRNAs need to be sufficiently long (~20bp) to ensure binding specificity in addition to requirement for PAM sequences which may be limiting.

In this study, we chose the FROS system for its robust labeling and fixed label size. In particular, we focused on the smaller 48-mer repeat sizes with randomized spacer sequences reported by Tasan et al. (*73*). These shorter labels have multiple advantages over previous FROS labels, 1) shorter label lengths for improved spatial resolution, 2) less perturbative to genome function, and 3) direct manipulation with PCR and other molecular cloning methods (*73*). For the second orthogonal label, we constructed a 48-mer CuO array with randomized spacer sequences through direct chemical synthesis, enabled by the smaller number of repeats and randomized spacers. Both TetO and CuO arrays have previously been used in chromatin live imaging and characterized to have little impact on genome organization even at larger repeat sizes (>100). We note that other FROS systems remain good candidates for miniaturization, including LacO/LacI (*29*, *122*) and Mut-TetO/Mut-TetR (*27*). However, additional considerations may be needed for these candidates such as potential genome instability of LacO arrays (*27*, *123*) and heterodimerization between wild type TetR and mut-TetR (*27*).

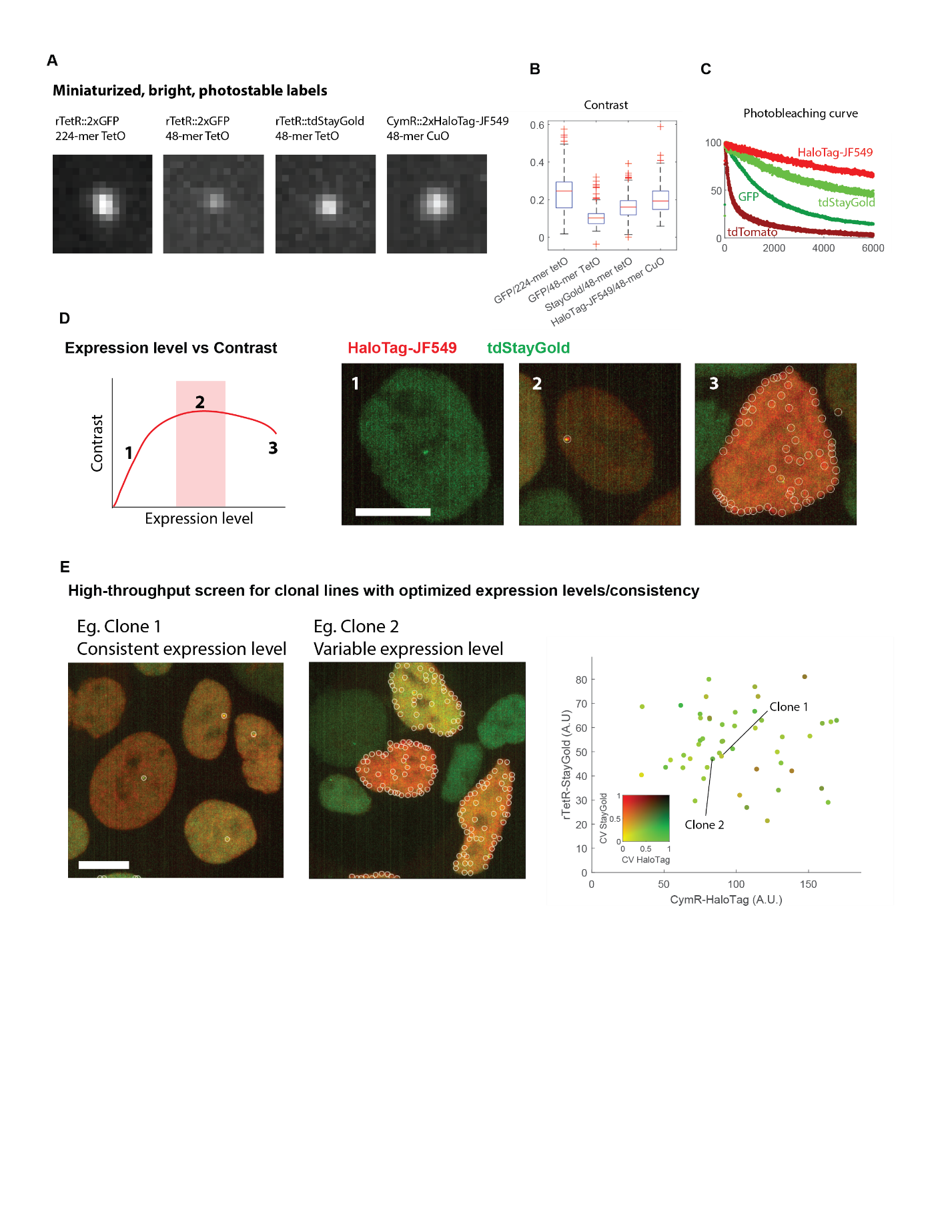

**Methods Fig. 1. Optimization of fluorescent labels for DNA tracking**. **(A)** Representative images of the fluorescent labels visualized in mESCs with different array lengths and fluorescent repressors. **(B)** Box plots of contrast (Intensityspot-IntensitybackgroundIntensityspot+Intensitybackground) of the four fluorescent labels.  *n* = 232, 352, 351, and 168 cells, respectively. **(C)** Photobleaching curves for fluorescent labels tested for DNA tracking. **(D)** left, schematic of expression level on contrast of the fluorescent label. Right, representative images of three regimes of CymR-HaloTag-JF549 expression level and effects on DNA tracking. White circles, fitted spots using Laplacian of Gaussian method. **(E)** left, representative images of two clonal lines showing the effects of consistent or variable expression of the fluorescent repressors. Right, example quantification of mean rTetR-tdStayGold and CymR-HaloTag-JF549 expression levels. Each data points were colored according to their coefficient of variation. *n* = 49 lines. Scale bars, 5 μm

The drawback of smaller repeat arrays is the reduced signal intensity, limiting its use for live imaging. We indeed found that our 48-mer TetO array has 2.4-fold reduced contrast against the nuclear background compared to the ~5-fold longer 224-mer TetO array both bound to rTetR-2xGFP (**Methods Fig. 1A-B**). We however note that while being ~5-fold shorter, 48-mer array was not 5-fold dimmer, suggesting that the label signal intensity does not necessarily linearly scale with the repeat sizes, with shorter arrays likely to have its binding sites more efficiently occupied by fluorescent repressors. This observation is in line with recent single molecule footprinting results of unsaturated (~20% occupancy) TetR binding to TetO arrays in mammalian cells by nucleosomal occlusion (*124*). It may be possible that larger arrays are prone to stronger nucleosomal occlusion due to its repetitive nature.

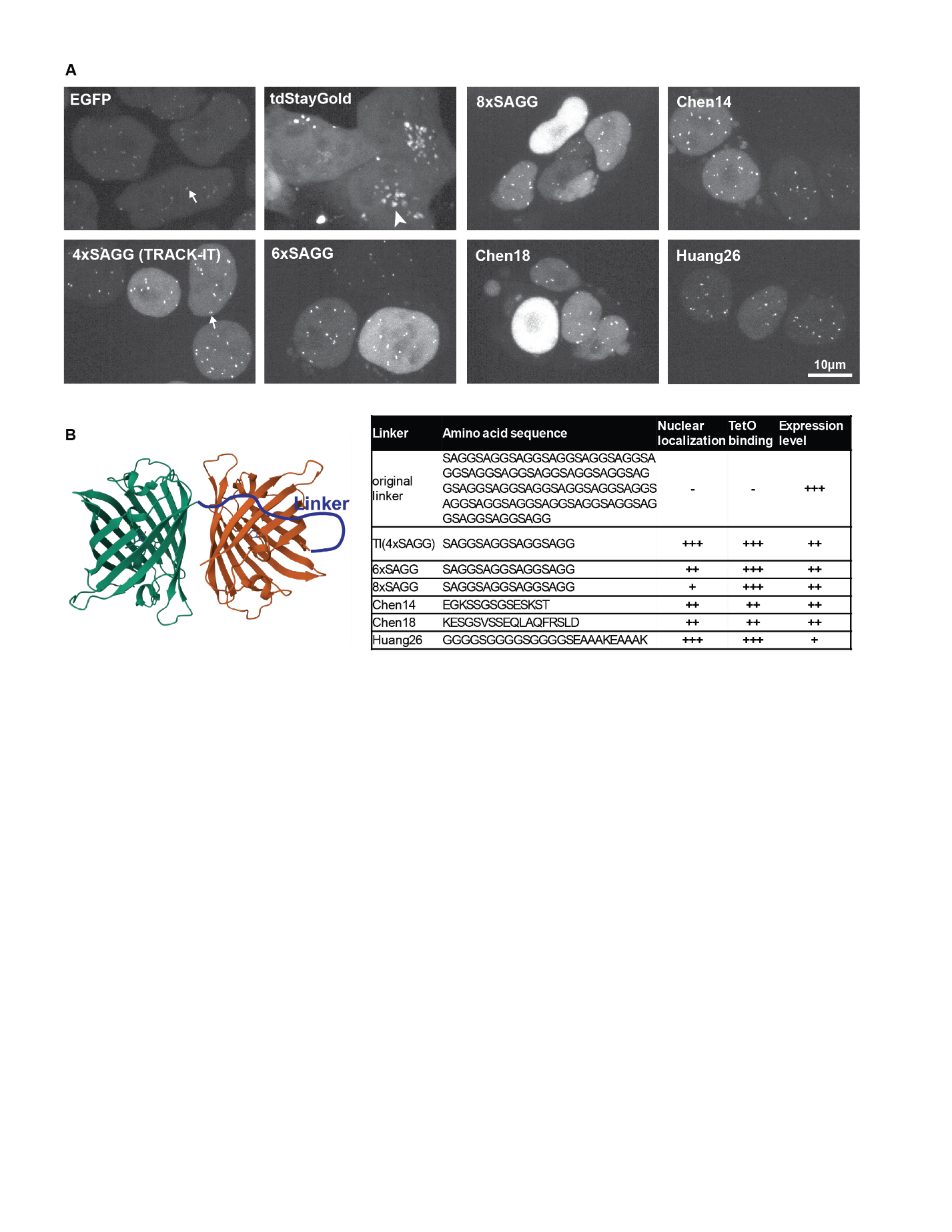

**Methods Fig. 2. Optimization of TI-StayGold in mouse ES cells.** Representative images of mESCs with randomly integrated 48-mer TetO arrays expressing rTetR tagged with GFP, tdStayGold or tdStayGold with alternative candidate linker sequences. Arrows, examples of rTetR bound to TetO arrays which appear as salt-and-pepper pattern in the nucleus. Arrowhead, example of mislocalization of tdStayGold to cytoplasmic bodies. Table, amino acid linker sequences and assessment of candidate linker sequences when transiently expressed in mESC cells.

We sought to overcome reduced signal intensity with brighter fluorescent proteins (ie StayGold) as well as synthetic fluor-conjugatable tags such as HaloTag. In addition to their increased brightness, both StayGold and JF549 (when conjugated to HaloTag), show stark improvement in their photostability compared to widely used fluorescent proteins GFP and tdTomato (**Methods Fig. 1C**), allowing for a larger total photon budget for photobleaching-constrained live-imaging experiments. Unfortunately, expression of rTetR-tdStayGold (tandem StayGold) in mouse embryonic stem cells resulted in aberrant mislocalization to cytoplasmic bodies (**Methods Fig. 2**). As the fluorescence appeared to be retained, we tested whether the localization defect could be rescued with a different linker between the two StayGold moieties. The original linker (EV linker) consists of 19xSAGG repeats, where its excessive length may hinder proper dimerization. Based on the available crystal structure of StayGold dimer(**Methods Fig. 2B**) (*125*), we posited that the flexible linkage between the two moieties would be achieved with significantly shorter linkers. As such, we cloned and transiently expressed 6 candidate linker sequences, shorter variants of SAGG repeats (4x, 6x, or 8x SAGG) and three other flexible linkers reported previously (**Methods Fig. 2**) (*126*, *127*), to test their proper expression and localization in mESCs with randomly integrated 48-mer TetO arrays. Rigid linkers were not considered as they are likely to hinder dimerization. While we found all six candidates to show improved localization, 4xSAGG linker overall showed best properties and thus was selected for future experiment.

Optimal contrast from the background is achieved at relatively low expression levels as signals from label-bound fluorescent repressors are diluted by unbound background signals (**Methods Fig. 1D**). Another perhaps more important consideration was the variability between single cells in a given field of view (FOV), as a single set of fitting parameters was used to fit spots in each FOV during image analysis.  Large variability between single cells results in more frequent false negative fitting of dimmer spots and/or false positive fitting of background signals (especially at nuclear edges). Therefore, we reasoned that the best strategy to achieve consistent, low levels of expression was through multi-copy integration of fluorescent ligand-expressing vectors under a constitutive but weak promoter such as UbC promoter (*128*).  To optimize expression parameters of rTetR-tdStayGold and CymR-2xHaloTag, we tested multiple integration conditions for respective plasmids using PiggyBac system.  We note that rTetR-tdStayGold, even with improved linker sequences, had overall lower levels of expression compared to GFP. To compensate for the reduced expression of rTetR-tdStayGold and brighter signals of CymR-2xHaloTag, we integrated these two ligands at 4 rTetR-tdStaygold:1 CymR-2xHaloTag molar ratio using PiggyBac system. Integration at 1:1 or 2:1 molar ratio largely resulted in too high or inconsistent expression of CymR-2xHaloTag.

Clonal lines were generated using FACS into 96-well plates and screened for expression levels for both fluorescent ligands (**Methods Fig. 1E**). Expression levels of individual clones were determined by first nuclear segmentation using CellPose (*129*) to create nuclear masks. Mean expression level and coefficient of variation for rTetR-tdStayGold and CymR-2xHaloTag were determined from raw images using regionprops function within Scikit-image (ver 0.24.0) (*130*) for each single clone, with cutoff for at least 50 cells measured. The clone with relatively low expression level and the least coefficient of variation for both channels were chosen for further experiments. We found that StayGold and HaloTag-JF549 bound to 48-mer arrays indeed resulted in stronger contrast compared to the GFP (**Methods Fig. 1B**).

**2. Generation and TRACK-IT plasmids**

TRACK-IT Cargo plasmid was generated as follows: First, the 48-mer CuO array plasmid was directly synthesized (GenScript). Sleeping Beauty 5’ and T7p::3’ ITR were synthesized as gBlocks (IDT) with 24-bp homology arms flanking the CuO array. The ITRs were cloned into 48-mer CuO array plasmid, linearized with EcoRI and KpnI, using Hi-Fi NEBuilder DNA Assembly (NEB, E2621L) following manufacturer’s protocols. Next, Sleeping Beauty transposon containing CuO array, SV40 promoter, and NeoR selection marker gene were amplified using Primestar GXL DNA polymerase (Takara, R051A) then cloned into 48-mer TetO array plasmid (generously gifted by Dr. Huimin Zhao) linearized with SpeI. Homology arms were amplified from CastX129 genomic DNA using Primestar GXL DNA polymerase and then cloned into the Track-IT Cargo plasmid linearized with EcoRI and NdeI.

To target TRACK-IT to the 129 allele in CastX129 mouse ES cells, 129 allele-specific gRNA sequence at chr6:51,320,704 (mm10) locus (TCAGATATGCTAAGGTAAAG) were cloned into Cas9 and sgRNA expression plasmid px458 (generous gift from Dr. Feng Zhang (Addgene plasmid # 48138 ; http://n2t.net/addgene:48138 ; RRID:Addgene_48138)) linearized with BsaI. Similarly, 3’ end of Rad21 coding region (chr15:51,964,057) targeting sgRNA (CTCAGATAATATGGAACCG) was cloned into px458 to generate Rad21 degrons.

rTetR-tdStayGold plasmid was generated as follows. tdStayGold sequence was directly synthesized (GenScript) and cloned into epB_CAG_TetRFlag-nls-GFPx2_DEx2 plasmid (a gift from Orion Weiner (Addgene plasmid # 119910 ; http://n2t.net/addgene:119910 ; RRID:Addgene_119910)) linearized via PCR amplification. Linker variants were generated by cloning in linker sequences synthesized as duplexed DNA ultramer (IDT) into rTetR-tdStayGold plasmid linearized via PCR amplification. See **table S1** for all PCR primers and synthesized DNA sequences.

**3. Generation and culture of cell lines**

**3.1 Cell culture**

Mouse embryonic stem cells (CastX129 hybrid F123 mESCs) were cultured in nDiff227 (Takara, Y40002) serum-free media supplemented with 2i/LIF (1μM PD0325901 (Stemcell Technologies, 72184), 3 μM CHIR99021 (Stemcell Technologies, 72054)), and 1000U/ml ESGRO mouse LIF (Sigma, ESG1107)). mESCs were cultured in 6-well plates pre-coated with 0.0015% poly-L-Ornithine (Sigma-Aldrich, A-004-C) and 10 µg/mL Laminin (Thermo-Fisher, 23017015). mESCs were fed daily by replacing with fresh media and passaged every two days with 0.25% Trypsin-EDTA (Gibco, 25300062). Each cell line generated by the study can be found in **table S2**.

**3.2 Genome editing and TRACK-IT cargo insertion**

To stably integrate rTetR::tdStayGold and CymR::tdHaloTag, 350,000 mESC cells were plated onto a 6-well plate the day before transfection. Cells were co-transfected with 1.2 μg ePB_UbC::rTetR::tdStayGold, 0.4 μg epB_UbC_CymRV5-nls-Halox2_DEx4 (a gift from Orion Weiner (Addgene plasmid # 119907 ; http://n2t.net/addgene:119907 ; RRID:Addgene_119907) and 1.2 μg Super PiggyBac expressing plasmid using 5 μl Lipofectamine 2000 (Thermo Fisher, 11668027). 5 days following transfection, rTetR::tdStayGold and CymR::tdHaloTag expressing cells were single cell sorted onto 96-well plates pre-coated with 0.0015% poly-L-Ornithine (Sigma-Aldrich, A-004-C) and 10 µg/mL Laminin (Thermo-Fisher, 23017015) using Fluorescence Activated Cell Sorting (FACS), gating for both tdStayGold and HaloTag-JF549 signals. 8 Days following FACS, plate was duplicated using 0.25% Trypsin-EDTA (Gibco, 25300062) onto a glass-bottom 96-well plate (Cellvis, P96-1.5H-N) pre coated with 10 μg/ml human fibronectin (Milipore, FC010-5MG) and an uncoated 96-well plate with 2i/LIF media supplemented with 10% FBS and 10% DMSO to facilitate freezing. Next day, glass bottom 96-well plates were screened for stable, optimal expression of both rTetR::tdStayGold and CymR::tdHaloTag.

CRISPR/Cas9-mediated genome editing was performed as previously described (*131*). 350,000 rTetR::tdStayGold and CymR::tdHaloTag expressing mESC cells were plated onto a 6-well plate the day before transfection. Cells were co-transfected with 1.2μg Cas9/sgRNA expressing plasmid and 1.2μg TRACK-IT Cargo donor plasmid using 5 μl Lipofectamine 2000 (Thermo Fisher, 11668027) following manufacturer’s protocols. Two days following transfection, antibiotic selection was performed with 2i/Lif media supplemented with 6 μg/ml Blasticidin (Thermo Fisher, A1113903) until resistant mESC colonies were evident. Resistant colonies were then passaged once and then single-cell sorted on 96-well plates by FACS. 8 days following FACS, 96-well plates were duplicated and genotyping was performed via genomic DNA extraction with QuickExtract™ DNA Extraction Solution (VWR, 76081-766) followed by qPCR to identify genome-edited colonies. Single copy insertion was validated with qPCR for a genomic locus and inserted sequence in addition to directly quantifying TetO and CuO label fluorescence intensities. Due to the less repetitive, smaller nature of our TetO and CuO arrays, no obvious genomic instability was observed.

To insert pairs of 48-mer TetO and CuO arrays across the TAD boundary (~chr15: 11,387,000), we chose to directly engineer in the shared TetO label and genomic separation-matched CuO label sites. 129 allele-specific insertion sites were selected based on sgRNA with Cast allele-specific mutations in the PAM sequence. sgRNA and Cas9 expressing vectors were cloned as described. Homology donors were PCR amplified from 48-mer TetO or CuO plasmids with flanking 50 bp homology arms to the target sites in a 200ul reaction with Primestar GXL polymerase. Correctly-sized amplicons were confirmed on an agarose gel and purified via ethanol precipitation. 350,000 rTetR::tdStayGold and CymR::tdHaloTag expressing mESC cells were plated onto a 6-well plate the day before transfection. Cells were co-transfected with 1.6μg Cas9/sgRNA expressing plasmid and 0.8 μg purified 48-mer TetO donor DNA using 5 μl Lipofectamine 2000 (Thermo Fisher, 11668027) following manufacturer’s protocols. Two days following transfection, antibiotic selection was performed with 2i/Lif media supplemented with  6 μg/ml Blasticidin (Thermo Fisher, A1113903) until resistant mESC colonies were evident. Resistant colonies were passaged once and then single-cell sorted on 96-well plates. 8 days following FACS, 96-well plates were duplicated and genotyping was performed via PCR amplification of genomic DNA and sanger sequencing. CuO array was next inserted 1) 100kb across the TAD boundary, 2) 100kb within the TAD boundary, 3) 400kb across the TAD boundary, or 4) 400kb across the TAD boundary (also see Figure 4). Similarly, cells were co-transfected with 1.6μg Cas9/sgRNA expressing plasmid and 0.8 μg purified 48-mer CuO donor DNA using 5 μl Lipofectamine 2000 (Thermo Fisher, 11668027) following manufacturer’s protocols. Two days following transfection, antibiotic selection was performed with 2i/Lif media supplemented with  200 μg/ml Geneticin (Gibco, 10131035) until resistant mESC colonies were evident. Resistant colonies were then passaged once and then single-cell sorted on 96-well plates. 8 days following FACS, 96-well plates were duplicated and genotyping was performed via PCR amplification of genomic DNA and sanger sequencing.

Rad21-degrons were generated through CRISPR/Cas9-mediated genome editing. To obtain homology donor, FLAG::FKBP^F36V^::P2A::PuroR cassette was first synthesized (GenScript) and PCR amplified with primers with flanking 50 bp homology arms to 3’ end of murine Rad21 coding sequence in a 200 µl reaction with Primestar GXL DNA polymerase. Correctly-sized amplicons were confirmed on an agarose gel and purified via ethanol precipitation method. 350,000 TRACK-IT parent cells were plated onto a 6-well plate the day before transfection. Cells were co-transfected with 1.6μg Cas9/sgRNA expressing plasmid and 0.8μg purified donor DNA using 5 μl Lipofectamine 2000 (Thermo Fisher, 11668027) following manufacturer’s protocols. Two days following transfection, antibiotic selection was performed with 2i/Lif media supplemented with 4 μg/ml Puromycin (Gibco, A1113803) until resistant mESC colonies were evident. Resistant colonies were then passaged once and then single-cell sorted on 96-well plates. 8 days following FACS, 96-well plates were duplicated and genotyping was performed via PCR amplification of genomic DNA and sanger sequencing.

**3.3 Sleeping Beauty transposition-mediated locus tiling**

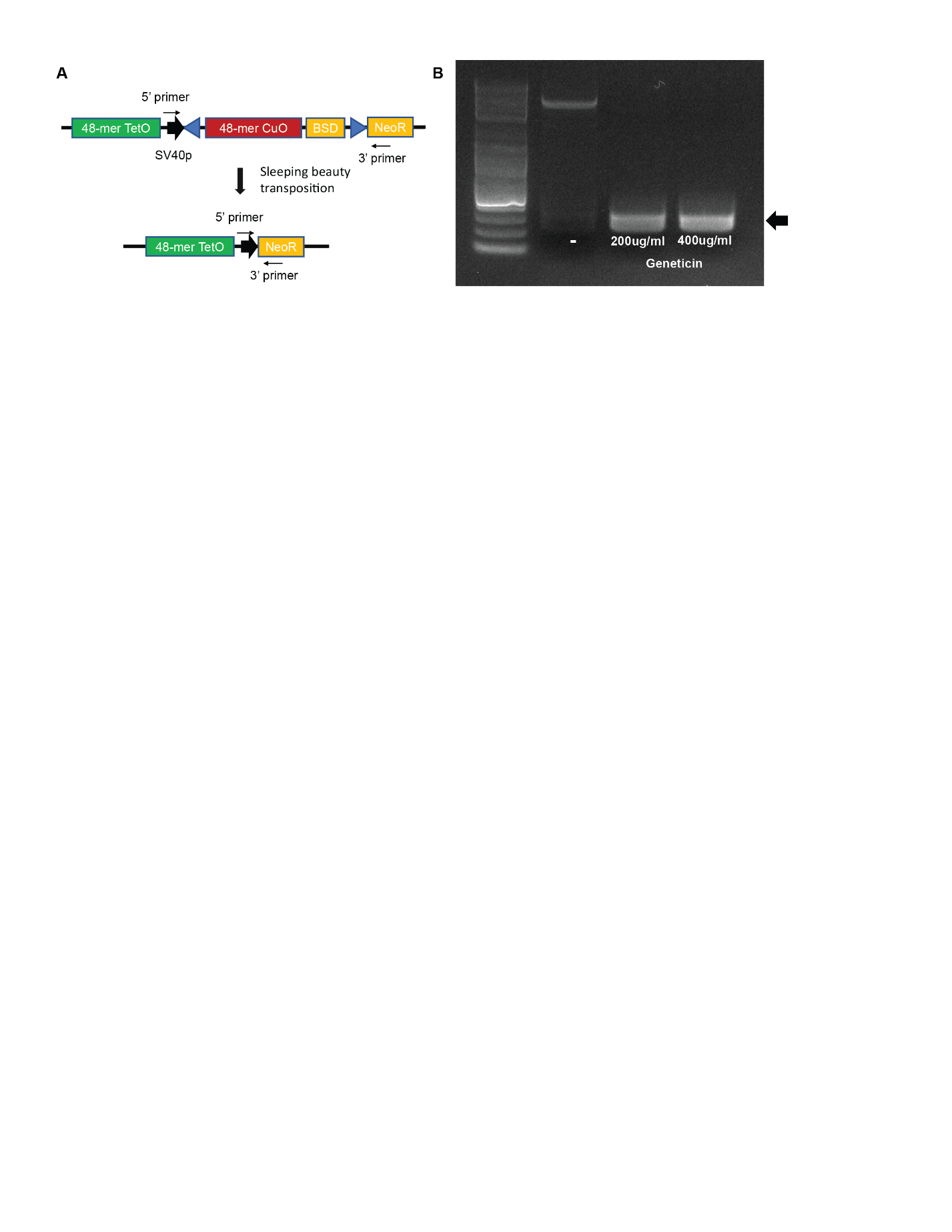

**Methods Fig. 3. Sleeping beauty transposition**. **(A)** Schematic of TRACK-IT locus and primers used to validate sleeping beauty transposition. **(B)** DNA gel electrophoresis showing post-transposition PCR products (black arrow) following sleeping beauty transfection, followed by no selection, 200 or 400 μg/ml Geneticin selection.

To remobilize CuO array-containing Sleeping Beauty transposon, 350,000 TRACK-IT parent cells were plated onto a 6-well plate the day before transfection. Cells were transfected with 2.5 μg SB100x expressing plasmid pCMV(CAT)T7-SB100 (a gift from Dr. Zsuzsanna Izsvak (Addgene plasmid # 34879 ; http://n2t.net/addgene:34879 ; RRID:Addgene_34879)) using 5 μl Lipofectamine 2000 (Thermo Fisher, 11668027) following manufacturer’s protocols. Transfected cells were supplemented with 0.3 μg/ml of cumate to prevent CymR binding to the CuO array within the transposon that may hinder Sleeping Beauty transposase activity. Two days following transfection, antibiotic selection for transposition was performed with 2i/Lif media supplemented with 200 μg/ml Geneticin (Gibco, 10131035) until resistant colonies were evident. Resistant colonies were then passaged once and then single-cell sorted on 96-well plates. PCR amplification with primers flanking the transposon were performed to confirm that resistant colonies indeed resulted from either 200 or 400 μg/ml Geneticin were indeed result of transposon remobilization (**Methods Fig. 3**).

8 Days following FACS, plate was duplicated using 0.25% Trypsin-EDTA (Gibco, 25300062) onto a glass-bottom 96-well plate (Cellvis, P96-1.5H-N) pre coated with 10 μg/ml human fibronectin (Milipore, FC010-5MG) and an uncoated 96-well plate with 2i/LIF media supplemented with 20% FBS and 20% DMSO to facilitate freezing. Next day, glass bottom 96-well plates were imaged and screened for local re-integration by measuring the 3-D distance between the TetO and CuO labels (for details see **4.4. high throughput imaging screens**).

**3.4 Transposon re-insertion site mapping via T7 in situ transcript sequencing (isT-seq)**

In order to rapidly map the integration sites of the mobilized CuO arrays we developed a protocol we call in situ Transcript sequencing (isT-seq) (**fig. S3A**), which used in situ transcription from the synthetic T7 promoter inside transposon, followed by RNA sequencing, to map integration sites.

Genomic DNA for candidate cells were purified using Quick-DNA Microprep Kit (Zymo Research, D3020) following manufacturer’s protocols with >500,000 input cells and 11 μl elution volume. Each in situ transcription reaction was prepared as follows: 10 μl purified genomic DNA, 2μl T7 RNA Polymerase Mix (NEB, E2050S), 1 μl murine RNAse inhibitor (NEB, M0314L), and 7μl milli-Q water were mixed and incubated at 37°C overnight. In situ transcripts were then purified using RNeasy Plus Micro Kit (Qiagen, 74034) following manufacturer’s protocols. RNA-seq libraries of in situ transcripts were prepared using Stranded Total RNA Prep kit (Illumina, 2004052s5) following manufacturer’s protocols.

Multiplexed library sequencing was performed on NovaSeq 6000 sequencing platform (150 bp paired-end reads). ~10 million reads for each clonal TRACK-IT lines ~25 million reads for each non-clonal lines (unsorted TRACK-IT post-mobilization cells and random sleeping beauty insertion controls) were generated following adaptor de-multiplexing.  Raw reads were quality checked with FastQC and selected for in situ transcripts by filtering for raw reads that align to sleeping beauty ITR sequence using BWA (*132*). Filtered in situ transcript reads were then aligned to the mouse genome (mm10) also using BWA.

Sleeping Beauty ITR-genome junctions were identified by a custom Python script, findJunctions (see **Code Availability**). In short, findJunctions takes in mapped sam files as inputs, and outputs the 1) genomic coordinates of uniquely mapped sleeping beauty ITR-genome junctions, 2) read counts, and 3) strand (+ for pre-jump orientation and - for flipped orientation) in bed file format. Each mapped reads are queried for the 3’ end of sleeping beauty ITR sequence (CTTCAACTGT) or its reverse complement and genomic coordinates and strand (+ for ITR sequence match, - for reverse complement) are returned.  Read counts were determined by the number of mapped reads that span the same ITR-genome junctions. For each clonal line, a single unique ITR-genome junction was unambiguously mapped with ~1000 read counts (**fig. S3B-D**). For ITR-genome junctions with higher read counts (>1000), a small number of variations in exact ITR-genome junction coordinates were found (+-5 bp) with very low read counts (<5) (**fig. S3C**), which may represent sequencing error. Insertion sites were further validated by Sleeping Beauty target sequence (TA). Coverage plots were generated with bamCoverage (*133*) and resulting bigwig (.bw) files were visualized on Integrated Genome Browser (*134*) or UCSC Genome Browser (*135*). (Note that while this workflow has been sufficient for mapping most *in situ* transcripts, it may be possible for insertions in highly repetitive sequences, initial filtering for ITR-sequence is insufficient for unique mapping to the genome. We found that in situ transcription from a known loci has mapped reads more than 5 kb downstream (**fig. S3B**), suggesting that in situ transcript-sequencing strategy that initially assembles the reads into contigs before filtering/mapping may be particularly suitable for mapping transposon insertions even in highly repetitive region of the genome.

We performed two additional control mapping experiments: 1) TRACK-IT transposon remobilized from plasmid (random transposon integration as opposed to a genomic reinsertion), and 2) negative control without any TRACK-IT transposons (**fig. S3E**). For the random integration control, we were able to identify >30,000 unique integration sites spread largely uniformly across the genome (**fig. S2C**), suggesting that in situ transcript sequencing can efficiently map a large number of heterogeneous integration events. Notably, not a single false-positive ITR-genome junction was identified for negative control (**fig. S3E**), demonstrating the remarkable accuracy of isT-seq. In comparison, PCR-based methods (ie splinkerette PCR, inverse PCR), often present with PCR-associated false-positive artifacts (*136*, *137*).

**4. Imaging and calibration**

***4.1 Optical setup***

Images were collected on a custom built confocal setup. Four solid state Obis lasers (Coherent), 405 nm 50mW, 488 nm 150 mW, 561 nm 150 mW, 647 nm 120 mW were combined using custom dichroics (Chroma) to an armored single-mode fiber (FO-2, Solamere). The fiber was attached to a Yokogawa spinning disk scan head, with a 50 um microlens array disk, dual camera ports, and the W1 Uniformizer calibrated for the Nikon Ti2 microscope body with a double stacked filter wheel, designed to reshape the illumination beam in the focal plane to a top-hat/square pattern. The Yokogawa scan head contained the following optics: a quadband dichroic to eliminate scattered light: zt405/488/561/640tpc (Chroma), dual filter wheels for both cameras with ET em band pass or long pass filters, AR coated: 460/50, 605/70, 700/75, 525/50 (Chroma), and a dichroic splitter for two-color imaging with T565lpxr and T640lpxr dichroics (Chroma). The T565lpxr was used for this study.  The scan head was attached to a Ti2 Nikon body (Nikon). The lower filter wheel was used in the empty position for confocal imaging (this wheel contains multichroic mirrors and filters which we use in widefield imaging on this setup). The upper filter wheel contains a ZT1064rdc-sp dichroic and an ET910sp filter (Chroma) which allowed a 980nm IR laser from a home-built autofocus system (Thorlabs) to be coupled to the sample. The construction and operation of the auto-focus is described in detail in the protocol paper, Mateo et al Nature Protocols 2021 (*138*). The Ti2 body was mounted with a precision 3D-stage (Ludl), with a 500 μm travel range piezo Z-stage. Microscope body and stage were enclosed in an environmental chamber to regulate temperature, humidity, and CO2 levels (Okolab).  Both camera ports used the 95% QE Fusion, scientific CMOS cameras (Hamamatsu). The cameras were aligned manually and residual offsets were computed by imaging beads before each experiment to finalize alignment.  The microscope system was controlled using the open-source software: storm-control/Hal4000, written in python and originally developed in the Zhuang Lab at Harvard University  (*113*).  Our branch of the software is available on Github (see Code Availability).

***4.2 Accelerated z-imaging and calibration of z-resolution***

In this section we describe our rationale for designing our rapid 3D scan approach and present some of the data which guided this design, in addition to reporting the approach selected. The Z-scan speed is limited by the speed and accuracy of the z-stage, which itself is affected by the mechanical precision of the stage, the mechanical inertia of the stage/sample combination, and the precision and accuracy of the control system. Z scans were performed using a piezo-z stage (see Optical Setup) under direct control of National Instruments DAQ-card (NI PCIe-6353). The DAQ provides synchronization of the stage movement with the camera exposure, to avoid catching frames in which the stage is scanning (something that is not always achieved when using software/CPU based timing). Piezo stages also provide rapid and precise movement for short scan range. We used the IR laser from our autofocus system to measure the position of the z-stage to test and validate the accuracy of our scan and its inertia. In this system, an IR laser is split into two beams, which are directed in a nearly parallel path to bounce off the coverglass and back to a sensitive high-speed CMOS camera. A slight tilt away from a perfect parallel results in a horizontal shift in the distance between the beams in the plane where they reach the IR camera that varies as a function of the path length (and thus the distance of the coverglass from both the IR cameras and imaging cameras), providing nanoscale accuracy - as described in earlier publications (*138*, *139*).

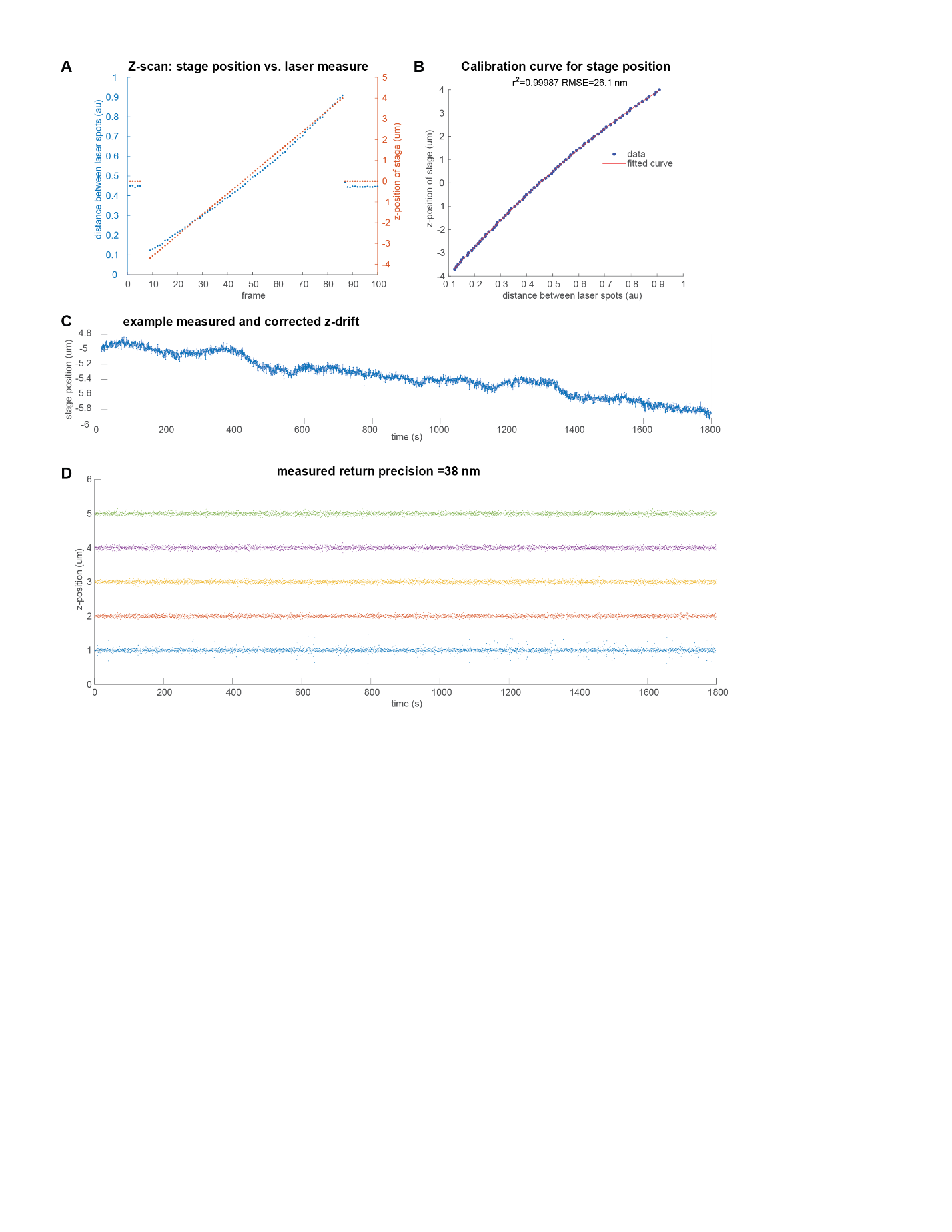

**Methods Fig. 4.** Calibration and validation of Z-scanning. **(A)** Plot of piezo-z stage position set by the software, in microns (orange), overlaid on plot of measured z-stage position, in terms of the distance between the reflected pair of IR laser beams incident on the IR focus-lock camera (blue), as functions of frame number.  **(B)** As in A, but plotting set-point in microns vs. measured distance. Red line shows a polynomial fit to the data, which is used in converting the measured z-positions to microns during fast imaging, bypassing the modest effects of stage inertia.  **(C)** Example of z-position data calibrated by direct measurement over a 30 minute movie.  **(D)** Example data showing the corrected positions of a 4 um scan, quantifying the return precision of the stage at each step.

To calibrate the accuracy of our stage and focus system we recorded a linear scan of 8 um in 100 nm steps, recording the separation of the reflected IR laser spots (**Methods Fig. 4A**). We observed a near linear scaling, (**Methods Fig. 4A**), which can be accurately fit with a second order polynomial with *r*^2^=0.9999 and a root mean square error of 26 nm (**Methods Fig. 4B**). We use this polynomial calibration to convert recorded stage position from the IR focus control into units of nanometers, facilitating tracking and correction of the modest stage drift and error in return-precision of the z-stage loaded with the sample (**Methods Fig. 4C**). When using a larger 1 um step per 100 ms, we still observed highly reproducible z-scan behavior (after z-drift correction) across the 30 min interval (compare **Methods Fig. 4C**-**D**), with a modest increase in variation of ~38 nm). These analyses demonstrated that the our z-stage is able to maintain the large 4 μm amplitude scan in 500 ms, and that stage inertia and settling time are only modestly impacting the accuracy and reproducibility of the scan. The modest increase in variation (12 nm) suggests further innovations would be needed to make deeper scans work with precision.

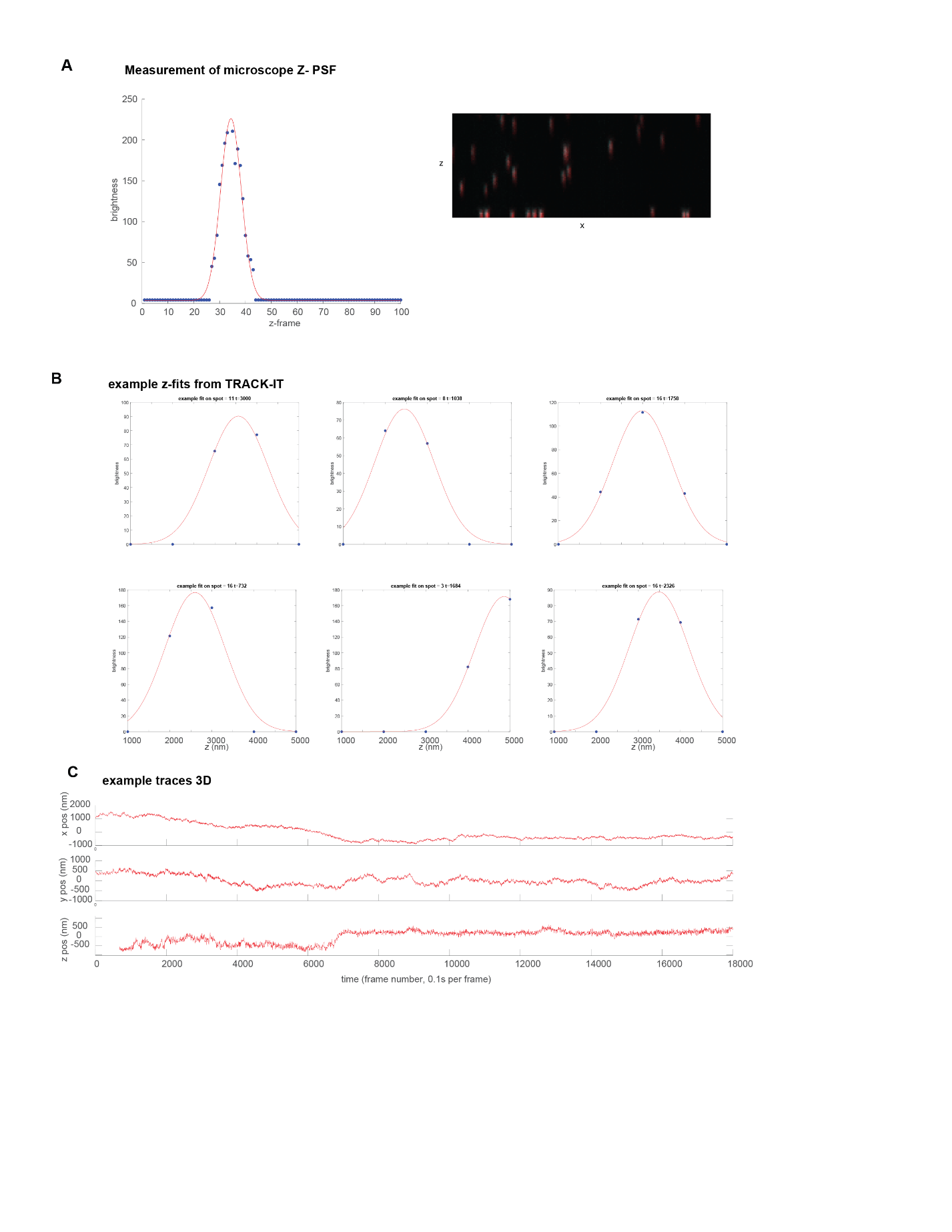

**Methods Fig. 5.**  Z fitting and calibration of axial PSF.  **(A)** Calibration of the axial (z) point spread function (PSF) of the microscope from an 8 micron scan of gel-embedded tetraspeck beads. Left - graph of the PSF from a single bead, the blue dots record brightness-per-frame. Red line shows a Gaussian fit to the data, used to approximate the PSF. Right - an example image showing an x-z projection of the scan.  **B.** Graphs showing example fits to fluorescent chromatin labels during track it, collected with the fast, sparse scan, fit to the PSF determined from (A). Only the height and z-position of the PSF were permitted to vary during the fit. The software prints fit graphs like these once per 1000 fits to facilitate curation of data quality.  **C.** Example of 3D traces from TRACK-it, showing x, y, and z, positions separately.  All graphs in nanometers.

To maximize our imaging speed, we sought to minimize the time required per z-scan while balancing spatial resolution and dynamic range. To determine our z-scan step size, we considered the trade-off between the number of data points in the axial fit, the size of the axial point-spread function (PSF), the motion blur in image acquisition, and the ultimate effect on the temporal resolution of the 3D trace. Fewer, larger steps naturally enables a faster z-scan, but also reduces the number of data points used in fitting the PSF to determine the axial position. We measured the axial PSF of our system in advance (**Methods Fig. 5A**), by imaging tetraspeck beads embedded in a 3D gel, using the approach described below (**Section 3.5 Chromatic calibration**). Having characterized the PSF, only the magnitude and axial (z) position of the PSF remain as free parameters, which can be determined as long as the spot signal is observed in at least two frames. This was achievable in our TRACK-IT data with 1 um step size  (**Methods Fig. 5B**) after we upgraded to brighter fluorophores and the higher quantum efficiency cameras. Notably we observed high quality fits to both background and signal when using a 1 um step size and a 4 um scan (**Methods Fig. 5B**), while simultaneously achieving a 3D stack in 0.5s. Shorter exposure times reduce photon-signal to electron-noise and larger, sparser steps provided insufficient constraint in fitting the axial PSF. Thus we selected the 1 um step size 5 um scan for our rapid z-scan, due to its high performance in this assay.

The resulting z-trajectories (**Methods Fig. 5C**) show considerably smoother paths than we were able to achieve in earlier iterations of our imaging using a previous generation of cameras and published cell lines, (generously provided by Dr. O. Weiner, UCSF).  As the axial position of a moving object is expected to be correlated with its previous position, whereas a noise-dominated measurement is not, we take these smooth trajectories as further evidence of the efficiency of this scan.

***4.3 Live microscopy***

Cells were plated onto glass-bottom 96-well plate (Cellvis, P96-1.5H-N) pre-coated with 10 μg/ml human fibronectin (Milipore, FC010-5MG) at a density of 50,000 cells/well the day before imaging. The next day, CymR-2xHaloTag was conjugated with 100 nM JF549 (Promega, GA1110) diluted in culture media for 2 hours and then media was replaced with imaging media (2i/LIF media supplemented with 1:100 v/v Prolong Live Antifade Reagent (Thermo Fisher, P36975), 50 µg/mL ascorbic acid, and 2.5 µg/mL doxycycline (Selleck Chemicals, S4163)) for 2 hours prior to imaging to reduce photobleaching. For Rad21 depletion experiments, cells were incubated with imaging media additionally supplemented with 500 nM dTag-13 for 2 hours.

Imaging was performed on our live-imaging microscopy setup with a custom enclosure (Okolab) maintained at 37°C and supplied with 5% CO_2_ pre-mixed air (Praxair). Automated imaging was performed on 2 FOV per well with 2 wells per condition (total of 8 FOV for a single imaging session). Unless stated otherwise, accelerated z-imaging method was used for 30 min/FOV (18,000 frames total). Two biological replicates, as defined by separate instances of cells cultured and plated for imaging, were performed for each experiment. For each cell line, we recovered 300-3000 traces from as many cells, as shown in **fig. S1B**.

***4.4 High-throughput imaging screens***

FACS-sorted plates were duplicated using 0.25% Trypsin-EDTA (Gibco, 25300062) onto a glass-bottom 96-well plate (Cellvis, P96-1.5H-N) pre coated with 10 μg/ml human fibronectin (Milipore, FC010-5MG) and an uncoated 96-well plate with 2i/LIF media supplemented with 20% FBS and 20% DMSO to facilitate freezing. The next day, CymR-2xHaloTag was conjugated with 100 nM JF549 (Promega, GA1110) diluted in culture media for 2 hours and then replaced with culture media for at least 1 hours. Prior to imaging, the plates were carefully turned over to avoid spilling the culture media and 2.5 μl of Zeiss Immersion Oil (Zeiss, 444970-9010-000) was applied on the bottom of each well, allowing oil-immersion lenses to be used for imaging the entire 96-well plates.

Imaging was performed on our live-imaging microscopy setup with a custom enclosure (Okolab) maintained at 37°C and supplied with 5% CO_2_ pre-mixed air (Praxair). Automated imaging was performed on 4 FOV/well with custom software. To minimize stage travel, wells were imaged in a zigzag fashion (A1~12, then to B12~1 and C1~12, and so on).

***4.5 Chromatic calibration***

After each imaging session, refractive index-matched (*n* = 1.36) 3D gel-embedded beads were imaged to measure chromatic aberration between the two channels. Refractive index-matched gel was made by diluting 0.1 μm Tetraspeck beads (Thermo Fisher, T7279) at 1:20 ratio in 2% w/v agarose and 7.4% w/v sucrose solution which was then solidified on a single well of glass bottom 96-well plate before each imaging session. Bead-gels were made fresh for each imaging session to minimize bead fluorescence loss and changes in gel consistency. 4 z-stack images, 0.1μm/frame across the 10 μm z-section centered on the z-plane of the corresponding experiment, were taken as reference for chromatic calibration. FOVs for each image were chosen for dense, uniform distribution of the beads.

**5. Image Analysis**

Our ProcessTraces pipeline was written from the ground up, guided by approaches used in previous work. Due to the fact our source data was up to several orders of magnitude larger than in prior work, computational efficiency and robustness to several rare potential sources of error were essential in the design. The large data size also makes it impractical to rely on manual curation steps used in some prior work. Here we describe the approach, noting that several steps in the pipeline are indeed more complicated than may be expected, which reflect additions and expansions added to make the system either more robust to certain rare types of error (described below), more computational efficient, or both. The code is written in Matlab(™), Python, open source and freely available, as described in the Code Availability statement.

***5.1 Chromatic correction***

Our chromatic correction software runs a matlab GUI and is available open source from our Github page (see **Code Availability**). Here we describe the steps performed for chromatic correction.  As the two color channels were collected simultaneously on two separate cameras (see above), the first step of the chromatic correction was fine-scale registration of the two cameras using the bead images.  Prior to correlation alignment, we also correct for the left-right reflections created by the dichroic mirror.  We then used an image-correlation approach for registration, which maximized the cross-correlation between the maximum-intensity-projection images to compute the appropriate x-y alignment, as described previously for fiducial alignment (*138*). Briefly, cross-correlation was computed for a range of x-y translations and rotations, using a multiscale approach that scans a broader range of parameters with a down-sampled image and then finer range of parameters for an upscaled section of the image. This provided robust visual alignment removing rotational and translational differences between the mounted cameras.  Next we performed 2D spot fitting of the bead images. Raw movies in .dax format were analyzed using the storm-analysis package in python, developed by the Zhuang lab (*140*) for long movies of high density single particle localizations, and used extensively in prior works. This software is open-source and thoroughly documented at *readthedocs* https://storm-analysis.readthedocs.io/en/latest/. Fits were computed using the following parameters:

```
<?xml version="1.0" encoding="ISO-8859-1"?>
<settings>
  <start_frame type="int">0</start_frame>
  <max_frame type="int">-1</max_frame>
  <model type="string">2d</model>
  <background_sigma type="float">8.0</background_sigma>
  <camera_gain type="float">1.0</camera_gain>
  <camera_offset type="float">100.0</camera_offset>
  <find_max_radius type="int">10</find_max_radius>
  <foreground_sigma type="float">2.0</foreground_sigma>
  <iterations type="int">20</iterations>
  <pixel_size type="float">108.0</pixel_size>
    <threshold type="float">35</threshold>
<!-- Threshold in units of sigma, as in "3 sigma event." -->
  <sigma type="float">1.3</sigma>
  <radius type="float">0.0</radius>
  <do_zfit type="int">0</do_zfit>
</settings>
```

The 2D positions of the fitted points were registered using the horizontal, vertical, and rotational translations computed. We next linked images of single beads across the Z-stack based on their *x,y* positions, separating beads that ended up vertically stacked using local-maxima detection.  We note that the storm-analysis tool does include 3D fitting modes, but note these are based on PSF engineering (cylindrical aberration), in which the axial position is imputed directly from the x-y position.  The resulting Z-linked data were then fit to an axial PSF (see example in **Methods Fig. 5A** and **fig. S5**) to create a 3D spot position to floating point precision (as opposed to pixel precision). Points from both color channels were paired by minimizing the total offset among all paired points within a 600 nm maximum cut-off distance.  We then used the paired spots from both cameras/color-channels to compute a 2nd order polynomial 3D map between the channels.  **fig. S5E**, shows the vector map of the difference in position between the two colors channels, before and after correction. The general shape of the 3D polynomial can be seen in the pattern on the uncorrected map (**fig. S5D**).  The Plan Apo chromatic correcting objective left a residual chromatic error of 177 nm in xy (296 nm in 3D).  After correction the paired points had an average offset of 13 nm (xy), 23 nm in 3D (**fig. S5E**).  The resulting polynomial function and camera registration file were saved alongside the data, to be used in chromatic correction of the cellular data in the following steps.

***5.2 Fitting spots***

Raw movies of live cells carrying TRACK-IT labels were recorded in .dax format were analyzed using daostorm_3D from storm-analysis software (*140*), a python based program developed by the Zhuang lab for long movies of high density single particle localizations, and used extensively in prior works. This software is open-source and highly documented <https://storm-analysis.readthedocs.io/en/latest/>. Importantly, this software allows resolution of two overlapped emitters that produce a unimodal PSF on the detector (below the classical Abbe limit), based on the perturbation to PSF shape, as described in prior publications (*140*, *141*).  This is helpful in improving tracking of sister chromatids from cells in the G2 stage of the cell-cycle, as described below.  Significance thresholds were calibrated by hand for each movie using the middle frame. Thresholds ranged from 10 to 30 sigma, depending on the cell line. Threshold is measured in units of sigma, a common metric of the confidence interval for the probability that background fluctuations are mistaken as a true signal, see (*140*) for further details. To ensure robust traces, we selected permissive thresholds to maximize the number of detected events. The spurious localizations sometimes captured by these permissive thresholds are filtered out in subsequent steps of the data analysis, as described below. The fit files were saved as hdf5 tables.

 Fitting parameters used were otherwise as follows (also provided as an xml file, see fitPars_C1.xml).

```
<?xml version="1.0" encoding="ISO-8859-1"?>
<settings>
  <start_frame type="int">0</start_frame>
  <max_frame type="int">-1</max_frame>
  <model type="string">2d</model>
  <background_sigma type="float">8.0</background_sigma>
    <camera_gain type="float">1.0</camera_gain>
  <camera_offset type="float">100.0</camera_offset>
  <find_max_radius type="int">10</find_max_radius>
   <foreground_sigma type="float">2.0</foreground_sigma>
  <iterations type="int">20</iterations>
  <pixel_size type="float">108.0</pixel_size>
  <threshold type="float">24</threshold>
  <sigma type="float">1.3</sigma>
  <descriptor type="string">2</descriptor>
  <radius type="float">0.0</radius>
  <do_zfit type="int">0</do_zfit>
</settings>
```

***5.3 Identification of seed points***

The next step was to identify small windows within the large field of view that contained sufficiently complete traces for tracking. For this, first the spot localizations per frame from daostorm_3D were imported from the hdf5 format, and transformed using the camera alignment and chromatic correction functions computed using 3D images of beads embedded in 3D gels as described above.  We then identified localizations that spatially clustered across the frames by binning the *x,y*, spot positions across all frames and searching for dense connected bins, using a minimum occupancy threshold of spots per bin to define connected regions, and a final minimum threshold of total spots within all bins of each connected region to restrict the analysis to sufficiently complete traces.  The centroid of each connected block of bins was recorded as the seed point for that color. An identical approach was applied to both color channels, and a further filter kept only those seed points for which a matched seed point in the same cell was detected.  This approach is much faster than other nearest neighbor clustering methods, an important factor when the individual movies contain many millions of recorded spots.  Binning parameters were first calibrated by hand, we found the same parameters worked well across our data set. These parameters were 756x756 nm bins (7-fold reduction relative to our pixel size), a minimum of 50 spots per bin and a minimum of 400 spots per trace.

***5.4 Linking localizations into 4D traces***

To assemble 4D traces for the detected spot pairs, we developed a layered approach that starts with a coarse assembly and refined the trace later with the single frame data. This approach prevented inaccurate traces arising due to the combination of stray spurious detection events and occasional missing data (e.g. where spots drifted out of the limited z-scan range or were below threshold required for accurate localization).  To compute coarse trajectories, we first binned the data from 18,000 frames into 50 blocks (360 frames each), and computed a centroid position for all localizations in each block in a similar approach to that used with seed points (see section 5.3), using a 432x432nm binning and minimum of 10 observations per bin to define connected regions. Then, all seed points were matched to one of these new centroids using an optimization strategy that minimizes the total distance of linked elements. This  image-wide optimization approach prevented errors that would arise in a closest-point scheme when one trace starts close to where another points average position, due to substantial spot motion (largely cell migration) that occurs during long movies.  These parameters were chosen by manual optimization and visual inspection of the assigned traces overlaid on the raw data. We found the same parameters performed well for all of our data sets.  The output of this first layer is a 50-step coarse trajectory.  This coarse trajectory was filtered to remove jumps (that arise from on rare occasions due to clusters of localizations that appear, for example by autofluorescent cellular debris). The jump filter uses a sliding window of 6 frames and removes steps that have more than 3 fold the median step size or an absolute step size of more than 7 pixels (756 nm). This adaptive filter helped address the substantial variation in the degree of motion relative to the camera reference frame exhibited by the different labeled loci in different cells.  We then refined the trace further by considering a 35 pixel radius around the centroid of each spot, and averaging the *xy* positions in each time block (360 frames) for only those points within 6 pixels of the coarse moving average.  This step helps remove the effect of outlier points on the moving average in the binning approach.  The restricted region of interest and the temporal binning makes this step fast (even looping over all spots) whereas the averaging positions specifically of localizations in the vicinity of the coarse trajectory improves the resolution of the trace and reduces the pull of occasional outliers. Should the previous jump filter erroneously lose part of the path, this step will restore it as it is based on the xy data of each localization - provided it is still in a connected path. This intermediate linking step produces a refined version of the moving-average trajectory. We then apply a smooth interpolation to the moving average trajectory, to allow the prediction of the expected location of the trace in each frame.  We then loop over all of the spots, preceding one frame at a time, adding to the trace the nearest localization. This simple nearest next spot linking algorithm works well on most of our data by itself, but occasionally jumps to stray distal points in frames where a spot was not recorded. To avoid this, in this final frame-by-frame linking step, the algorithm rejects adding to the trace any points that are more than 540 nm (5 pixels) distal from the interpolated moving-average trajectory, or more than 432 nm (4 pixels) away from the last linked point.  In this way, the full time-course of the data provides a guiding reference to interpret if individual steps are real, or are just transient stray points that might pull the trace off-track. This filtering/threshold approach successfully excluded the rare cases where background signal was detected during an interval in which the tracked spots had disappeared (by diffusing momentarily out of the z-scan range, for example). To facilitate identification of sister chromatids, at each step we take up to 2 closest points within the 35 pixel radius. In G1 cells this second trace list is largely blank, with rare (<3%) of steps having background localizations recorded, which typically lack any spatially clustered organization.  In G2 cells, we observe two consistent doublets in most frames. Due to the high temporal resolution of the data for the spot linking, the segregation of the sister chromatids was generally unambiguous - though downsampling this data from 0.1s steps to 2s steps, or 20s steps, removes this clarity. Accordingly, short intervals with missed detection events (<10 s) in rapidly moving cells sometimes resulted in misassignment of the sisters, where cellular motion positioned a label's sister chromatid closer to where it had last been seen than the true label. These cases can be disambiguated by comparing the traces of both sisters, and referencing them to the other color channel, as described in the next section.

***5.5 Identification and pairing of sister chromatids in 3D***

The next step uses the trace data from the previous step to identify if cells are in G1 or G2, and cross-references the data from both channels and both sisters in order to catch and correct rare errors (like that alluded to above), in which sister assignments are confused.  This step first computes the fraction of frames in which a doublet of spots was detected compared to only single spots. We classified cells with over 5% doublet detection as G2, which corresponded to 60-70% of cells, consistent with our measurements of the fraction of G2 vs G1 cells in the population as measured by fluorescent activated cell sorting (FACS). For G2, traces were then processed as follows. First we removed large-scale correlated motion arising from cell movement which occasionally confused assignment of sisters when confronted with occasional gaps in data. We accomplished this as follows. First, we computed a 3D moving average path through the data in each of the 4 traces (two sisters, two colors), where z-is only tracked at present to the nearest frame resolution (this will be refined following sister designation, disambiguation of sisters before the refined z-analysis reduced some sources of errors in the z-analysis derived from converging sisters). The moving average calculation used a window size of 50, and a generous minimum of two localizations detected in the window to buffer against rare spurious detections. The resulting data was averaged across the 4 traces (both sisters, both colors) and was smoothed with an outlier resistant version of a least squares filter to a second-degree polynomial (rloess algorithm) with a window size of 90% of the trace length. This smooth trace was subtracted from all the individual traces which removed the persistent directional motion which tended to complicate accurate assignment of sisters. These same values are added back to the motion path after disambiguation, so this adjustment does not ultimately affect the motion traces themselves, only their assignments between sisters.  In this motion-corrected reference frame, we then repeated the nearest neighbor per frame assignments simultaneously for both sisters, minimizing the sum of the displacements of both sisters and requiring a maximum distance to the last observed point of 8 pixels (this is larger than we ever observe between consecutive frames, but traces with gaps of 30s or more do achieve such gaps on events). We then computed an additional reference trace for each of the 4 paths using a linear interpolation, allowing an estimate of the approximate position of each of the 4 spots in every frame, for the purpose of assigning sisters. Note this interpolated data is only used as a fiducial reference line for sister assignment, missing observations in the saved traces remain populated by NaNs. We then compute the distance for all frames among all four observations per frame. We pair each sister in the first color channel with the sister in the second color channel that is on average closest to it.  We then identify all instances where the two sisters come with-in 300 nm of one another (approaching the diffraction limited resolution of the PSF) as potential intervals where the sister identities may be swapped within a path. Between any two such points, (or any one such point and the start or end of the trace), we compute the distances among all 4 points across all observed frames.  If the intra-sister and intra-color distances across the entire trace would be shorter by accepting a swap of sister identities in this interval, we accept the swap. This is motivated by the assumption that we expect more coherent motion from two loci that are linked in cis or linked as sisters than we expect from a trace that is a mashup that bounces between sister assignments.

We note this is related in concept to the analysis approach we recently published describing a method for back-propagation of the identities of individual chromatin loci imaged in the same color and decoded by end-point imaging (*142*).  More details on the principle can be found in this publication (*142*).

***5.6 Fine fitting of the Z-position***

We next refine the accuracy of the z-position by fitting the axial PSF for each localization for each sister in each color. As discussed in Section 4.2 and illustrated in **Methods Fig. 4**, this is accomplished using the pre-computed axial PSF for the microscope, determined from bead imaging, allowing only the axial position and the overall height over background of the Gaussian approximation of the PSF to vary. The fits are applied to the spot brightness computed by daostorm_3D in the spot-fitting step, and propagated along with the x,y position and z-frame data in the assembly of 3D traces and assignment of sisters. The background value is set to be equal to the dimmest pixel in the cropped region around the fluorescent signal that was recorded in the z-scan.  In general we found most spots to show good agreement with the pre-computed PSF.  In instances where the 75% confidence interval for the centroid was larger than the step size, we rejected the fit and recorded a NaN as the z-position.  For further validation, every 1000s fit was displayed for manual validation, examples of these appear in **Methods Fig. 3**.

***5.7 Merging traces split or duplicated due to seed point errors***

occasionally a single fast moving pair of spots is mis-annotated as two separate seed points by the approach above, resulting in a split of data (e.g. the first 9,000 frames are assigned as one trace and the second 9,000 frames of the same spot pair are assigned as a separate trace). To correct these rare errors, a subsequent step compares all traces. If two traces have perfectly complementary observation windows, such that the total number of shared frames is less than 1% of the total number of unique frames in the two traces, and the two trace centroids are within 30 pixels of one-another, they are merged. We did not observe any discontinuities between traces in the manual inspection of the traces automatically selected for merging.

***5.8 Computing 3D distances***

From the 3D *x,y,z* coordinates of each point, we then compute the 3D (and *xy* 2D) distances between both colored probes. At this point in the analysis we still track all of the data per camera frame (at 0.1s) resolution through the *z*-stack, with the fitted *z*-positions applied to all the non-empty positions for each block of 5 frames in the stack.  For the 3D distance, we compute the average *xy* position across the *z*-stack in each color channel and report the distance between these 3D points as the 3D distance. We note in cases where the two different labeled points reside at axial positions separated by > 1 um, there is an additional 0.1s lag in the observation of these points. The average step size between 0.1 s separated points was ~20 nm, the average xy step size between 0.5s separated points was ~30 nm. This is on the scale of the amount of expected by the Rouse polymer model with negligible measurement error (which expects the mean squared distance change between steps separated by time *dt* should scale as *dt*^1/2^), indicating a modest motion blur trade-off in computing the 3D distance.

**6. Trajectory analysis**

Code implementing the following analyses is included in our Github repository for this paper, see Code Availability.

***6.1 Photostability analysis***

In order to compare across published datasets in which brightness data was not reported, as well as to focus on the most important consequence of photobleaching for these experiments – the presence/absence of detected spots amidst the background signal, rather than photon counts – we computed how the number of total spots detected per frame varied across the length of the movie as a meaningful proxy of bleaching.  We started from the matrix of 3D distances (n_cells x t_frames), took the top 100 most complete traces, and computed the fraction of traces which contained data at each frame. Traces for individual cells may have missing data in any given frame for a variety of reasons, including poor fitting, diffusing outside the imaging area, or photobleaching. Of these, only photobleaching is expected to systematically increase over time, with  later frames in the movies systematically having fewer observations. This effect can be clearly seen in published data in the steady fall off of the observed fraction as the movie continues, and several of these publications comment directly upon the role of bleaching in determining the maximum length of the imaging time (*27*, *28*, *30*). We restrict the analysis to the top 100 most complete traces as the different studies vary in the use of arbitrary thresholds on the number of data points required to be included as a trace, and as excessively short traces may give a negative bias to the estimates of photostability. We required 100 traces to have a reasonable statistical sampling. This approach of computing the impact of photobleaching reveals the key functional consequence of bleaching on data quality (namely how it limits movie duration, since after a certain interval few traces are left) and bypasses issues about how absolute brightness was computed, how it varies across the field of view, etc.  A typical photostability trace from one of our cell lines is shown in **Fig. 2J**. Photostability traces for the remaining lines are shown in **fig. S6**.

***6.2 step-size vs. dynamic range***

For all traces, we computed the median step-size between adjacent frames (*F*), and compared it to the interquartile range (25th percentile to 75th percentile distances) of the individual trace (*D*).  We computed the median F and median D for each experiment, as well as the median of the F to D ratio (where the median is taken after computing the ratio of the single-trace F and single-trace D).  We applied this procedure to all the published data and all the TRACK-IT data.  For the comparison shown in **Fig. 2I**, we averaged the results from all experiments within a single paper for brevity. These underlying experiment-by-experiment results are shown in **fig. S1B** for each of the 40 published experimental datasets from the 5 comparison publications, along side the 30 individual datasets from this paper.

***6.3 Mean-square change in displacement analysis***

From the matrix of 3D distances, we computed the average change in the squared 3D distance for 500 logarithmically spaced time intervals between 0.5s and 1800s, for each individual cell, averaging over different lags (i.e. all data separated by x seconds). Downsampling the total number of lags (instead of using all 3,600) accelerated computation without an observable effect on mean squared change in distance. We used logarithmic spacing in downsampling the data as MSCDs are plotted on log-log scales for the ease of interpreting powerlaw scaling, the logarithmic spacing provides uniform spacing of the data points.  We report the median of the single-cell MSCD curves as the population MSCD.

For transparency, we report these raw MSCD curves, rather than showing localization error corrected curves in the main text (as several recent publications have elected (*29*, *30*, *57*)). We note that our uncorrected MCSD log-log plots show linear scaling of the MSCD from the start of the plot, as opposed to showing a flat shoulder that gets steeper as expected when localization error is large enough to confound the curve (*143*, *144*). This linear scaling in the TRACK-IT data implies our localization error is substantially smaller than in recent studies relying on error-corrected plots, and suggests minimal advantage from further editing the raw data.

This in turn bypasses the uncertainties associated with the correcting motion errors without ground truth fiducial signals with which to unambiguously quantify this error. For example, the signal-to-noise ratio of the fluorescent intensity of the label compared to the background of unbound fluorophores and dark current in the camera pixels results in some uncertainty in the localization precision, but one that can vary experiment to experiment and even cell-to-cell. Most prior publications do not include a quantification of the contribution of motion blur during image acquisition to localization error, and this value may differ between experiments, depending on how the average speed of the fluorescent targets changes. These examples highlight the challenges and limitations to interpretation introduced in “error-correction” of MSCD. Notably, these challenges have been substantially ameliorated by the recent development of a data-driven Bayesian approach for estimating the MSCD and the associated localization error from the 3D trajectory information (*30*). In estimating localization error on an experiment-by-experiment basis, this approach highlights how variable such errors can be across experiments (192-308 nm in that study). However, the Bayesian approach is constructed on the prior assumption that the data come from a specified underlying process such as the random walk of a Rouse Polymer, (specified prior to the fit). If this prior is not a good description of the data, the underlying error assessment will also be less accurate. We elected to bypass these concerns by plotting the measured mean square change in displacement.

To quantify the error associated with the MSCD measurements, we plot the 95% confidence interval of the population-average MSCD by bootstrapping over the individual MSCDs per trace that we average. If we instead plot the mean-square error associated with the total number of observations (across all time points and all traces for each lag), the error bars become invisibly small due to the large number of total observations in our study.

To facilitate interpretation of the MSCD data, we plotted a power law with a scaling coefficient = 0.4, the predicted scaling of a crumpled polymer (*77*, *78*), along with the scaling expected for the simple Rouse polymer = 0.5, shown as a reference in prior works (*30*, *57*).

***6.4 Passage Time***

For each dataset, we identify first the typical 3D distance between the pair in question in the current conditions by computing the median 3D distance between the spot loci, which we refer to as *d_start* for the search event analysis. We then scan through each distance trace, and identify all time points in which the observation points were within the interval *d_start* to *d_start* + 50 nm, and all the intervals at which the observation points were within the interval 0 to 0 + 50 nm.  When exploring alternate contact thresholds (e.g. 100 nm) the calculation was performed in the same manner except for using this alternate distance in the identification of the start and stop intervals.  For every *d_start*, we computed the amount of time between when the trace first left this interval to the time when it first entered the contact interval, and recorded this event as the search event.  The effect of changing contact threshold is shown in **fig. S11**.  Events that leave the starting interval and return to the starting interval again without having reached the contact interval are not counted as passage events.  The effect of censoring on estimating the median and distribution of passage times was corrected using the Kaplan-Meier estimator as described below. The 95% confidence interval on the estimate also computed Greenwood’s formula for the Kaplan-Meier estimator, as described below.

***6.5 Search frequency***

Search frequency was computed by first identifying search events, as described below (see **6.6 Search Time),** and recording the total number of events per hour of observation across all traces.  Uncertainty in the search time was computed by bootstrapping. For this we resampled among our total N traces a new set of N traces, drawn at random with replacement, so the new list may include some traces twice and some not all. We then recomputed for 100 such random draws the search frequency.  For errorbars, we plot the confidence interval from the 5th percentile to the 95th percentile from these resamplings.

***6.6 Search Time***

We defined the search time as the amount of time it takes any pair of observation points to travel from their median distance to contact. We used the same 50 nm windows to define the distal and proximal states as with the passage time. The search time differs from the passage time as it is not conditional on the trajectory not returning to the starting state. Instead, in the interval before any contact event (or between any two contact events) we identify all the times at which the 3D distance was within the distal window (between the median distance and the median distance + 50 nm).  Among these times we selected one at random, and pooled all of the selections from all contact intervals as the distribution of search times. The random selection process avoids biasing the data in the search time with the single contact events that were preceded by long intervals of no-contact yet where the domain spent a substantial fraction of time at the median distance.  The effect of censoring on estimating the median and distribution of passage times was corrected using the Kaplan-Meier estimator as described below. Search times that began before the start of the observation period or ended after the observation window (i.e. never reached contact) were considered to be denoted as right censored events for this calculation. Search times that completed in only 1 frame were denoted as left censored, as it is possible these events took less than 1 frame. We note single frame search times were an extremely rare event when using a 50 nm threshold, but one that becomes more common with larger thresholds. The 95% confidence interval on the estimate also computed Greenwood’s formula for the Kaplan-Meier estimator, as described below.

***6.7 Contact duration***

Contact duration was computed from the average number of consecutive windows that the trace remained below the indicated threshold. The effect of changing contact threshold is shown in **fig. S11**. Since we plot the average contact duration we plot the standard error of the mean as a measure of the uncertainty, (standard deviation of square root of the number of observations). The standard deviation and the mean were both computed using the Kaplan-Meier corrected data. Where the search time tended to be “right-censored” - meaning that longer searches tended to be truncated by the finite size of the search window, the contact duration data were primarily “left-censored”, with some contact events lasting only 1 frame (0.5 s). We denoted all contacts of 1 frame as left-censored data in computing the Kaplan Meier correction. Contact events that began before the start of the movie or persisted after the end of the movie were considered right-censored. We select the mean rather than the median as the median is restricted to integer values of the frame number (half-second intervals), whereas the mean uncover subtle yet statistically significant differences among the thousands of contact events observed in our different conditions, providing more information in this plot. We used the Cox proportional hazards model to compute the statistical significance of the difference in the contact durations, using the Matlab™ function, fitcox. This is a widely used approach for comparing lifetime distributions (*145*–*147*).

***6.8 Correction for censoring in time measurements***

In measuring the time spent in different processes (contact duration, search time) during the 30 minute observation window, naturally only events that occurred in less than 30 minutes can be observed. Events that started before the observation window began, or which completed after the 30 minute window ended, or both, go undetected. This leads to a systematic underestimation of the distribution of times, which is more acute for slower processes. To understand this process in the context of polymer diffusion, we conduct simulations of a Rouse polymer for 100,000 timesteps, and then examined the observed median search time for observations windows of: 100, 1000, 10,000, or 100,000 timesteps, as a function of monomer spacing from 1 to 100,000 monomers. Notably, the search time dependence for the Rouse polymer can be computed analytically, and scales with the second power of the interlocus spacing. As expected, for data from the largest sampling window of 100,000 steps, we recover this expected scaling (see **fig. S9B**). Truncating the data to shorter sampling windows lead to progressive loss of data to estimate median passage time from in an interlocus dependent manner, and a systematic reduction in median search time of the more distal loci (**fig. S9B**). The former effect leads to the appearance of a noisy and flatter pattern of search times for distal loci pairs (with complete loss of detected search events for the most separated pairs). The latter effect leads to a re-scaling of the power-law dependence of the search time, even while the confidence intervals remain tight and the trend of distance remains relatively clear.

The time truncation effect is similar to a well known effect in estimating lifetime distributions (e.g. from lightbulbs, or clinical statistics) (*148*). In its classic construction, lifetime measurements of a population of, say, lightbulbs, are truncated by the fact some bulbs had not yet burned out by the end of the study, and some bulbs had already been used for an unknown amount of time before the study started. Similarly, some search events had started (dipped below the average distance) before the end of the observation window but not ended by the end of the window (not reached proximity or returned to average distance). Other searches started before the beginning of the observation window (the points were already within average distance at the start), and then reached proximity during the window. For a raw search time analysis, we exclude these events and focus only on the complete passages. However, points with especially slow search times are more likely to exhibit what few searches they did complete if they were already below average at the start, and are less likely than fast searches to complete any search they did begin during the observation window. So these incomplete searches still hold information we can use to better estimate the true distribution, much like the incomplete lightbulb data. We record these as “censored” events, and use the Kaplan-Meier estimator

 
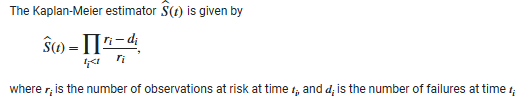

We compute the 95% confidence interval using Greenwood’s formula, an approximation for the variance of the Kaplan-Meier estimator,

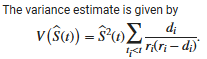

These computations are implemented in Matlab(™) R2023b, using the function *ecdf()*.

We validated this approach on our simulated data of the Rouse polymer, before seeking to apply it to our experimental data. As expected, the correction makes little difference to the shorter search times observed among proximal points, while for more distal pairs and shorter observation windows the correction substantially increases the computed median search. With the corrected values, we recover the expected exponent *a* = 2.0 even with a 10 fold shorter observation window. With yet shorter observation periods, the ability to correct the longest searches saturates, but the underestimation bias is  still reduced. The residual bias still leaves a scaling coefficient less than the expected *a* = 2.0, but provides a more accurate picture than without the correction bias.

To improve our ability to measure and correct for long search events, we explored an adaptive thresholding approach, which we also benchmarked on simulations. For multi-megabase separated elements, we expect to only observe the very fastest of the searches. Rather than restrict the analysis only to those events that successfully transition all the way from distances of up to a couple microns to 50 nm contact threshold, we can relax the threshold and see how long it took other traces to transition to 200 or 500 nm. This dramatically increases our pool of events, and allows us to examine how increase in genomic separation affects the scaling of the search time on the megabase scale (see **fig. S11**).

In contrast to other time estimation processes where the Kaplan-Meier correction can adjust for the censor bias, the search time estimate for long searches can also be informed by exploring the contact threshold. By relaxing the contact threshold, we can quantify the search time for searches that almost made it to contact. The search time scaling is not substantially affected by the choice of contact threshold (as we can see from simulation **fig. S11**).

***6.9 Cell Cycle analysis***

The effect of cell cycle state on the metrics described above are plotted in **fig. S10**. Cell cycle state was determined by analysis of the presence or absence of sister chromatids in the movies of each cell, as described above **(see section 5.5)**.

**6.10 Processivity Analysis and Extrusion Speed Measurements**

We used polymer simulations in which the position and speed of the loop extruding cohesin and the 3D positions of all the monomers were known explicitly to test and validate an approach for computing (1) which intervals of the 3D distance vs. time traces were unique to intervals in which cohesin was actively pulling the monomers together and (2) how fast cohesin is moving in the chromatin (in nm/s and kb/s).  Note, even in cells or simulations with cohesin, there is not always a cohesin bound between any two observation points, and when it is bound, it is often involved in extruding the slack between the two sites and thus exerting minimal force on either, thus for (1) we cannot simply compare traces from cells with and without cohesin.  This process is described below in **Section 7**. With this empirically derived set of properties, determined from the simulations, we identified intervals likely containing extruding cohesin leading to processive motion of our labeled sites in several of our cell lines, as described in the main text. We found nearly all of these processive traces occurred in cells in which cohesin was intact, and not cells in which cohesin had been depleted (17 out of 19). It is possible that individual cohesin complexes escape degradation (as seen previously in other cohesin depletion experiments (*17*, *29*, *30*)) during the imaging time in individual cells to provide apparently processive motion in these 2 cases, or that other molecular motors  can occasionally induce processive motion.

To convert the measured extrusion speeds from nm/s to kb/s, we considered the size of both the nucleosome and the chromatin fiber as determined in cryoelectron tomography (*85*).

**7. Polymer Simulations for Processive Motion and Extrusion Speed**

***7.1 Simulation setup***

Simulations of the motion of confined chromosome polymers with and without loop extruding cohesin were performed using the polychrom package from open2c, as described in our previous publications. Code for replicating these simulations is included, with all the parameters defined and described therein.  Briefly, a polymer chain of 100,000 monomers (approximating 150 Mb of chromatin) was spatially confined using periodic boundary conditions to a density of 10% excluded volume. This corresponds to 1.5 kb per monomer, estimated at ~50 nm diameter, based on the size of nucleosome clutches in electron-tomography and STORM imaging data and the 150 bp per nucleosome (*85*, *149*). The simulations were run for 10,000 cohesin steps, with 200 steps of the molecular dynamics engine per cohesin step, a characteristic lifetime of cohesin of 100 steps, and an average separation of 200 monomers. Cohesin molecules loaded randomly as a pair of motor heads that bind adjacent monomers. These heads extrude loops through processive bidirectional motion of each head in opposite directions, 1 monomer per saved frame (simulating 1 second of real time) for an effective extrusion rate of 2 monomers per s (or 3 kb/s at our simulation resolution of 1 monomer per 1.5 kb). These parameter values were selected to be similar to those described in recent publications  (*4*, *20*, *30*, *95*, *150*), based on the current best estimates in the field for the density and spacing of cohesin based on comparisons of simulations with Hi-C data (*4*, *30*).

***7.2 Detection of processive motion***

We analyzed motion between pairs of monomers, not separated by and not proximal to CTCF sites, across a range of linker lengths.  To identify processive traces, we developed a list of criteria associated with processive motion, and explored different thresholds on these criteria to determine a set of parameters that provided reliable identification of (1) intervals when cohesin is bound to the polymer in the interval between the observations sites and (2) never occurred in simulations lacking cohesin. Note that since cohesin in these simulations binds stochastically, extrudes and falls off stochastically, even in the “with cohesin” simulations, two monomers may have no cohesin bound between them.  The processive criteria used were as follows: (1) total travel distance – the distance traveled needs to be sufficiently large relative to the typical distance between observation points to ensure that the points start from a stochastically stretched configuration. (2) The total number of observations which exhibited processive motion – a particle experiencing Brownian motion will still often step in the same direction in two (or five) consecutive intervals, but the probability of such a run decreases exponentially with the number of steps. Whereas a particle being pulled under an external force in a specific direction should move in the direction of that force for the duration of the force (as long as the force exceeds the forces exerted by the Brownian motion / interactions with the bath. (3) Minimal fraction of steps which reversed direction - As the extrusion simulation polymer is still experiencing Brownian motion as well, we found requiring perfectly processive motion to be too strict and eliminate traces that were otherwise unique to the extrusions simulations.  We therefore allowed a moderate level of back-stepping events, and varied this threshold as well.  (4) Maximal amount of reverse-travel distance during the interval – a few steps that technically reverse direction but with a close-to-zero amount are much more likely to be the effect of Brownian motion stalling/countering the external force of the cohesin motor than large steps in the reverse direction. Therefore we also limited the total distance of reverse steps allowed.  By empirically exploring these thresholds we found that a distance traveled equal twice the average separation between the polymer pair, with 16 total observations and a maximum of 20% reversed direction and a maxim of total distance reversed equal 20% provided 15 traces from the cohesin simulations that all had cohesin during the transition, and which enformed a relatively accurate calculation of extrusion speed from the slope of the trace (see **Fig. 4C-G**). This approach establishes an easy to implement empirical procedure for identifying intervals in the trajectories where cohesin is actively pulling. It is not an ‘optimal’ approach in any sense - the parameter choices are sufficient to identify intervals of cohesin pulling without generating false positives – it is undoubtable that many intervals of cohesin directed motion are still missed. We expect future investigations with more sophisticated tools from non-equilibrium statistical mechanics will provide better discrimination with greater sensitivity.

***7.3 Quantification of extrusion speed from single trajectories***

Extrusion speed in units of 3D distance (monomer diameters) was computed directly from the average slope of the processive search events (see **Fig. 4C** for example), as the total distance traveled / time. This was converted into kb (monomer steps by cohesin) by estimating the stretched length (observed at the start of the search event), guided by the known size of the polymer in the case of simulations and by the size of short stretched nucleosomal arrays in the case of the experimental data. Direct inspection of the full 3D position of all the intervening monomers in these simulations offered further support for this conversion factor. Using this scaling, we converted the 3D distance per unit time to kb per unit time - and recovered quantities close to the actual extrusion speed encoded at the start of the simulations. This suggests this protocol provides a workable estimate of extrusion speed in both 3D and 1D distance per unit time, using only the 3D positions of the individual loci pairs along the chromatin polymer, for traces that exhibit the processive characteristics described above.

***7.4 Quantification of extrusion bounds***

Simulation data can also be used to validate an approach for estimating confidence bounds on the extrusion speed. While we cannot be certain of the amount of chromatin that is stretched the ~1000 nm gap at the start of the extrusion process, we can be certain it is no longer than genomic separation between the labeled points and no shorter than an unbroken linear chain. In the case of simulations, if the length of chromatin between the “labeled” points is 150 kb, which puts an upper limit on the amount of extrusion which could have happened even if there had been no slack for cohesin to remove before the onset of processive motion. For the trace that covered 1000 nm in 20 frames this translates to an upper bound on the speed estimate 7.5 kb/fr. Meanwhile, it would require breaking the polymer backbone to span the 1000 nm with less than 20 kb of simulated polymer, meaning not less than 20 kb/20 fr for this trace or no-less than 1kb/fr - a generous lower bound on the extrusion speed. Note even these extreme bounds confine the true value within an order of magnitude.

As with the simulation data, we can use additional information to put strong bounds on the speed estimate.  For example, the processive walks of 10s in the cells with probes separated by 130 kb put a firm upper bound of 13 kb/s. To span the ~1200 nm requires at least 4kb of DNA (0.3 nm/bp), which puts a firm lower bound of 4kb/10s = 0.4kb/s. Note that this lower bound is a theoretical lower extreme as it requires DNA to be stretched into a naked rigid rod, allowing no nucleosome wrapping, or DNA bending. In most physiological contexts, the likely lower bound would be significantly larger due to nucleosome wrapping. For (fully stretched) linker lengths of 25~75 bp and ~150 bp of nucleosome wrapped DNA, corresponding range of lower bounds would be estimated to be 1.2 kb~2.8 kb/s (as nucleosome wrapping would reduce the effective length of the DNA by 3~7-fold), an order of magnitude larger than *in vivo* estimates.

**8. Other Analyses**

**8.1 Fluorescent in situ Microscopy (FISH) with Optical Reconstruction of Chromatin Architecture**

Fixed cell data to compare 3D distances from FISH with TRACK-IT was taken from our previously published investigations of chr6 from 60 kb to whole chromosome (150 Mb). We matched TRACK-IT cell lines with the closest possible ORCA probes for the comparison.  The positions of the TRACK-IT probes are as follows (in mm10 coordinates, chr6):

tetO = 51320704;

cuO= [51321893

51336653

51371176

51387332

51451617

51576421

51724680

50525643

49294139

38529185

124883209];

The positions corresponding centers of the ORCA probes selected were (in mm10 coordinates, chr6):

```
  51315001    51340001
    51315001    51365001
    51315001    51390001
    51315001    51440001
    51315001    51565001
    51315001    51715001
    51300000    50550000
    51300000    49300000
    51300000    38550000
    51500000   125000000
```

**8.2 Analysis of Hi-C data**

For comparison of TRACK-IT data with Hi-C, we used previously published data from mouse embryonic stem cells from Bonev et al 2017 (*62*), which remains the deepest sequenced mESC data available.  We compared the contact frequencies measured between our labeled TRACK-IT cell lines with the contact frequencies in the corresponding bins from the Hi-C data. Hi-C data was downloaded using Juicebox and normalized using the “balanced” option in Juicebox (*151*).  For loci pairs < 6 Mb a part, we binned the Hi-C data at 5kb using Juicebox (*151*). For loci pairs 6 - 30 Mb apart, we used a 25 kb bin and for loci >30 Mb apart we used a 100 kb bin size.  The use of larger bin sizes at greater separations was chosen to suppress the sampling noise in Hi-C, as even with the deepest sequencing ever published, Hi-C recovers relatively few reads per 5 kb bin at separations of >6 Mb, resulting in a “salt-and-pepper” pattern of reads counts that are not reproduced over replicates. This sampling error is typically addressed by using larger bins, as we do here.  In order to compare contact frequencies across bin sizes, all counts per bin were normalized to the 5 kb bin by dividing by the number of bins averaged – e.g. 25 kb bins represent a 5x5 tile of 5 kb bins, so the normalized read count was adjusted by dividing by 5^2^.  To facilitate visual comparison of normalized reads with the absolute contact frequency measured by live cell imaging (fraction of events across all traces in which the tracked loci were within 50 nm), we rescaled the normalized Hi-C reads by a factor 5e3 (**fig. S4B**). This is a purely visual alignment to facilitate comparison of the relative scaling – though it also represents an approximation of the conversion factor of Hi-C contacts, measured in units of “normalized Hi-C readcounts”, to physical contact frequency, measured as an actual fraction of the observed population, between 0 and 1.

**8.3 Analysis of ChIP-seq data**

Previously published ChIP-seq datasets for histone modifications (H3K27ac (GSE146451), H3K27me3 (GSE146451), H3K36me1 (GSE146451), H3K36me3 (GSE146451), and H3K9me3 (GSE180003)) and CTCF (GSE146451) as well as ATAC-seq (GSE99746) data were obtained from GEO as bigwig files (.bw format). Normalized signal intensities in 10kb windows with 10bp resolution centered around the genomic coordinates of each mapped transposons were generated using computeMatrix (*133*), resulting in a matrix (.mat) file. For tSNE analysis, signals spanning the 10kb window were first integrated, resulting in a single value for each dataset for each transposon reinsertion site. This 7-dimensional matrix was then put into tSNE embedding with Matlab. Epigenomic states were additionally annotated with previously published mESC ChromHMM data (*59*) by querying the corresponding genomic coordinates with a python script.

Supplement: Polymer Physics

As genetic elements connected in *cis* are by definition part of the same long polymer (a single molecule of DNA), some fundamental results from polymer physics can be useful in sanity checking the data, as well as defining appropriate minimal null models against which to compare and interpret polymer behavior. These comparisons have become standard practice in recent works using live-cell chromatin imaging. In this section we summarize some key results from both classic polymer physics and recent analytic theory and simulation, and discuss how they relate to the interpretation of our data.

**1. Mean squared change of displacement (MSCD) of simple polymer models**

The MSCD has been widely used in analysis of the dynamics of chromatin and its protein regulators. Here we briefly discuss the approach, its strengths, and its limitations, in light of both prior data and that presented in our work. The MSCD is defined as the average of the squared change in the 3D distance, <(*R*(s,*t_2_*) - *R*(*s,t_1_*))^2^ >, where *R* is the 3D distance between two monomers in a polymer, *s* is the number of monomers between them and *t* is time interval between the measurements of the 3D distance. It is thus an intuitive quantity of the magnitude of how much the two monomers change their separation over time.The MSCD is also called the pair-MSD, *M_2_*(*t*), in some works.  It is important to note that this quantity measures motion of the polymer relative to another point (monomer) on the same polymer.  It should not be confused with the mean square displacement (MSD, *M*(*t*)) of a single labeled point, relative to the lab frame (or the center of mass of the cell), which is also a popular metric. We focus on the reference frame of distance of one DNA locus relative to another, as this is the  biological frame of interest in the case of understanding cis-regulatory contacts, and the relevant reference for relating live-cell data to insights from Hi-C and chromosome-tracing microscopy approaches like ORCA.  It is also a convenient reference to bypass many other sources of motion which can be difficult to accurately track independently at the necessary nanoscale precision. These sources include the motion of coverglass relative to the objective, due for example to vibration of the stage or microscope, motion of the live cell relative to the coverglass, motion of the nucleus relative to the cell (in several cell types, including mESCs, the nuclei frequently rotate), motion of the whole chromosome relative to the nucleus, and changing deformation of the nucleus due to motion of the cytoskeleton. Many works have observed individual fluorescent chromatin loci in a single color, some of which have come to opposing biological conclusions based on the single-point MSDs (e.g. (*44*, *48*, *56*, *57*, *79*)). The data naturally measure displacements relative to the reference frame of the camera (the lab frame), it is challenging to measure the many additional sources of potential motion, and different approaches used require different assumptions about which form of motion is at play. In contrast, if two bright, point-source fluorescent labels are present on the same chromosome, their motion relative to one-another (rather than relative to the lab frame), is independent of these many additional sources of motion like stage-vibration or nuclear rotation. For these biological and technical reasons, we focus on MSCD.

The simplest model of a polymer structure is formed by connecting uniform steps of a (Brownian) random walk to form a freely jointed chain, or ***ideal chain*** (*76*, *155*). Just as the distance traveled by a random-walk scales with the square-root of time, *R*(*s*), the average 3D distance between two monomers separated by a chain-length of *s* along an ideal chain, scales with the square root of the number of monomers between them *R*(*s*) ~ *s*^½^. The simplest model of how this polymer moves is to assume harmonic bonds between the monomers, and to ignore any hydrodynamic effects of solvent. This model is typically called the ***Rouse polymer***, and it is simple enough for the MSCD to have an analytically tractable solution.

The limiting behaviors of the MSCD for the Rouse polymer have a simple form. For a single free Brownian particle, *M*(*t*) = D*t*, where D is the diffusion constant. For sufficiently large *s* and short *t*, the two monomers in an ideal chain move independently, and the MSCD, *M_2_*(*t*) ~ *t*^½^. At *t* sufficiently large for the particles to have explored completely the full range of distances allowed by the tether between them, the system is at steady state and *M_2_*(*t*) no longer increases with increasing *t*, but saturates at *M_2_*(*s*,*t*) ~ 2<*R(s)*^2^>, twice the variance of the 3D distances distribution at steady-state. The amount of time to approach this limit naturally increases with the intervening polymer length *s*.  It can be shown (*30*) that MSCD of the ideal chain follows:

$$M_{2}\left( t,s \right) \sim Dt^{\frac{1}{2}}(1-exp(\frac{-cs^{2}}{t})) +2<R(s)^{2}> erfc({(\frac{cs^{2}}{t})}^{\frac{1}{2}})$$

Where *a* and *b* are constants that respectively determine the diffusion speed at small *t* and the cross-over time between diffusion-dominated and tether dominated behavior, while <*R(s)*^2^> is the average square distance between the monomers at equilibrium.

A polymer that is confined in a spherical volume of radius much smaller than the relaxed radius of gyration of the corresponding ideal chain, or a polymer that is poorly soluble and minimizes surface interactions with the surrounding solvent by collapsing on itself, adopts a globular organization, with spatial scaling and movement dynamics that are distinct from the ideal chain. At equilibrium, the globule is highly knotted (and does not exhibit fractal scaling). However, if the polymer was initially unknotted, its collapsed configuration adopts a fractal organization (like the ideal chain), which is known as the “***crumpled globule***” (*156*, *157*) or “fractal globule” (*9*, *10*, *156*). In this case *R*(*s*) ~ *s*^⅓^, (*10*). Given the densely packed nature of chromosomes in mammalian nuclei, and the requirement that in mitosis at least they be unknotted so the chromosomes can segregate, it has been proposed that the crumpled/fractal globule is a reasonable model of genome packaging (or at least a better null model than the ideal chain) (*10*, *158*). In support of this, microscopy evidence has shown repeated support for the *s*^⅓^ scaling (*57*, *74*, *159*), and Hi-C has shown the contact frequency scales with *s*^-1^, which is expected consequence of *R*(*s*)~*s*^⅓^ (*9*). Motivated in part by the potential relevance of this polymer model for genome organization, theorists have predicted a short-time scale scaling for the crumpled globule of *M_2_*(*t*) ~ *t*^2/5^, which is supported as well by simulations (*77*, *78*).  The steady state still converges to  2<*R(s)*^2^> in this crumpled model as well.

**2. Effect of additional tethers on MSCD, and simulations of cohesin tethers**

The basic theory for the MSCD provides a prediction for how addition of crosslinks spanning from a few to a few hundred kilobases might affect the MSCD of polymer segments a few kilobases to a hundred thousand kilobases in length. Prior work has estimated the abundance of extrusive cohesin in G1-stage mouse embryonic stem cells to be on the scale of 1 molecule per hundred kilobases (186-372 kb) (*87*). For the shortest polymer segments, we expect most of the time there will be no cohesin bound, and thus the impact on the average displacement will be minimal compared to a cohesin free case. For polymer segments sufficiently long to typically have a cohesin we expect cohesin to have a significant impact on the MSCD. One simple prediction is that on average these segments will behave like a simple polymer where the tether length has been shortened by an amount equal to this average loop size (assuming the effects of the added energy from cohesin extrusion are negligible).

To test these predictions, we conducted simulations of crumpled polymers with and without loop extrusion and computed the MSCD for monomers at different separations.  All conditions exhibited a MSCD  scaling  ~ *t*^2/5^ at short time scales (**Fig A**).  At longer timescales, the MSCD deflected towards saturation, as expected by the theory (**Fig A**). Simulations with cohesin deflected sooner and saturated at lower MSCD than without cohesin. Notably, these MSD curves from more highly spaced monomers with cohesin/extruders (e.g. 499 monomers, as shown in Fig A), collapse to follow the profile exhibited by those of shorter length without cohesin (somewhere between the 240 monomer and 166 monomers in the simulations in **Fig A**). A similar behavior was seen in our experimental data, where the MSCDs start with scaling  ~ *t*^2/5^, and the 134 kb separated pair with cohesin behaves more like the 55 kb cohesin free case (or a little between the 20 kb and 55 kb case). This also suggests that the 134 kb domain is on average a little more than 60% extruded.

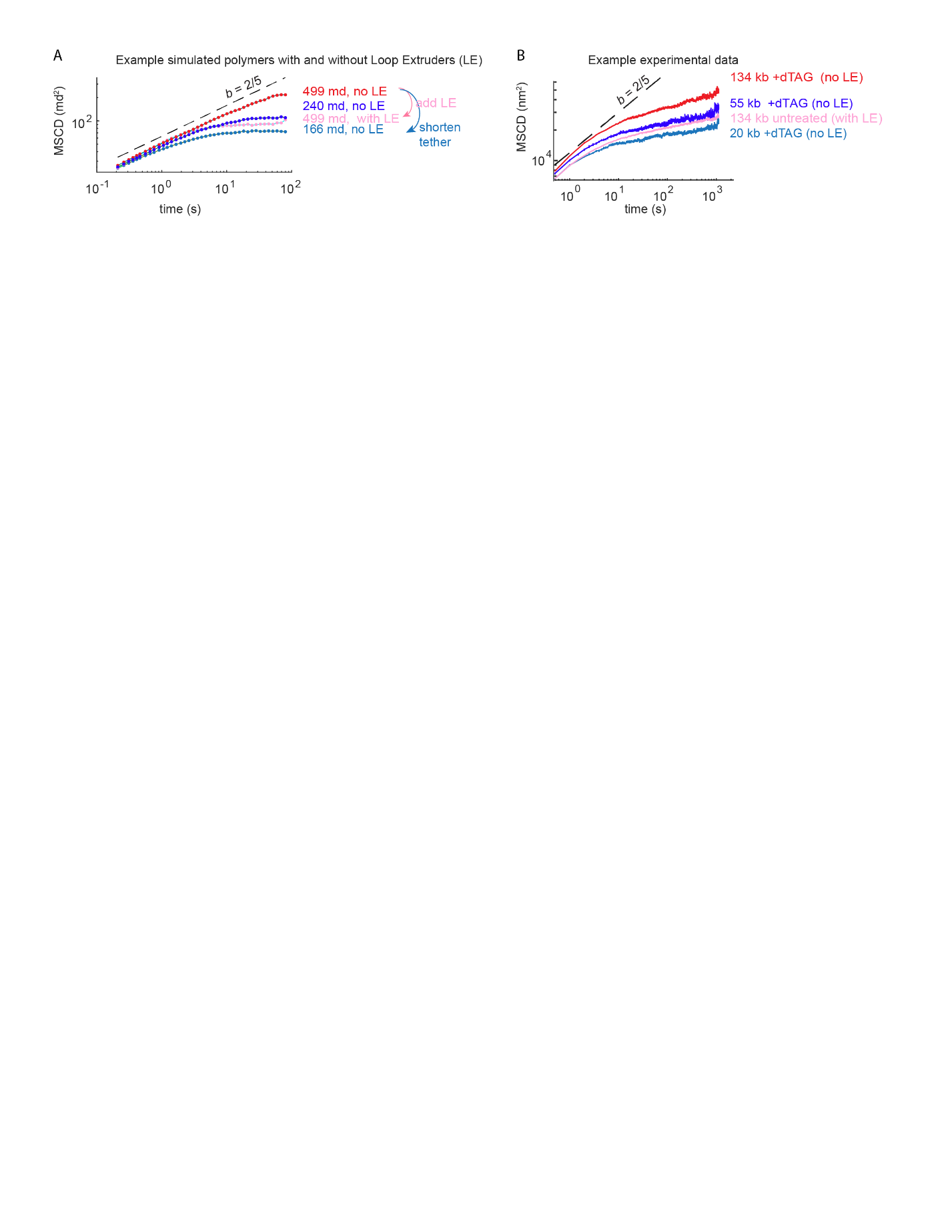

**Figure | Loop extrusion behaves like a tether from the standpoint of mean square change in displacement. A** MSCD (in squared monomer diameters (md)), for four different simulated polymers, initialized in an unknotted and yet spatially confined configuration.   **B**. Experimental MSCD data from Figure 3, overlaying four different experimental conditions indicated. Dashed line shows the slope from a power-law function with an exponent b=2/5th, as predicted for the initial scaling of the MSCD of the crumpled globule.

Notably, recent work by Gabriele and colleagues (*30*), and work by Mach and colleagues (*29*), also measured MSCDs in mouse embryonic stem cells with and without cohesin, and both reported a similar increase in the MSCD upon cohesin depletion, which was interpreted as an effect of tether length. It should be noted that both studies covered around two-orders of magnitude of time and focused primarily on the long-term dynamics, and neither capture the short-term dynamics of the polymer due to the 20 to 30s sampling intervals chosen.  This limited sampling limits the ability to assess power-law scaling behavior (*160*), and likely explains why the MSCD curves in these earlier studies are primarily shifted on the y-axis, rather than starting equivalent and deflecting away from the power-law scaling at the cross-over times expected for polymers with shorter tethers. The longer sampling interval may also contribute to differences in the observed initial scaling exponents reported in each of these studies. Mach et al also presented simulations to show this shift.

**3. Search time behavior in simple polymers**

Search processes are typically characterized by the search time, the average time it takes for the system to go from some initial condition to some ‘search complete’ state (*161*).  This analysis has been central to the previous studies like how transcription factors find their target sites (*162*–*164*).  In the case of contacts in a polymer, a natural choice for the starting state is the average distance of the pair of monomers of interest, while contact can be defined with a proximity threshold. Unlike the search time for enzymes and their substrates, which often diffuse free in a confined 3D (or 2D/membrane) environment, elements of a polymer are tethered together, and shorter tethers will decrease the search time, so the search time, *T*, is naturally a function of the polymer length between the monomers of interest, *s*.

The dependence of the average search time on the linear separation of the polymer can also be approximated by extracting the characteristic time-scale of diffusion from the MSCD(*t*,*s*), and the characteristic length-scales from the spatial scaling relation *R*(*s*) (see definitions in section 1 above) (*57*, *78*). Recall for a simple unconstrained random walk, *R*(s)~*s*^½^, whereas for the self-avoiding random-walk *R*(*s*)~*s*^⅗^, and for a crumpled globule, *R*(*s*)^⅓^ (*10*, *76*).

Using equation (1) above for the simple polymer MSCD, in the limit where *cs*^2^*^/t^* is small, i.e. not steady-state since we are interested in search times, we can ignore the steady state part of the MSCD (which will be negligibly small) and write the MSCD as:

M_2_(*s,t*)  ~ *t*^½^(1-exp(*x*))    with  *x* = -c*s^2/t^*

Taylor expanding exp(*x*) => 1+*x*, we have

M_2_(*s,t*)  ~ *t*^½^(1- (1+*x*)) = *t*^½^(-*x*)  = t^½^*cs*^2/^*^t^* = $\frac{cs^{2}}{t^{-1/2}}$

M_2_(*s,t*) ~ *R*(*s*)^2^

Substituting in the length scaling relationship for the ideal polymer from *R*(*s*)~ s^½^, we have:

$\frac{cs^{2}}{t^{-1/2}}$ ~ s^1/2×2^

And simplifying and ignoring the constant terms since we are interested in the scaling relation

*s* ~ *t*^½^

*t* ~ *s*^2^

Which gives the prediction that the search time, *T*, scales as the square of the separation, *T_ideal_*(*s*)~*s*^2^. This is also known as the Rouse time.

Or, substituting in for the length scaling of a crumpled structure instead of an ideal polymer, we have:

M_2_(*s,t*)  ~ *t*^⅖^  and *R*(*s*)^2^ ~ *s*^⅓×2^

*t*~*s*^⅔^*s*^5/2^ = *s*^5/3^.

Here we see an improvement in the search time scaling, though it is still super-linear, *T_crumpled_*(*s*)~*s*^5/3^.

Finally, a search process driven by a single processive motor that binds one site and pulls in the polymer at a fixed speed *v* kb/s, will obviously scale linearly in its search time with the separation, so *T_motor_*(*s*) ~*s*^1^.

Notably, in all of these cases the search time grows with inter-loci genomic separations faster than observed for intra-TAD elements in cells with cohesin, T(s) ~ s^0.60^,  95%-CI=(0.44, 0.76)).

**4. Is the genome a self-similar polymer with scale-free search times?**

While we assert that the search time for intra-TAD elements with cohesin is scaling with a lower exponent than the powerlaws above, resulting in ever faster searches as a function of intra-TAD separation, we caution that it should not be inferred that cohesin mediated intraTAD search is itself a powerlaw / scale-free behavior. The cohesin depleted data roughly follow the power-law scaling across nearly four-orders of magnitude in genomic separation.  In contrast, the search time scaling with cohesin in our data persists across a little less than two orders of magnitude, which is too short an interval to make a reliable evaluation of scale-free behavior, nor is the data a particularly tight fit to the line.

**Supplemental Figure Legends**

**Fig. S1. Comparison of recent live-cell imaging approaches and results.  (A)** Schematic of the different labeling approaches used. **(B)** Comparison of trace properties across published experiments and the current work. All cell lines are color-coded by the genomic separation between the labels.

**Fig. S2. Sleeping Beauty insertions of tile TRACK labels across the genome**

**(A)** Karyogram with TRACK-IT label reinsertions (red lines). Green arrowhead, parental insertion location. **(B)** Quantification of reinsertion numbers by chromosome. **(C)** Distribution of genomic separation from the parental site (within chr6) following mobilization with Sleeping Beauty. **(D)** ChromHMM states for local (*n* = 131, within 1 Mb from the initial insertion) and non-local (*n* = 623, >1 Mb from the initial insertion or trans chromosomal) reinsertions. **(E)** Sleeping Beauty reinsertions occur neutral to chromatin states. Local and non-local reinsertions both show no evidence of enrichment for the assessed epigenetic marks (H3K27ac (*152*), accessibility (ATAC-seq) (*153*), H3K36me1 (*152*), H3K36me3 (*152*), CTCF (*152*), H3K27me3 (*152*) and H3K9me3 (*154*)).

**Fig. S3. Reintegration site mapping via in situ transcript sequencing (isT-seq). (A)** in situ transcript sequencing workflow. **(B)** in situ transcript mapped to the genome before ITR filtering (Raw reads) or after ITR filtering. **(C)** Example bed file output from findJunctions. **(D)** Example mapping results visualized on Integrated Genome Browser. **(E)** Mapping results for parental TRACK-IT cell line, post-remobilization, randomly integrated TRACK-IT transposon from a plasmid donor, and no transposon control.

**Fig. S4. Cross-validation of TRACK-IT with Hi-C and ORCA.  (A)** Comparison of the contact frequency (50 nm threshold) from two biological replicate TRACK-IT experiments. **(B)** Comparison of contact frequencies from Hi-C and TRACK-IT as a function of genomic separation.  Both replicates combined and overlaid are shown. Correlation strength is quantified by Pearson’s *R* at top.  **(C)** Comparison of 3D distance measured by FISH vs. TRACK-IT.  **(D)** Effect of threshold choice (shown at top) on the correlation of Hi-C and TRACK-IT contact frequencies.

**Fig. S5. Chromatic registration. (A)** schematic of the chromatic registration process is shown in the top left, with six numbered steps. Remaining panels illustrate the steps of chromatic correction, and show snapshots of image output by our chromatic correction software to validate the process.  See Code Availability to access the code and sample data for the chromatic corrections, which will produce interactive versions of these plots for greater detail.  **(B)** Step 1: Three panels show the raw image before correction (left), with the two color channels shown in red and cyan, and corrected image of the whole field (middle) -  note the background appears cyan due to the padding of the cyan channel with zeros in the alignment. The middle panel is aligned to 7 pixel resolution (which dramatically increases the processing speed). The right panel shows the final alignment, to single pixel resolution, which is computed using only a fraction of the image for speed (since the coarse 7-pixel registration now guarantees the images will be aligned).  Units are in camera pixels (108 nm/pixel). The magnitude of the shift and rotation needed to complete the alignment is printed at top.  **(C)** Step 2: Projections of the red and cyan channel spots, shown in *xy, yz*, and *xz*. The cyan and red circles are centered on the 3D fitted locations. A modest offset due to chromatic aberration from the lenses can be seen.  **(D)** Step 3: Top  panels, scatter plots, in *xy* and *xz* respectively, of the original position (open cyan circles) and corrected position (filled cyan circles) of the cyan channel spots, relative to the target position of the spots appearance in the red channel, following the application of a second order polynomial correction field to maximize alignment. Bottom panels show vector fields of displacement (in *xy* and *yz*) for all corrected beads. The length of the arrows is enlarged for visualization, which shows the expected distance dependent effect. The correction minimum is off-center relative to the field of view due to the residual, sub-pixel error in aligning the cameras in the first step.  **(E)** Step 4: Top panels - original distribution of 3D displacements and *xy* displacements due to chromatic aberration, shown as a histogram. The median of the distribution is provided at top.  Bottom panels show the distribution of displacements between corrected localization between the channels, as in top panels.

**Fig. S6. Photostability for all 11 cell lines, compared to earlier labeling approaches.**  **(A)** Graph of the fraction of traces with detected data per frame for all experiment sets reported (shown as different colors) in each of the indicated publications (*4*–*8*).  **(B)** As in A, but for the 11 TRACK-IT cell lines introduced in Figs. 1-2 (color coded by genomic separation, as indicated in **Fig. 1**).  Note some transient autofluorescence background in the first frames impacts the ability to reliably detect and track some spots, resulting in a systematic increase in detection fraction.  Some variation in the detection fraction resulted from loci moving temporarily out of the narrow z-scan range.  Lower panel shows results for the 4 cell lines first introduced in **Fig. 3G-K**.

**Fig. S7. Sister chromatid analysis. (A)** Unreplicated loci from cells in the G1 state of the cell-cycle appear as singlet spots. In G2, the replicated sisters can be resolved as doublets, which stochastically overlap and separate.  By detecting the appearance of doublets at any point through the movie, traces can be assigned as G2.  **(B)** Schematic illustration of the effect of temporal resolution on doublet tracing.  Individual spots move between frames, and sister loci can also appear at different positions in the *z*-stack (and thus also separate in acquisition time.  At relatively slow acquisition rates (top panel) it can be ambiguous which spot belongs to which sister. This problem is substantially reduced by high temporal resolution sampling.  **(C)** Example TRACK-IT data of a challenging pair of sisters tracked through time (left panels). A gap in the detection of sister 2 (black dashed box) during a window in which both sisters were moving in a coherent direction, (*x* and *y* are both steadily increasing by chance in this window, probably as a result of cell movement) - leads the nearest-step tracing algorithm to inaccurately swap sisters. Correcting for motion in the reconstruction allows a more likely assignment of the correct sister identities (right panels). Data from the other color channel (cyan) shows the more cis-paired locus on sister exhibiting correlated motion and closer proximity to its molecular partner.

**Fig. S8. Genomic separation-cumulated MSCD plots**. **(A)** Overlay of MSCD on log-scale axes (top) and linear scale axes (bottom). Black dotted line shows power-law scaling with an exponent of 1/2, and 2/5, as expected by different polymer diffusion models. Colored lines on the far right indicate the 2<R^2^> population variance, at which the MSCD is expected to saturate.   **(B)** As in (A), but for cells with cohesin depletion.

**Fig. S9. Search time corrections and validation. (A) Raw observed vs. KM corrected search times** Search time data as in **Fig. 2D**, left panel showing the difference between raw, observed +/- cohesin, middle panel the effect of KM correction on the data from untreated cells, and right panel the effect of KM correction on the data from +dTAG (cohesin degraded) cells.

**(B) KM correction compensates for undersampling the search times at large separations in Rouse Polymer simulations.**  Simulations of Rouse Polymers for 100,000 s - search processes and passage times follow power-law scaling with the theoretically predicted scaling relation time ~ separation squared (far right). Light blue dots indicate the raw / observed median passage times, dark blue dots are the KM corrected values.  The pink line is a linear fit to the linear portion of the raw data, the dark blue line is a linear fit to the KM corrected values. The slopes of these lines (on the log-log graph, i.e. scaling exponents in the power-law) are indicated.  Observe that truncating the simulations after shorter observation windows (10,000, 1,000, or 100 frames) leads to undersampling of the long search times, which preferentially occur for long separations. This undersampling leads to a flattening of the scaling, which systematically depresses the exponent).  When the data is sufficiently sampled, this effect is negligible (both the KM corrected data and the raw data are close to the theoretically expected exponent 2 for much of the trace).  However, as truncation becomes more significant at shorter observation windows, the raw passage time scaling is more affected. At the shortest observation window, the KM correction does not fully correct the data to achieve the theoretical value, but it substantially reduces the effect of undersampling.  **(C) Estimating long search times by changing the threshold and rescaling.**  Black dots show the ‘true’ search process scaling for a Rouse polymer using a contact threshold of theta=2 and 100,000 frames of simulated data.  Red markers show the scaling of the search time to reach theta = 4 (circle), 8 (plus), or 16 (square), using only 3,300 frames after rescaling by a simple multiplicative factor. Red dots below these symbols show the original, non-rescaled values.  The multiplicative factor for each is computed simply by matching the last value in the previous series and the first value of the next series.  So T1(x1:x2), the time to reach theta = D1 for separation x1:x2 is estimated by time to reach theta=D2 from x1:x2, T2(x1:x2), then rescaling T2 by *a* so that T1(x1) = a*T2(x1).

**Fig. S10. Cell cycle effects.** Comparison of G1 vs G2 traces, with a contact threshold of 50 nm, for **(A)** 11 TRACK-IT cell lines (**Fig. 1D** and **3D-F**) Data from untreated cells (intact cohesin) are in red, dTag treated (cohesin degraded) cells in blue. Error bars show 95% confidence intervals. Solid lines, data fit for untreated cells (red, intra-TAD only) and cohesin depleted cells (blue). Black dashed line, a = 5/3, predicted from crumpled polymer. Grey dashed line, a = 2, predicted from ideal chain. **(B)** as in (A) but for 4 lines in chr 15 (**Fig. 3G-K**).

**Fig. S11.  Contact threshold effects.**  **(A)** search time (top row), search frequency (middle row), and average contact duration (bottom row) shown for the indicated contact thresholds, from 30 to 500 nm, from 11 TRACK-IT cell lines (**Fig. 1D** and **3D-F**). Data from untreated cells (intact cohesin) are in red, dTag treated (cohesin depleted) cells in blue. Errorbars show 95% confidence intervals for search time and search frequency, standard-error of the mean (SEM) for contact duration.  For search time distribution for 50 or 100 nm thresholds, **see fig. S13**. **(B)** as in (A) but for 4 cell lines in chr15 (**Fig. 3G-K**) shown for the indicated contact thresholds from 30 to 200nm.

**Fig. S12. correlation between contact duration and chromatin state.** **(A)** Correlation between average chromatin profile of the indicated mark (e.g. H3K27ac ChIP-seq signal or ATAC-seq signal, as described in **fig. S2**). in a 5 kb region around the insertion site and the contact duration (with a 50 nm contact threshold) for all 15 loci-pairs analyzed (top row). **(B)** As in (A), but for the correlation between the peak value of the indicated chromatin mark and the contact duration. Individual insertion sites are color-coded as indicated.  **(C)** Graphs of the normalized chromatin profiles (ChIP-seq or ATAC-seq) at each of the insertion sites, colored as in (A).

**Fig. S13. Search time distributions. (A)** Cumulative distribution functions (CDF) for the observed passage times for probes of different separation. A 50 nm contact threshold was used to define contact in the search. The separation between the probes are listed above each plot. Data in red is from cells with cohesin (untreated) and data in blue is from cells in which cohesin was depleted with dTag treatment. **(B)** As in A, except for using a 100 nm contact threshold.  Data with fewer than 5 search events was not plotted, as sparse data provides little characterization of the distribution.

Table S1. DNA sequences (oligos, gBlocks, and plasmids) used in the study

| **DNA Oligos** |  |  |
| --- | --- | --- |
| **Name** | **Sequence** | **Notes** |
| HoxA_2_1_left_F | acgaggccctttcgtcttcaagaaTGTGGTAGTCATCACAGTGACG | Used to clone left homology arm (chr6:51319538-51320703 (mm10)) into TRACK-IT cargo plasmid with EcoRI |
| HoxA_2_1_left_R | ggactaacccattcctagggcgaaTACCTTAGCATATCTGACAAAGAAT | " |
| HoxA_2_1_right_F | atgtatgctatacgaagttatcatAAGTGGGATAGAAGCCGACTCC | Used to clone right homology arm (chr6:51320704-51321899 (mm10)) into TRACK-IT cargo plasmid with KpnI |
| HoxA_2_1_right_R | ccactcctttcaagacctagccatTGAGCTATACAAACCGAGACC | " |
| NeoR_5half_F | agtgatagagagggttgtgatactctgaggcggaaagaaccagc | Used to clone SV40 promoter 5' to sleeping beauty transposon into TetO array vectors with SpeI, 460bp product |
| NeoR_5half_R | taaacttccgacttcaactgtatatctcttgatcgatctttgcaaaagc | " |
| NeoR_3half_F | taaacttccgacttcaactgtataaggatcgtttcgcatgattgaac | Used to clone NeoR gene (Aminoglycoside 3'-phosphotransferase) 3' to sleeping beauty transposon into TetO array vector with SpeI, 1160bp product |
| NeoR_3half_R | gcatacattatacgaagttatactgaacaaacgacccaacaccg | " |
| SB | tatacagttgaagtcggaagtttac | To clone SB_CuO into 48x TetO homology donor vector with SpeI along with NeoR fragments. The sequence is same for R2 ~3900bp product. |
| HoxA_2_1_sg_F | CACCGTCAGATATGCTAAGGTAAAG | sgRNA oligo cloned into px458 vector linearized with BsaI to target TRACK-IT cargo to 129 allele-specific chr6:51,320,704 (mm10) locus. |
| HoxA_2_1_sg_R | AAACCTTTACCTTAGCATATCTGAC | " |
| ePB_UbC_rTetR_tdSG_1_F | gtagtcccttctcggcgattctg | To linearize ePB_UbC_rTetR_tdSG plasmid via PCR to assemble with candidate linker sequences to improve tdSG expression in mESCs. ePB_UbC_rTetR_tdSG plasmid is amplified in three fragments with 24bp homology between each fragments to promote isothermal assembly. |
| ePB_UbC_rTetR_tdSG_1_R | GAGATGGGCTTCTAAAGTTTCAGATTGAC | " |
| ePB_UbC_rTetR_tdSG_2_F | ATGGCCTCTACCCCATTCAAGTTC | " |
| ePB_UbC_rTetR_tdSG_2_R | gtcagaagtaagttggccgcagtg | " |
| ePB_UbC_rTetR_tdSG_3_F | cactgcggccaacttacttctgac | " |
| ePB_UbC_rTetR_tdSG_3_R | cagaatcgccgagaagggactac | " |
| Huang_26_F | TCTGAAACTTTAGAAGCCCATCTCGGTGGAGGAGGCAGCGGGGGAGGTGGCTCCGGGGGGGGCGGCTCTGAAGCAGCGGCCAAGGAGGCTGCTGCCAAA ATGGCCTCTACCCCATTCAAGTTC | Huang_26 candidate linker sequence between the two StayGold sequences in ePB_UbC_rTetR_tdSG plasmid. |
| Huang_26_R | GAACTTGAATGGGGTAGAGGCCATTTTGGCAGCAGCCTCCTTGGCCGCTGCTTCAGAGCCGCCCCCCCCGGAGCCACCTCCCCCGCTGCCTCCTCCACCGAGATGGGCTTCTAAAGTTTCAGA | " |
| Chen_18_F | TCTGAAACTTTAGAAGCCCATCTCAAGGAAAGTGGCTCTGTGTCCTCAGAGCAACTCGCCCAGTTCAGAAGCCTGGACATGGCCTCTACCCCATTCAAGTTC | Chen_18 candidate linker sequence between the two StayGold sequences in ePB_UbC_rTetR_tdSG plasmid. |
| Chen_18_R | GAACTTGAATGGGGTAGAGGCCATGTCCAGGCTTCTGAACTGGGCGAGTTGCTCTGAGGACACAGAGCCACTTTCCTTGAGATGGGCTTCTAAAGTTTCAGA | " |
| Chen_14_F | TCTGAAACTTTAGAAGCCCATCTCGAAGGGAAGAGCAGTGGCTCTGGATCAGAGAGCAAATCCACCATGGCCTCTACCCCATTCAAGTTC | Chen14 candidate linker sequence between the two StayGold sequences in ePB_UbC_rTetR_tdSG plasmid. |
| Chen_14_R | GAACTTGAATGGGGTAGAGGCCATGGTGGATTTGCTCTCTGATCCAGAGCCACTGCTCTTCCCTTCGAGATGGGCTTCTAAAGTTTCAGA | " |
| 4xSAGG_F | TCTGAAACTTTAGAAGCCCATCTCTCTGCCGGTGGCAGCGCCGGGGGCTCCGCAGGAGGCAGTGCTGGAGGGATGGCCTCTACCCCATTCAAGTTC | 4xSAGG candidate linker sequence between the two StayGold sequences in ePB_UbC_rTetR_tdSG plasmid. |
| 4xSAGG_R | GAACTTGAATGGGGTAGAGGCCATCCCTCCAGCACTGCCTCCTGCGGAGCCCCCGGCGCTGCCACCGGCAGAGAGATGGGCTTCTAAAGTTTCAGA | " |
| 6xSAGG_F | TCTGAAACTTTAGAAGCCCATCTCTCAGCTGGGGGCTCCGCAGGAGGGAGCGCGGGTGGCAGTGCCGGCGGCAGCGCCGGGGGTTCTGCTGGAGGAATGGCCTCTACCCCATTCAAGTTC | 6xSAGG candidate linker sequence between the two StayGold sequences in ePB_UbC_rTetR_tdSG plasmid. |
| 6xSAGG_R | GAACTTGAATGGGGTAGAGGCCATTCCTCCAGCAGAACCCCCGGCGCTGCCGCCGGCACTGCCACCCGCGCTCCCTCCTGCGGAGCCCCCAGCTGAGAGATGGGCTTCTAAAGTTTCAGA | " |
| 8xSAGG_F | TCTGAAACTTTAGAAGCCCATCTCAGTGCCGGAGGAAGCGCTGGCGGCAGCGCCGGTGGCTCCGCAGGTGGGTCCGCGGGAGGCTCAGCCGGGGGTTCTGCTGGGGGCTCTGCAGGAGGGATGGCCTCTACCCCATTCAAGTTC | 8xSAGG candidate linker sequence between the two StayGold sequences in ePB_UbC_rTetR_tdSG plasmid. |
| 8xSAGG_R | GAACTTGAATGGGGTAGAGGCCATCCCTCCTGCAGAGCCCCCAGCAGAACCCCCGGCTGAGCCTCCCGCGGACCCACCTGCGGAGCCACCGGCGCTGCCGCCAGCGCTTCCTCCGGCACTGAGATGGGCTTCTAAAGTTTCAGA | " |
| 1147_7_gRNA_F | CACCGGATCTCTTAATAAGACCTGG | sgRNA oligo cloned into px458 vector linearized with BsaI to target 48-mer TetO array into 129 allele-specific chr15:11,477,063 (mm10) locus ("empty TAD") |
| 1147_7_gRNA_R | AAACCCAGGTCTTATTAAGAGATCC | " |
| 1107_gRNA_F | CACCGGTTGCAGTCTCAGCCCCCAA | sgRNA oligo cloned into px458 vector linearized with BsaI to target 48-mer CuO array into 129 allele-specific chr15:11,071,424 (mm10) locus (~400kb across empty TAD border) |
| 1107_gRNA_R | AAACTTGGGGGCTGAGACTGCAACC | " |
| 1137_gRNA_F | caccgAGGAGGTATTTGGAATCACA | sgRNA oligo cloned into px458 vector linearized with BsaI to target 48-mer CuO array into 129 allele-specific chr15:11,373,113 (mm10) locus (~100kb across empty TAD border) |
| 1137_gRNA_R | aaacTGTGATTCCAAATACCTCCTc | " |
| 1157_gRNA_F | caccgTATAATCTTCAAAGCCATCA | sgRNA oligo cloned into px458 vector linearized with BsaI to target 48-mer CuO array into 129 allele-specific chr15:11,568,485 (mm10) locus (~100kb within empty TAD border) |
| 1157_gRNA_R | aaacTGATGGCTTTGAAGATTATAc | " |
| 1187_gRNA_F | CACCGCATTGGATGGAATTCAAACA | sgRNA oligo cloned into px458 vector linearized with BsaI to target 48-mer CuO array into 129 allele-specific chr15:11,880,005 (mm10) locus (~400kb within empty TAD border) |
| 1187_gRNA_R | AAACTGTTTGAATTCCATCCAATGC | " |
| 1147_donor_F | A*T*TTGTAATATTACAATATATACATATATAAATTAGACACTGATGAGAAAAAAAGTGCCACCTGACGTCTAAG | Used to PCR 48-mer TetO array to insert into chr15:11,477,063 (mm10) locus ("empty TAD") with 50bp homology arms on each side. Stars (*) denote phosphorothioate bond modification to improve editing efficiency |
| 1147_donor_R | A*A*ATGTCCTCTTCCAATCACTTTCTCTGTTTCCATTTTATCTTCCCTTGTCTTAAGCTAGCAGCGCTCTCG | " |
| 1187_donor_F | A*A*AGTGAAAAGAAAAAAAAAAAACATATCAAAGCATTGGATGGAATTCAAAAAAGTGCCACCTGACGTCTAAG | Used to PCR 48-mer CuO array to insert into chr15:11,880,005 (mm10) locus (~400kb within empty TAD border) with 50bp homology arms on each side. Stars (*) denote phosphorothioate bond modification to improve editing efficiency |
| 1187_donor_R | T*C*GTGTGTGTGGTAAGCTCTGTGCTTTATCGCCTGTGTAGATTTCCATGTCTTAAGCTAGCAGCGCTCTCG | " |
| 1107_donor_F | C*C*TCTAAGCTTTAAGTCAGAGGTGGTTTTGCAAGTTGCAGTCTCAGCCCCAAAAGTGCCACCTGACGTCTAAG | Used to PCR 48-mer CuO array to insert into chr15:11,071,424 (mm10) locus (~400kb across empty TAD border) with 50bp homology arms on each side. Stars (*) denote phosphorothioate bond modification to improve editing efficiency |
| 1107_donor_R | A*A*AGTCAGAGACCCAGGCTGAGCAGACGCAAGACAAACCCACACCCATTGCTTAAGCTAGCAGCGCTCTCG | " |
| 1137_donor_F | G*G*AGGGCCTTGCCTTTGGAAGTTGTTGCCATTTTCATAACTTCTCCATGTAAAAGTGCCACCTGACGTCTAAG | Used to PCR 48-mer CuO array to insert into chr15:11,373,113 (mm10) locus (~100kb across empty TAD border) with 50bp homology arms on each side. Stars (*) denote phosphorothioate bond modification to improve editing efficiency |
| 1137_donor_R | T*T*TTTTTCCAAAGAAGAGGAAAGGATAATCTATAGGAGGTATTTGGAATCCTTAAGCTAGCAGCGCTCTCG | " |
| 1157_donor_F | G*A*CTCCAGTGACCTCTTAGACCTTTCTGATGTAAATGTCGATGACCTTGAAAAAGTGCCACCTGACGTCTAAG | Used to PCR 48-mer CuO array to insert into chr15:11,568,485 (mm10) locus (~100kb within empty TAD border) with 50bp homology arms on each side. Stars (*) denote phosphorothioate bond modification to improve editing efficiency |
| 1157_donor_R | G*A*AATTCTTTAAGAAAAAAAACTTCTGACCAAGTATAATCTTCAAAGCCACTTAAGCTAGCAGCGCTCTCG | " |
| Rad21_c-term_sgRNA_F | CACCG CCACGGTTCCATATTATCTG | sgRNA oligo cloned into px458 vector linearized with BsaI to target FLAG::dTag::P2A::PuroR cassette into C-term of Rad21 (chr15:51,964,057(mm10)) |
| Rad21_c-term_sgRNA_R | AAAC CAGATAATATGGAACCGTGG C | " |
| Rad21_dTag_F | A*G*CCGTACAGTGACATCATTGCAACCCCTGGACCACGGTTCCATATTATCggcggtggaggatccatggg | Used to PCR FLAG::dTag::P2A::PuroR insert with 50bp homology arms for C-term of Rad21 on each side. |
| Rad21_dTag_R2 | t*t*tgtatgtactagtgagttatcactagctcgaacacatctagctcctcacgctccaggcttccgtg | " |
| **gBlocks** |  |  |
| SB_5'ITR | acgaggccctttcgtcttcaagaatatacagttgaagtcggaagtttacatacacttaagttggagtcattaaaactcgtttttcaactactccacaaatttcttgttaacaaacaatagttttggcaagtcagttaggacatctactttgtgcatgacacaagtcatttttccaacaattgtttacagacagattatttcacttataattcactgtatcacaattccagtgggtcagaagtttacatacactaagttttcgccctaggaatgggttagtcc | Sleeping beauty 5' ITR sequence with homology arms for assemblying TRACK-IT cargo plasmid. |
| T7_SB_3'ITR | tcgagagcgctgctagcttaaggtTAATACGACTCACTATAGGGATAATggagtcgagtgtatgtaaacttctgacccactgggaatgtgatgaaagaaataaaagctgaaatgaatcattctctctactattattctgatatttcacattcttaaaataaagtggtgatcctaactgacctaagacagggaatttttactaggattaaatgtcaggaattgtgaaaaagtgagtttaaatgtatttggctaaggtgtatgtaaacttccgacttcaactgtataaccatcgatagatctgggcccgtg | T7 promoter and Sleeping beauty 3' ITR sequence with homology arms for assembling TRACK-IT cargo plasmid. |
| **Plasmid** |  |  |
| **Name** | **Notes** | **Source** |
| TRACK-IT_HoxA_2-1 | TRACK-IT cargo with homology arms for targeting into chr6:51,320,704 (mm10) locus. | This study |
| ePB_UbC_rTetR_tdStayGold-4xSAGG | PiggyBac transposon vector used to stably express rTetR::StayGold-TI | This study |
| 48-mer_CuO-BlaR | 48-mer CuO array plasmid with blasticidin resistance cassette | This study |
| 48-mer_CuO-NeoR | 48-mer CuO array plasmid with neomycin resistance cassette | This study |
| 4_flag_dtag_p2a_puroR_pUC57 | FLAG::dTag::P2A::PuroR cassette | This study |
| pSP2-48-merTetO-EFS-BLaR | 48-mer TetO array plasmid with blasticidin resistance cassette | a gift from Huimin Zhao (Addgene plasmid # 118712 ; http://n2t.net/addgene:118712 ; RRID:Addgene_118712) |
| epB_UbC_CymRV5-nls-Halox2_DEx4 | PiggyBac transposon vector used to stably express CymR::2xHaloTag a gift from Orion Weiner (Addgene plasmid # 119907 ; http://n2t.net/addgene:119907 ; RRID:Addgene_119907 | a gift from Orion Weiner (Addgene plasmid # 119907 ; http://n2t.net/addgene:119907 ; RRID:Addgene_119907) |
| pCMV(CAT)T7-SB100 | Sleeping Beauty transposase (SB100) expression vector | a gift from Dr. Zsuzsanna Izsvak (Addgene plasmid # 34879 ; http://n2t.net/addgene:34879 ; RRID:Addgene_34879) |

**Table S2. Cell lines generated in the study**

| **Name** | **Description** |
| --- | --- |
| TetO/CuO NeoR C2 | TRACK-IT parent strain in HoxA locus chr:6 51320704, + strand ("pre-jump"). First clonal line |
| TetO/CuO NeoR C12 | TRACK-IT parent strain in HoxA locus chr:6 51320704, + strand ("pre-jump"). Second clonal line |
| 1.2kb rad21 dTag A2 | "5kb" line in HoxA locus (the genomic separation in the paper is shown as center-to-center between TetO and CuO array). CuO reinsertion coordinate is chr6:51321893, - strand (mm10). |
| 16kb rad21 dTag A9 | "20kb" line in HoxA locus . CuO reinsertion coordinate is chr6:51336653, - strand (mm10). This line has an additional CuO reinsertion in chr11:6430191, - strand. |
| 50kb rad21 dTag D5 | "55kb" line in HoxA locus . CuO reinsertion coordinate is chr6:51371176, + strand (mm10). |
| 76kb rad21 dTag A7 | "70kb" line in HoxA locus . CuO reinsertion coordinate is chr6:51387332, - strand (mm10). |
| 130kb rad21 dTag B1 | "134kb" line in HoxA locus . CuO reinsertion coordinate is chr6:51451617, + strand (mm10). |
| 255kb rad21 dTag C2 | "260kb" line in HoxA locus . CuO reinsertion coordinate is chr6:51576421, + strand (mm10). |
| 400kb rad21 dTag A2 | "407kb" line in HoxA locus . CuO was inserted in chr6:51724680, + strand (mm10). |
| 800kb rad21 dTag A5 | "799kb" line in HoxA locus . CuO was inserted in chr6:50525643, + strand (mm10). |
| 2Mb rad21 dTag A4 | "2Mb" line in HoxA locus . CuO reinsertion coordinate is chr6:49294139, + strand (mm10). |
| 12Mb rad21 dTag B4 | "12Mb" line in HoxA locus . CuO reinsertion coordinate is chr6:38529185, - strand (mm10). |
| 73Mb rad21 dTag A12 | "73Mb" line in HoxA locus . CuO reinsertion coordinate is chr6:124883209, + strand (mm10). |
| 1107_A11 | 409kb across the "Empty TAD" border. TetO was inserted in chr15:11477063, + strand, and CuO was inserted in chr15:11071425, + strand. |
| 1137_E7 | 108kb across the "Empty TAD" border. TetO was inserted in chr15:11477063, + strand, and CuO was inserted in chr15:11373113, + strand. |
| 1157_G9 | 95kb within the "Empty TAD" border. TetO was inserted in chr15:11477063, + strand, and CuO was inserted in chr15:11568485, + strand. |
| 1187_E9 | 407kb within the "Empty TAD" border. TetO was inserted in chr15:11477063, + strand, and CuO was inserted in chr15:11880005, + strand. |

**Table S3. Datasets used in the study**

| **Cell line** | **Dataset** | **Series** | **Series with GSM** | **Source** |
| --- | --- | --- | --- | --- |
| 1.2kb rad21 dTag A2 | RNA-seq (isT-seq mapping) | GSE289566 | GSM8794322 | this study |
| 16kb rad21 dTag A9 | RNA-seq (isT-seq mapping) | GSE289566 | GSM8794323 | this study |
| 50kb rad21 dTag D5 | RNA-seq (isT-seq mapping) | GSE289566 | GSM8794324 | this study |
| 76kb rad21 dTag A7 | RNA-seq (isT-seq mapping) | GSE289566 | GSM8794325 | this study |
| 130kb rad21 dTag B1 | RNA-seq (isT-seq mapping) | GSE289566 | GSM8794326 | this study |
| 255kb rad21 dTag C2 | RNA-seq (isT-seq mapping) | GSE289566 | GSM8794327 | this study |
| 2Mb rad21 dTag A4 | RNA-seq (isT-seq mapping) | GSE289566 | GSM8794328 | this study |
| 12Mb rad21 dTag B4 | RNA-seq (isT-seq mapping) | GSE289566 | GSM8794329 | this study |
| 73Mb rad21 dTag A12 | RNA-seq (isT-seq mapping) | GSE289566 | GSM8794330 | this study |
| TetO/CuO NeoR C12 | RNA-seq (isT-seq mapping) | GSE289566 | GSM8794319 | this study |
| TetO/CuO NeoR C12 | RNA-seq (isT-seq mapping) | GSE289566 | GSM8794320 | this study |
| F123 | RNA-seq (isT-seq mapping) | GSE289566 | GSM8794318 | this study |
| F123 | RNA-seq (isT-seq mapping) | GSE289566 | GSM8794321 | this study |
| F123 | CTCF ChIP-seq | GSE146451 | GSM4386047 | Kubo et al. 2021 |
| F123 | H3K27ac ChIP-seq | GSE146451 | GSM4386085 | Kubo et al. 2021 |
| F123 | H3K27me3 ChIP-seq | GSE146451 | GSM4386143 | Kubo et al. 2021 |
| F123 | H3K4me1 ChIP-seq | GSE146451 | GSM4386107 | Kubo et al. 2021 |
| F123 | H3K4me3 ChIP-seq | GSE146451 | GSM4386125 | Kubo et al. 2021 |
| J1 | H3K9me3 ChIP-seq | GSE180003 | GSM5444713 | Zhang et al. 2019 |
| E14 | ATAC-seq | GSE99746 | GSM2651154 | Tastemel et al. 2017 |
| JM8.N4 | Micro-C | GSE130275 | GSE130275 | Hsieh et al. 2019 |
| E14 | Hi-C | GSE96107 | GSE96107 | Bonev et al. 2017 |
