## Supplemental Figures for "Kinetic organization of the genome revealed by ultra-resolution, multiscale live imaging"

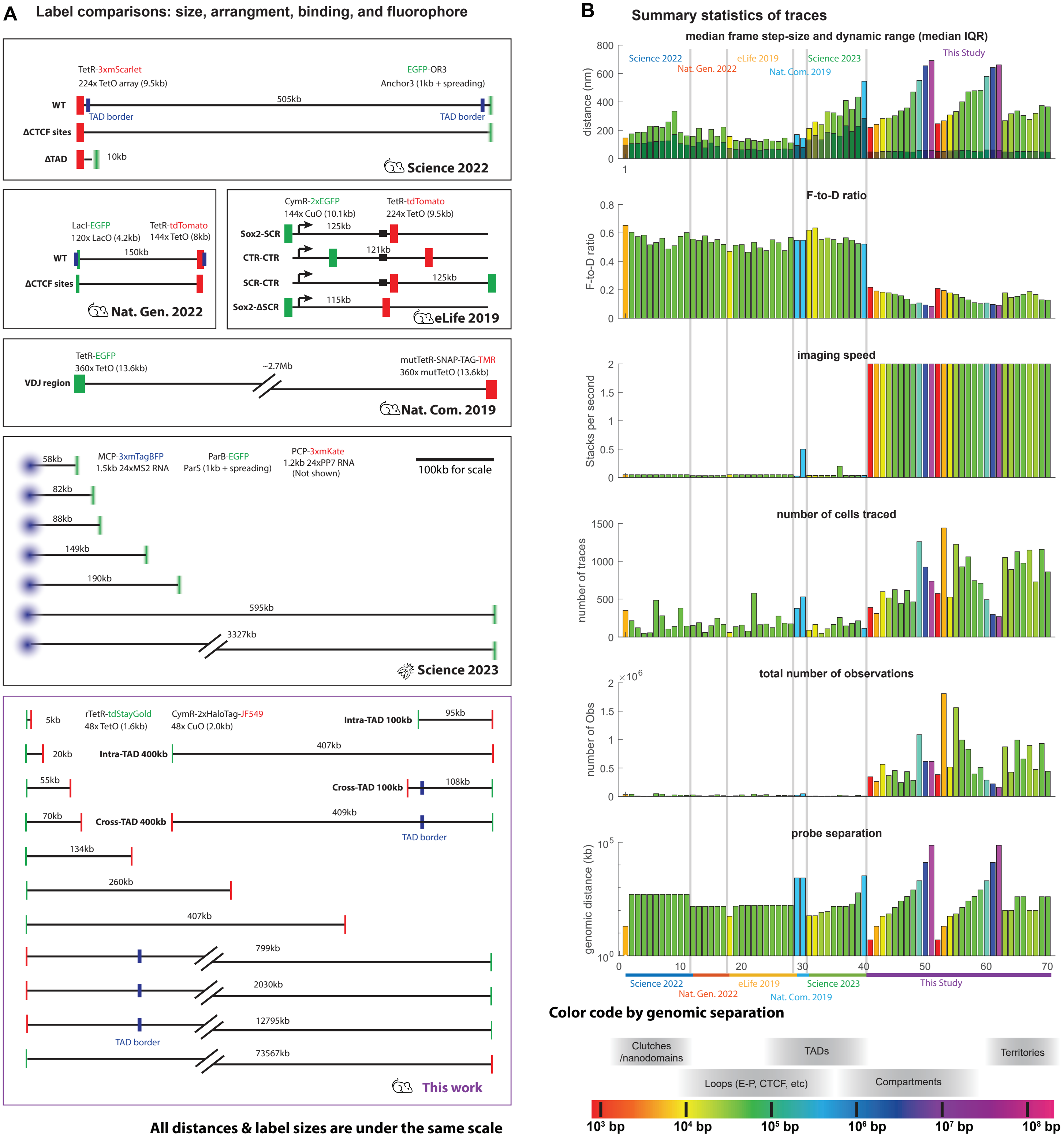
A
Label comparisons: size, arrangment, binding, and fluorophore
TetR-3xmScarlet
224x TetO array (9.5kb)
EGFP-OR3
Anchor3 (1kb + spreading)
WT
505kb
TAD border
TAD border
ΔCTCF sites
ΔTAD
10kb
Science 2022
LacI-EGFP
120x LacO (4.2kb)
TetR-tdTomato
144x TetO (8kb)
150kb
ΔCTCF sites
Nat. Gen. 2022
CymR-2xEGFP
144x CuO (10.1kb)
TetR-tdTomato
224x TetO (9.5kb)
Sox2-SCR
125kb
CTR-CTR
121kb
SCR-CTR
125kb
Sox2-ΔSCR
115kb
eLife 2019
TetR-EGFP
360x TetO (13.6kb)
~2.7Mb
mutTetR-SNAP-TAG-TMR
360x mutTetO (13.6kb)
VDJ region
Nat. Com. 2019
58kb
MCP-3xmTagBFP
1.5kb 24xMS2 RNA
ParB-EGFP
ParS (1kb + spreading)
PCP-3xmKate
1.2kb 24xPP7 RNA
(Not shown)
100kb for scale
82kb
88kb
149kb
190kb
595kb
3327kb
Science 2023
5kb
rTetR-tdStayGold
48x TetO (1.6kb)
CymR-2xHaloTag-JF549
48x CuO (2.0kb)
Intra-TAD 100kb
95kb
20kb
Intra-TAD 400kb
407kb
55kb
Cross-TAD 100kb
108kb
70kb
Cross-TAD 400kb
409kb
TAD border
134kb
260kb
407kb
799kb
2030kb
12795kb
TAD border
73567kb
This work
All distances & label sizes are under the same scale
B
Summary statistics of traces
median frame step-size and dynamic range (median IQR)
distance (nm)
Science 2022
Nat. Gen. 2022
eLife 2019
Nat. Com. 2019
Science 2023
This Study
F-to-D ratio
imaging speed
Stacks per second
number of cells traced
number of traces
total number of observations
number of Obs
probe separation
genomic distance (kb)
Science 2022
Nat. Gen. 2022
eLife 2019
Nat. Com. 2019
Science 2023
This Study
Color code by genomic separation
Clutches
/nanodomains
TADs
Territories
Loops (E-P, CTCF, etc)
Compartments
10³ bp
10⁴ bp
10⁵ bp
10⁶ bp
10⁷ bp
10⁸ bp

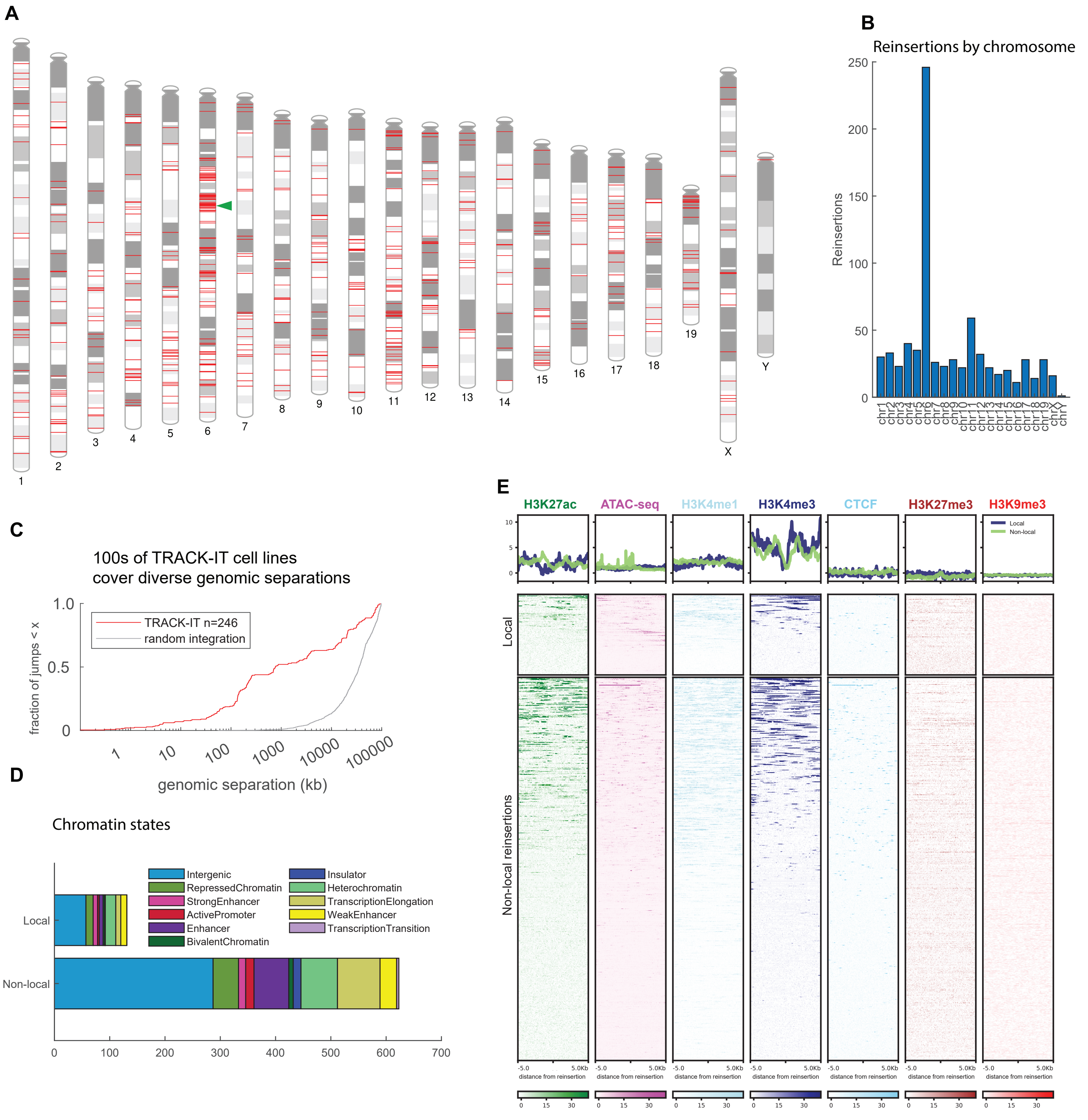
A
X
Y
B
Reinsertions by chromosome
Reinsertions
250
200
150
100
50
0
chr1 chr2 chr3 chr4 chr5 chr6 chr7 chr8 chr9 chr10 chr11 chr12 chr13 chr14 chr15 chr16 chr17 chr18 chr19 chrX chrY
C
100s of TRACK-IT cell lines
cover diverse genomic separations
fraction of jumps < x
1.0
0.5
0
TRACK-IT n=246
random integration
1
10
100
1000
10000
100000
genomic separation (kb)
D
Chromatin states
Local
Non-local
Intergenic
RepressedChromatin
StrongEnhancer
ActivePromoter
Enhancer
BivalentChromatin
Insulator
Heterochromatin
TranscriptionElongation
WeakEnhancer
TranscriptionTransition
0
100
200
300
400
500
600
700
E
H3K27ac
ATAC-seq
H3K4me1
H3K4me3
CTCF
H3K27me3
H3K9me3
Local
Non-local
10
5
0
Local
Non-local reinsertions
-5.0
5.0Kb
distance from reinsertion
0
15
30

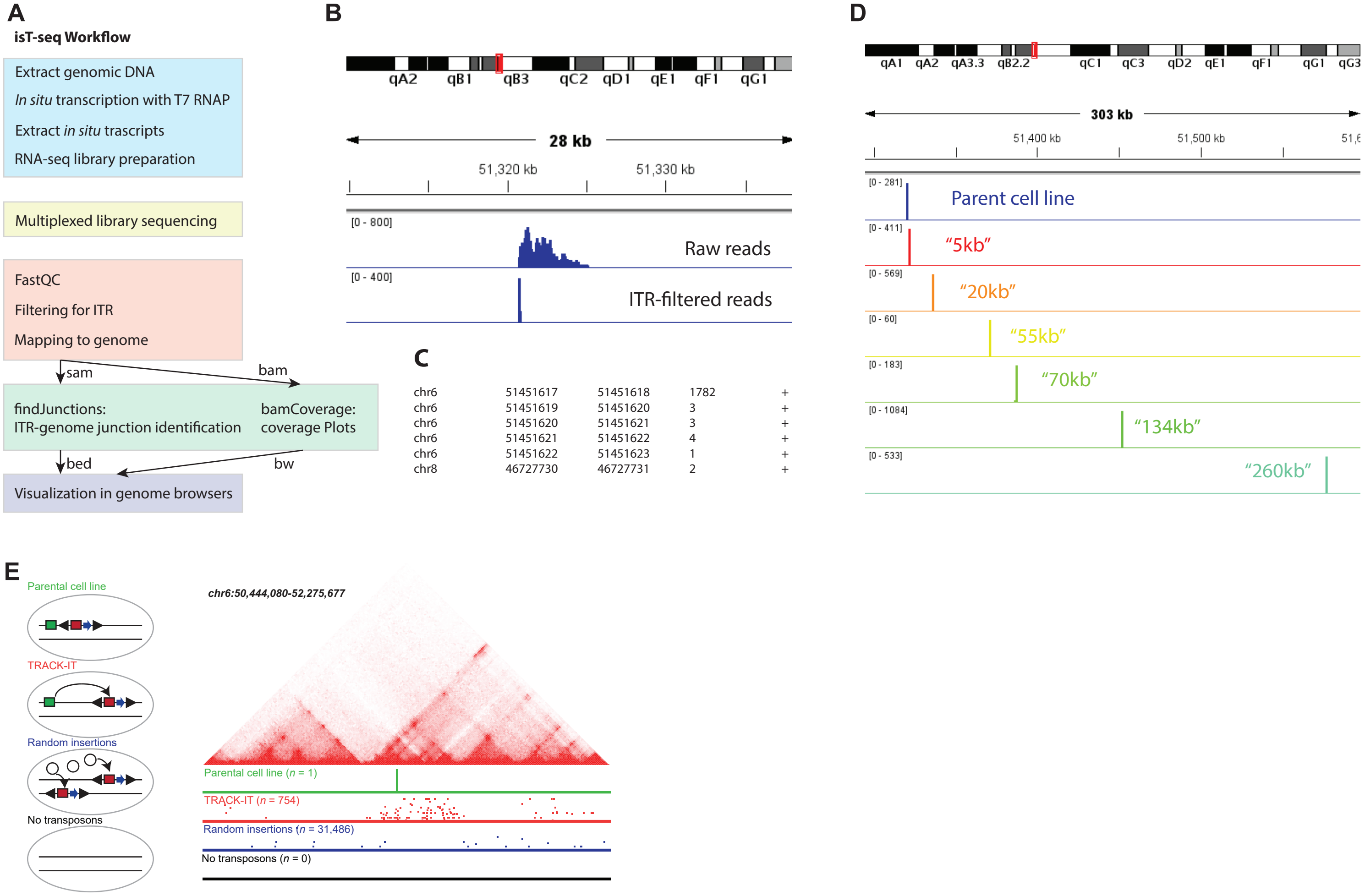
A
isT-seq Workflow
Extract genomic DNA
In situ transcription with T7 RNAP
Extract in situ trascripts
RNA-seq library preparation
Multiplexed library sequencing
FastQC
Filtering for ITR
Mapping to genome
sam
bam
findJunctions:
ITR-genome junction identification
bamCoverage:
coverage Plots
bed
bw
Visualization in genome browsers
B
qA2 qB1 qB3 qC2 qD1 qE1 qF1 qG1
28 kb
51,320 kb
51,330 kb
[0 - 800]
Raw reads
[0 - 400]
ITR-filtered reads
C
chr6 51451617 51451618 1782 +
chr6 51451619 51451620 3 +
chr6 51451620 51451621 3 +
chr6 51451621 51451622 4 +
chr6 51451622 51451623 1 +
chr8 46727730 46727731 2 +
D
qA1 qA2 qA3.3 qB2.2 qC1 qC3 qD2 qE1 qF1 qG1 qG3
303 kb
51,400 kb
51,500 kb
[0 - 281]
Parent cell line
[0 - 411]
"5kb"
[0 - 569]
"20kb"
[0 - 60]
"55kb"
[0 - 183]
"70kb"
[0 - 1084]
"134kb"
[0 - 533]
"260kb"
E
Parental cell line
TRACK-IT
Random insertions
No transposons
chr6:50,444,080-52,275,677
Parental cell line (n = 1)
TRACK-IT (n = 754)
Random insertions (n = 31,486)
No transposons (n = 0)

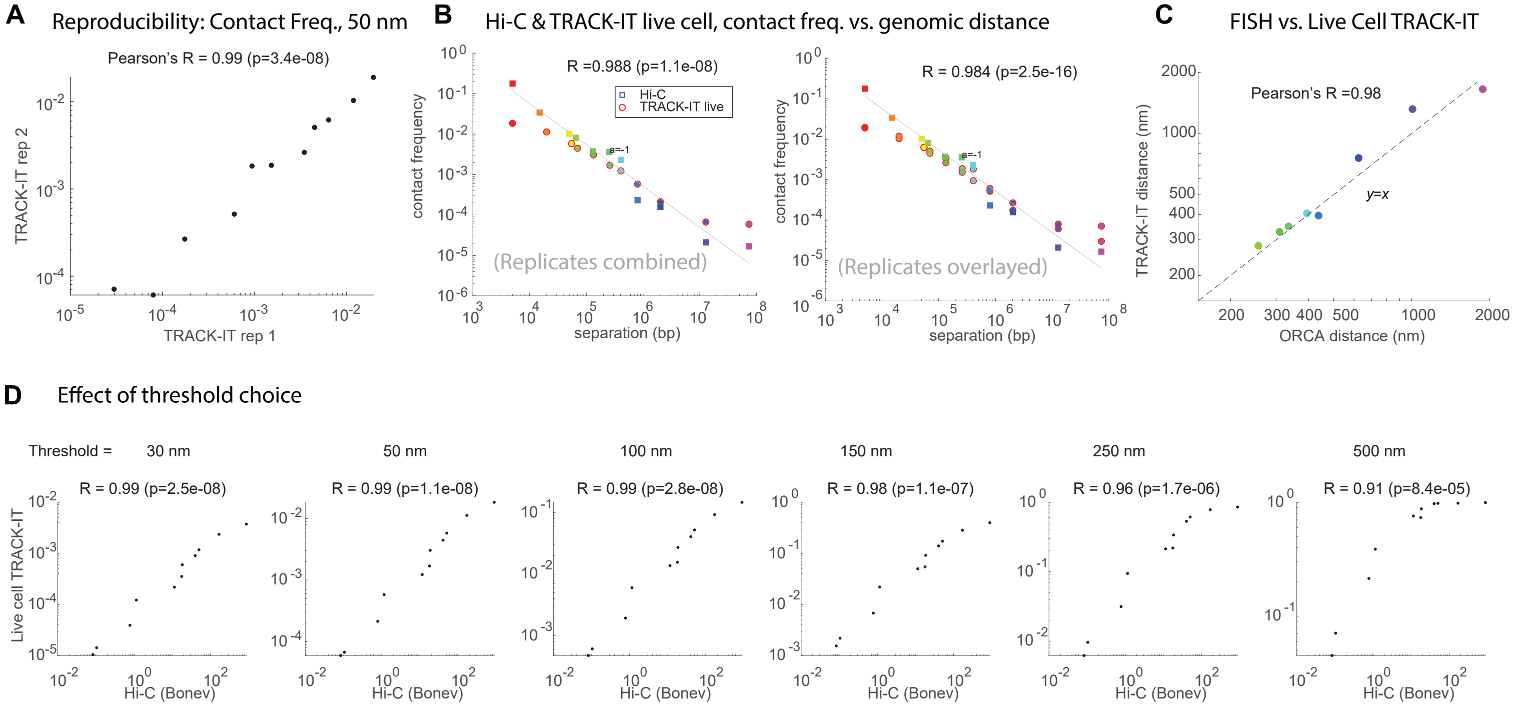
A
Reproducibility: Contact Freq., 50 nm
Pearson's R = 0.99 (p=3.4e-08)
TRACK-IT rep 2
TRACK-IT rep 1
B
Hi-C & TRACK-IT live cell, contact freq. vs. genomic distance
R =0.988 (p=1.1e-08)
Hi-C
TRACK-IT live
a=-1
contact frequency
separation (bp)
(Replicates combined)
R = 0.984 (p=2.5e-16)
(Replicates overlayed)
C
FISH vs. Live Cell TRACK-IT
Pearson's R =0.98
y=x
TRACK-IT distance (nm)
ORCA distance (nm)
D
Effect of threshold choice
Threshold = 30 nm
50 nm
100 nm
150 nm
250 nm
500 nm
R = 0.99 (p=2.5e-08)
R = 0.99 (p=1.1e-08)
R = 0.99 (p=2.8e-08)
R = 0.98 (p=1.1e-07)
R = 0.96 (p=1.7e-06)
R = 0.91 (p=8.4e-05)
Live cell TRACK-IT
Hi-C (Bonev)

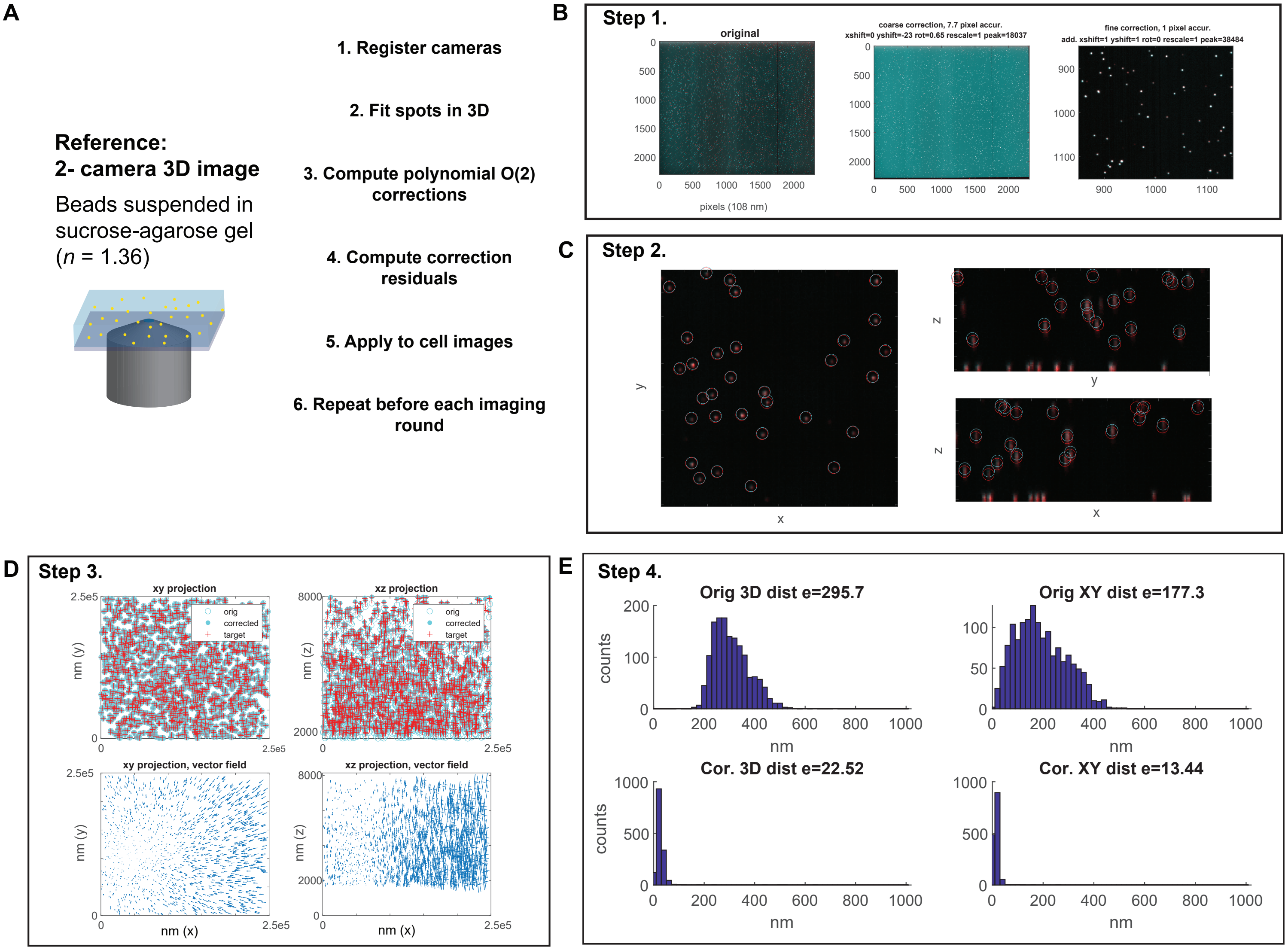
A
Reference:
2- camera 3D image
Beads suspended in sucrose-agarose gel (n = 1.36)
1. Register cameras
2. Fit spots in 3D
3. Compute polynomial O(2) corrections
4. Compute correction residuals
5. Apply to cell images
6. Repeat before each imaging round
B
Step 1.
original
pixels (108 nm)
coarse correction, 7.7 pixel accur.
xshift=0 yshift=-23 rot=0.65 rescale=1 peak=18037
fine correction, 1 pixel accur.
add. xshift=1 yshift=1 rot=0 rescale=1 peak=38484
C
Step 2.
D
Step 3.
xy projection
xz projection
orig
corrected
target
nm (y)
nm (z)
xy projection, vector field
xz projection, vector field
nm (x)
E
Step 4.
Orig 3D dist e=295.7
Orig XY dist e=177.3
Cor. 3D dist e=22.52
Cor. XY dist e=13.44
counts
nm

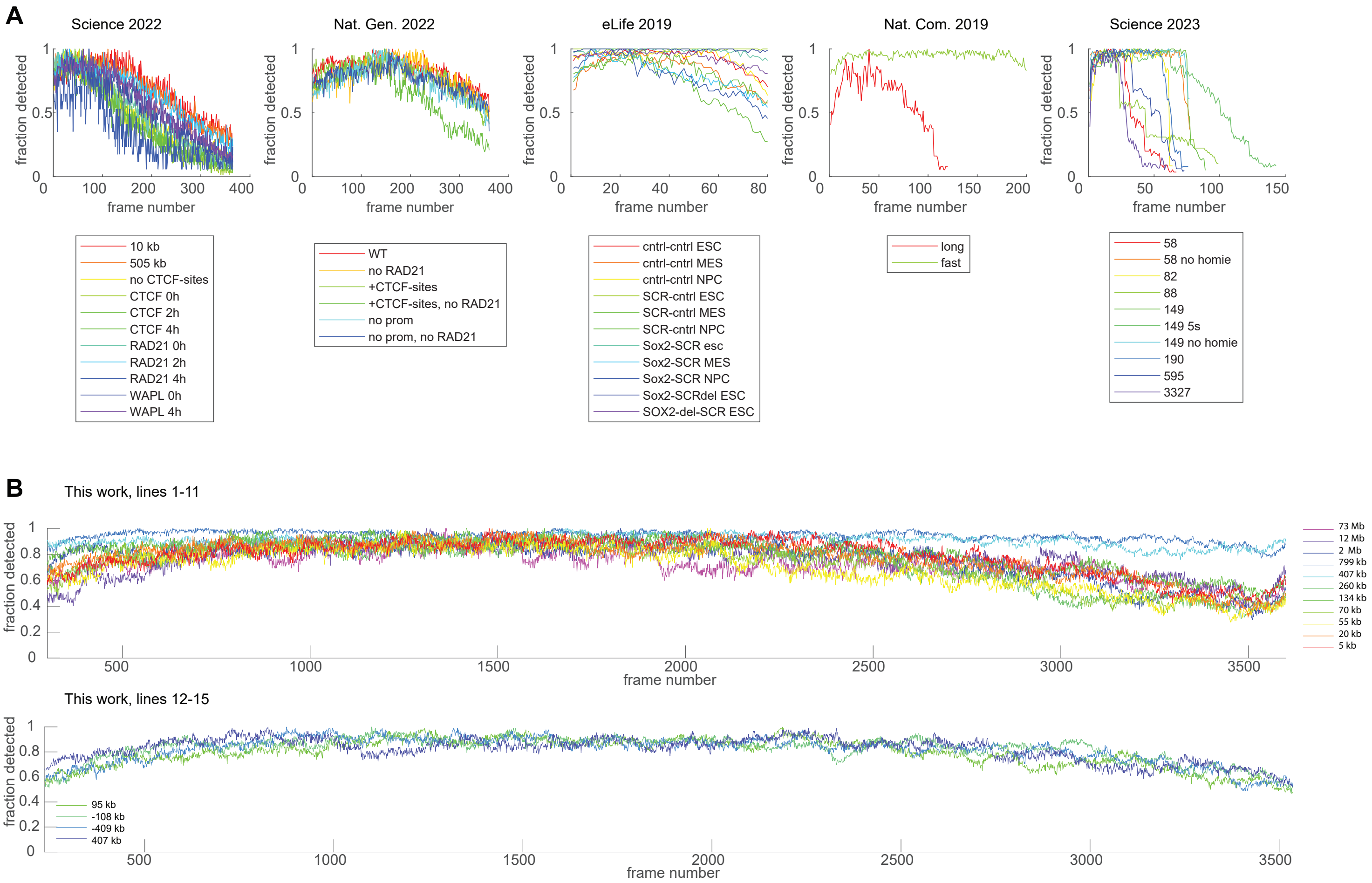
A
Science 2022
Nat. Gen. 2022
eLife 2019
Nat. Com. 2019
Science 2023
fraction detected
frame number
10 kb
505 kb
no CTCF-sites
CTCF 0h
CTCF 2h
CTCF 4h
RAD21 0h
RAD21 2h
RAD21 4h
WAPL 0h
WAPL 4h
WT
no RAD21
+CTCF-sites
+CTCF-sites, no RAD21
no prom
no prom, no RAD21
cntrl-cntrl ESC
cntrl-cntrl MES
cntrl-cntrl NPC
SCR-cntrl ESC
SCR-cntrl MES
SCR-cntrl NPC
Sox2-SCR esc
Sox2-SCR MES
Sox2-SCR NPC
Sox2-SCRdel ESC
SOX2-del-SCR ESC
long
fast
58
58 no homie
82
88
149
149 5s
149 no homie
190
595
3327
B
This work, lines 1-11
73 Mb
12 Mb
2 Mb
799 kb
407 kb
260 kb
134 kb
70 kb
55 kb
20 kb
5 kb
This work, lines 12-15
95 kb
-108 kb
-409 kb
407 kb

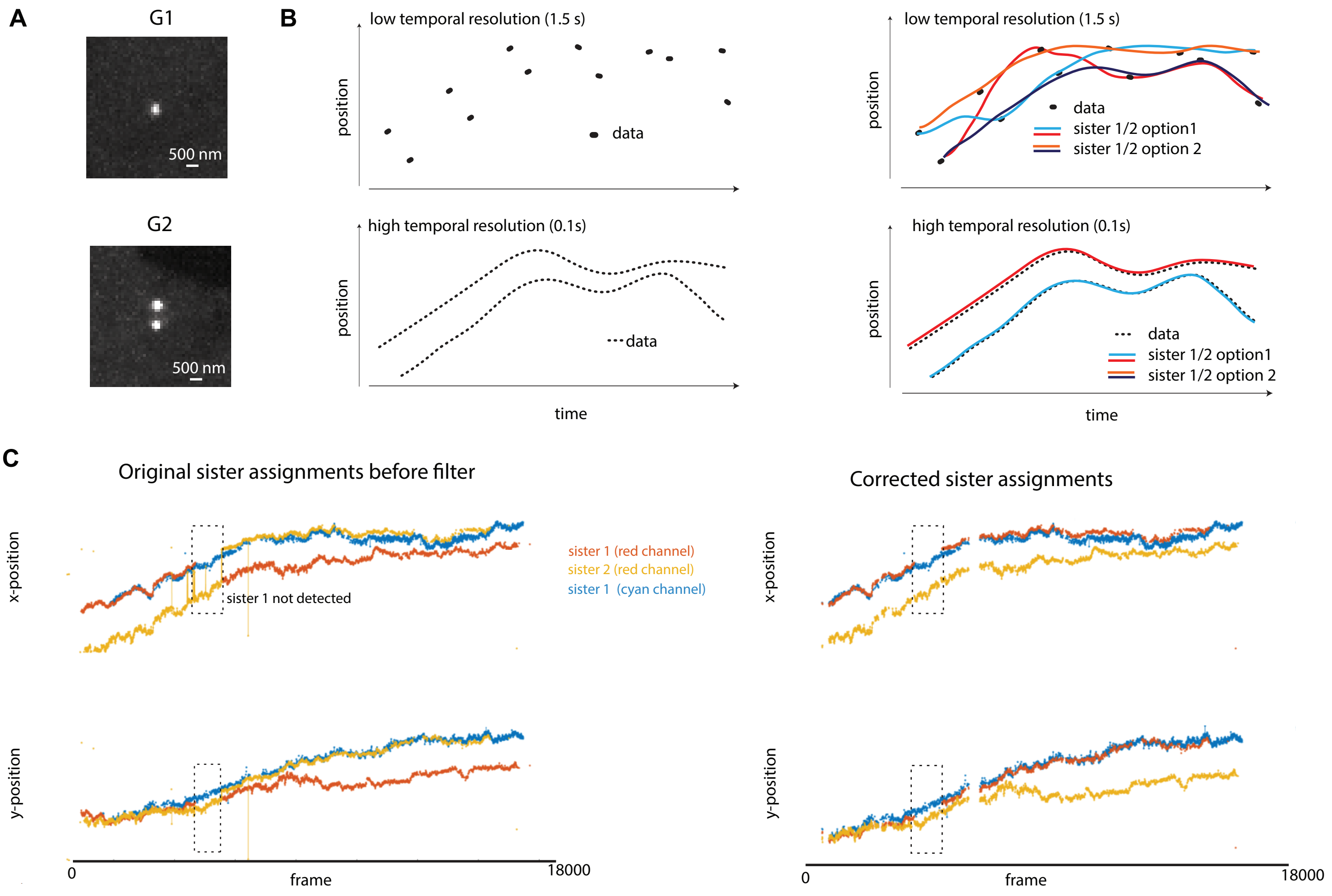

A
G1
500 nm
G2
500 nm
B
low temporal resolution (1.5 s)
position
data
low temporal resolution (1.5 s)
position
data
sister 1/2 option1
sister 1/2 option 2
high temporal resolution (0.1s)
position
data
time
high temporal resolution (0.1s)
position
data
sister 1/2 option1
sister 1/2 option 2
time
C
Original sister assignments before filter
x-position
sister 1 not detected
sister 1 (red channel)
sister 2 (red channel)
sister 1 (cyan channel)
y-position
0
frame
18000
Corrected sister assignments
x-position
y-position
0
frame
18000

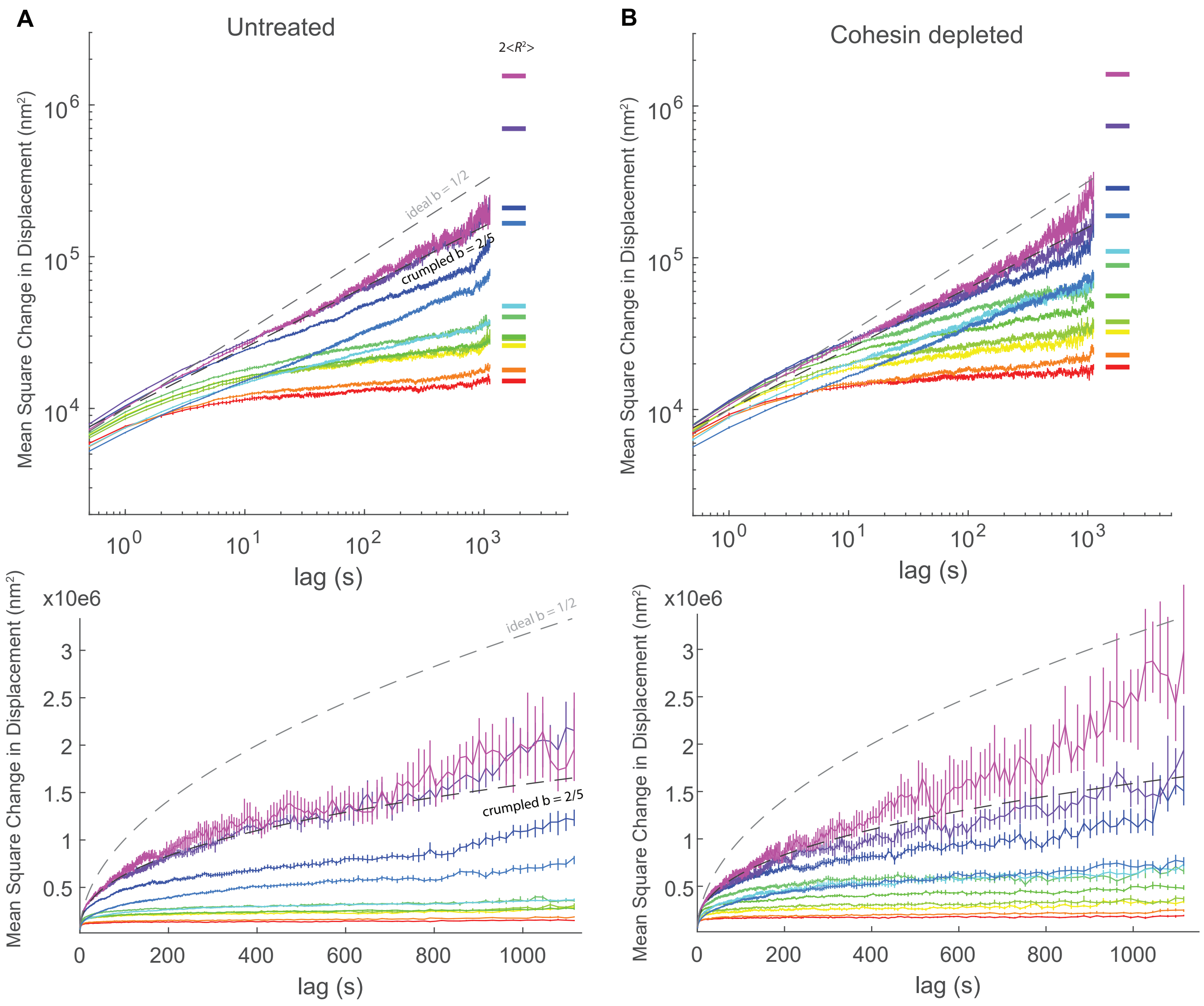
A
Untreated
B
Cohesin depleted
2<R²>
ideal b = 1/2
crumpled b = 2/5
Mean Square Change in Displacement (nm²)
lag (s)
x10e6

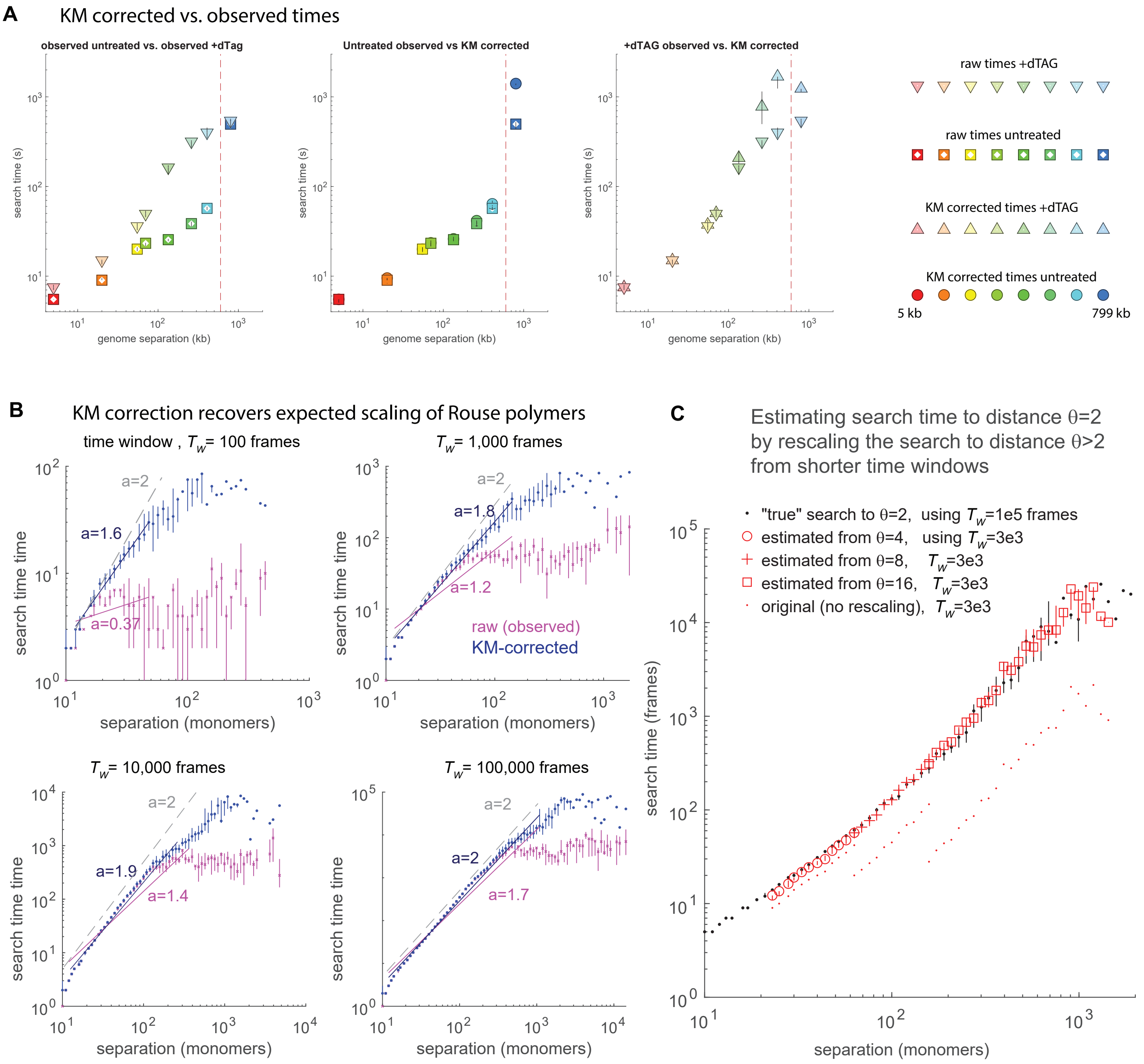
A
KM corrected vs. observed times
observed untreated vs. observed +dTag
Untreated observed vs KM corrected
+dTAG observed vs. KM corrected
search time (s)
genome separation (kb)
raw times +dTAG
raw times untreated
KM corrected times +dTAG
KM corrected times untreated
5 kb
799 kb
B
KM correction recovers expected scaling of Rouse polymers
time window , $T_w$= 100 frames
$T_w$= 1,000 frames
$T_w$= 10,000 frames
$T_w$= 100,000 frames
search time time
separation (monomers)
a=2
a=1.6
a=0.37
a=1.8
a=1.2
a=1.9
a=1.4
a=1.7
raw (observed)
KM-corrected
C
Estimating search time to distance θ=2
by rescaling the search to distance θ>2
from shorter time windows
"true" search to θ=2, using $T_w$=1e5 frames
estimated from θ=4, using $T_w$=3e3
estimated from θ=8, $T_w$=3e3
estimated from θ=16, $T_w$=3e3
original (no rescaling), $T_w$=3e3
search time (frames)
separation (monomers)

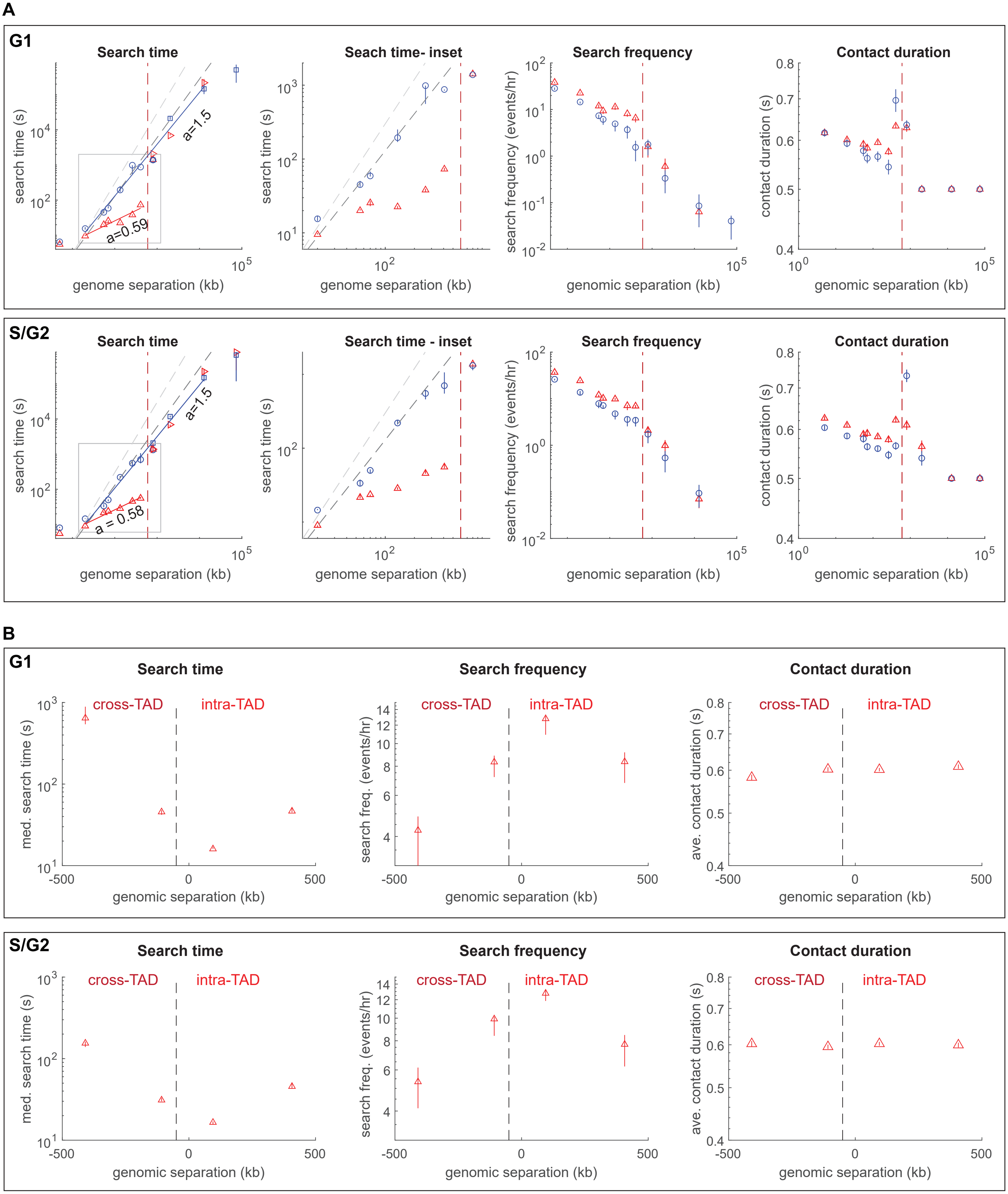
A
G1
Search time
search time (s)
a=1.5
a=0.59
genome separation (kb)
Seach time- inset
search time (s)
genome separation (kb)
Search frequency
search frequency (events/hr)
genomic separation (kb)
Contact duration
contact duration (s)
genomic separation (kb)
S/G2
Search time
search time (s)
a=1.5
a = 0.58
genome separation (kb)
Search time - inset
search time (s)
genome separation (kb)
Search frequency
search frequency (events/hr)
genomic separation (kb)
Contact duration
contact duration (s)
genomic separation (kb)
B
G1
Search time
med. search time (s)
cross-TAD
intra-TAD
genomic separation (kb)
Search frequency
search freq. (events/hr)
cross-TAD
intra-TAD
genomic separation (kb)
Contact duration
ave. contact duration (s)
cross-TAD
intra-TAD
genomic separation (kb)
S/G2
Search time
med. search time (s)
cross-TAD
intra-TAD
genomic separation (kb)
Search frequency
search freq. (events/hr)
cross-TAD
intra-TAD
genomic separation (kb)
Contact duration
ave. contact duration (s)
cross-TAD
intra-TAD
genomic separation (kb)

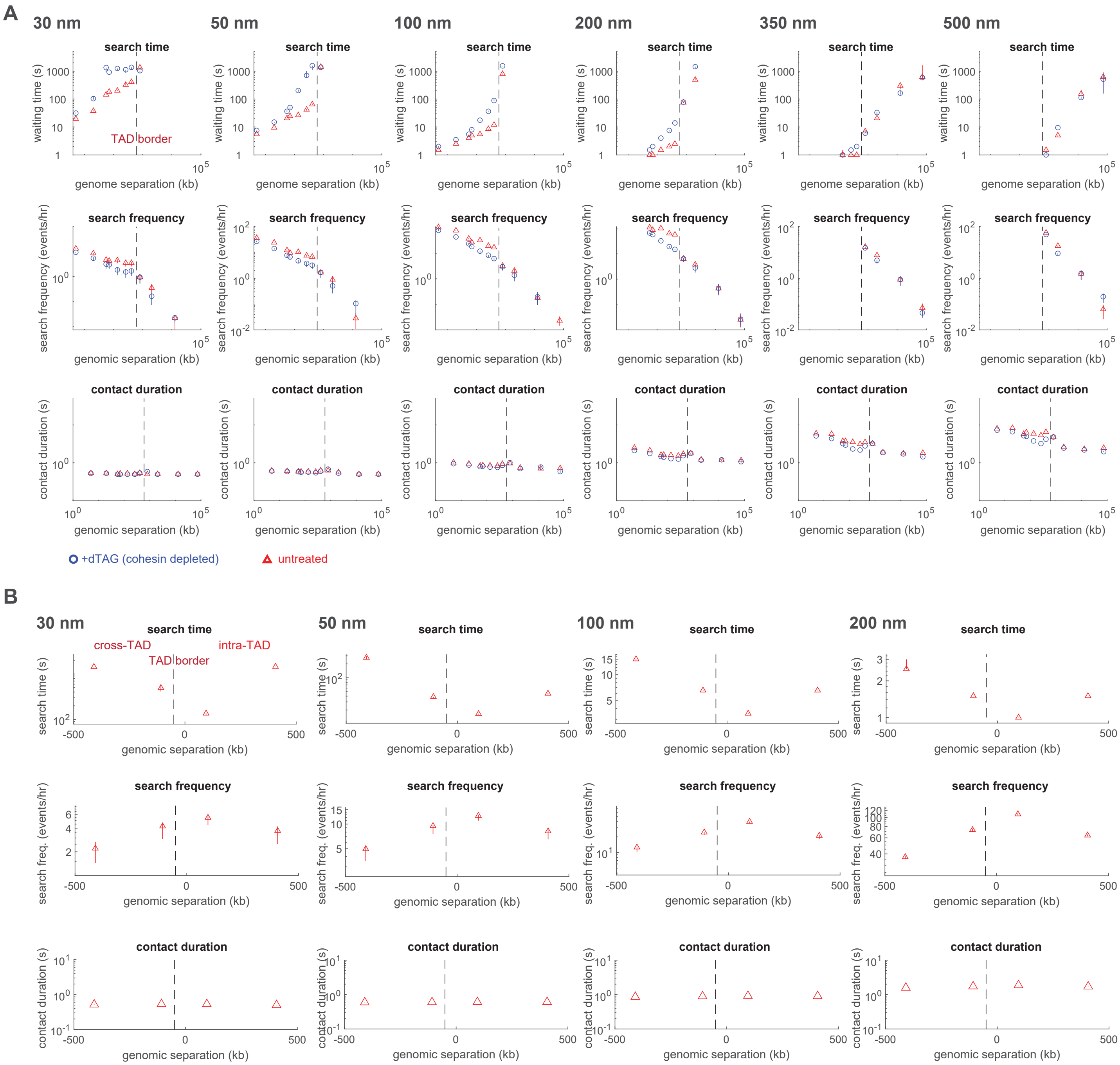
A
30 nm
50 nm
100 nm
200 nm
350 nm
500 nm
search time
waiting time (s)
genome separation (kb)
TAD border
search frequency
search frequency (events/hr)
genomic separation (kb)
contact duration
contact duration (s)
+dTAG (cohesin depleted)
untreated
B
cross-TAD
intra-TAD
search time (s)
search freq. (events/hr)

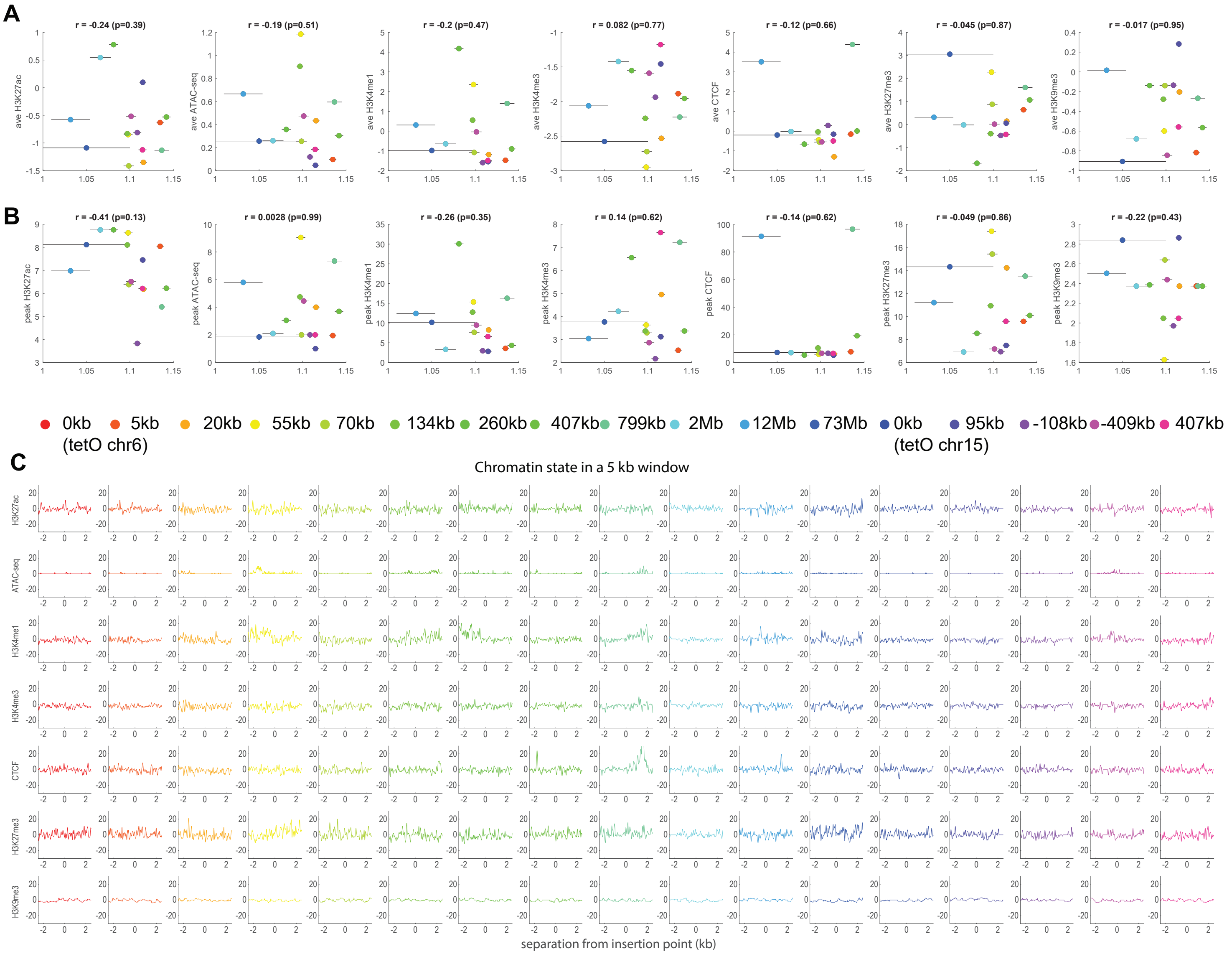

A
r = -0.24 (p=0.39)
r = -0.19 (p=0.51)
r = -0.2 (p=0.47)
r = 0.082 (p=0.77)
r = -0.12 (p=0.66)
r = -0.045 (p=0.87)
r = -0.017 (p=0.95)
ave H3K27ac
ave ATAC-seq
ave H3K4me1
ave H3K4me3
ave CTCF
ave H3K27me3
ave H3K9me3
B
r = -0.41 (p=0.13)
r = 0.0028 (p=0.99)
r = -0.26 (p=0.35)
r = 0.14 (p=0.62)
r = -0.14 (p=0.62)
r = -0.049 (p=0.86)
r = -0.22 (p=0.43)
peak H3K27ac
peak ATAC-seq
peak H3K4me1
peak H3K4me3
peak CTCF
peak H3K27me3
peak H3K9me3
0kb (tetO chr6)
5kb
20kb
55kb
70kb
134kb
260kb
407kb
799kb
2Mb
12Mb
73Mb
0kb (tetO chr15)
95kb
-108kb
-409kb
407kb
C
Chromatin state in a 5 kb window
H3K27ac
ATAC-seq
H3K4me1
H3K4me3
CTCF
H3K27me3
H3K9me3
separation from insertion point (kb)

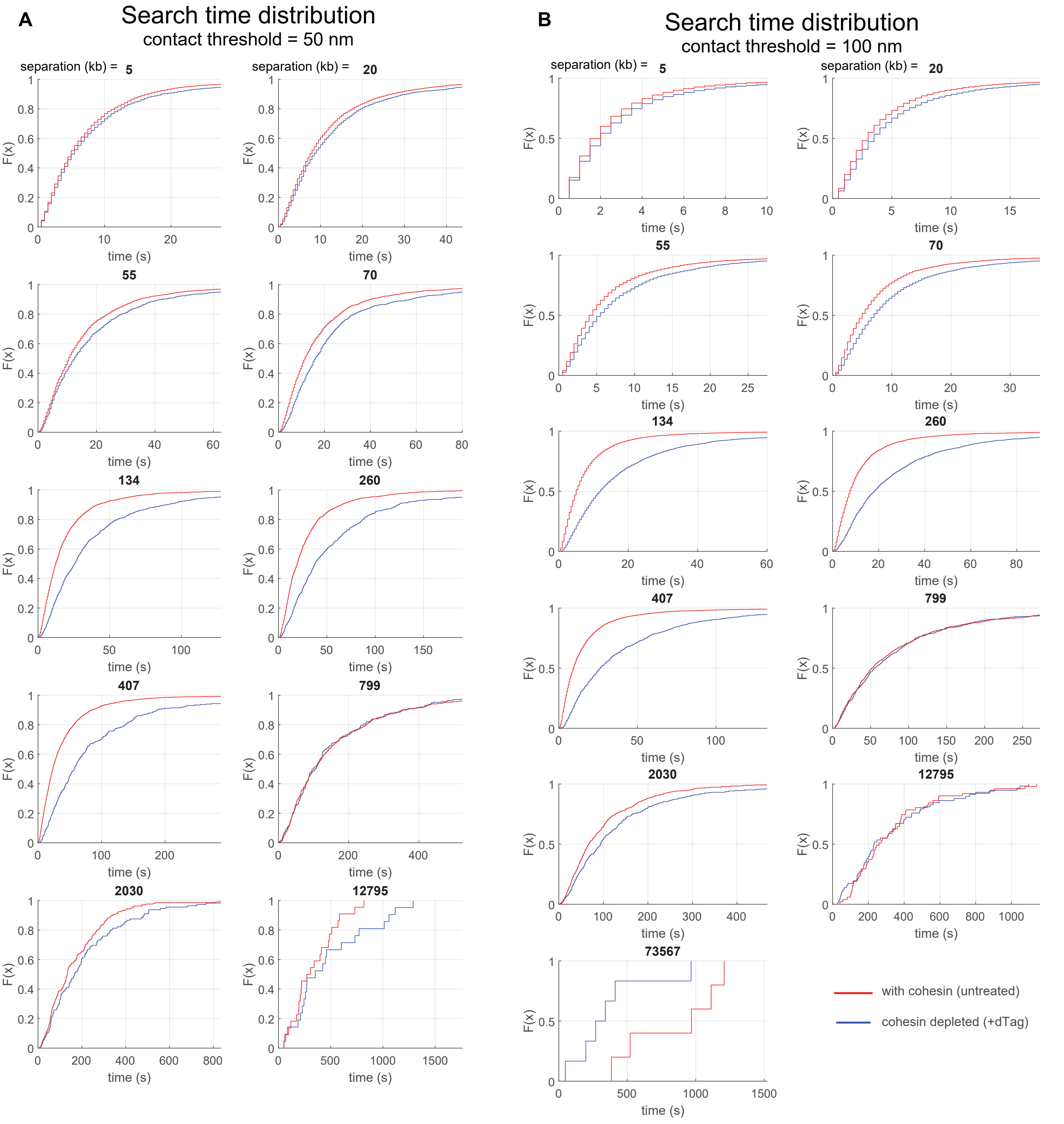
A
Search time distribution
contact threshold = 50 nm
separation (kb) = 5
20
55
70
134
260
407
799
2030
12795
F(x)
time (s)
B
Search time distribution
contact threshold = 100 nm
separation (kb) = 5
20
55
70
134
260
407
799
2030
12795
73567
F(x)
time (s)
with cohesin (untreated)
cohesin depleted (+dTag)
